# Mutations in bacterial regulatory genes are linked with chronic ash tree infections

**DOI:** 10.64898/2026.05.12.723614

**Authors:** K.G. Hinton, D. Vinchira-Villarraga, S. Dhaouadi, G.B. Thomas, M. Rabiey, H.C. McCann, M.C. McDonald, R.W. Jackson

## Abstract

Long term chronic infections of plants by bacterial pathogens are largely unknown. Understanding how pathogens adapt during chronic infection provides a key insight to pathogen evolutionary strategy both for persistence and survival, but also for potential future outbreaks. *Pseudomonas savastanoi pv. fraxini* (*Psf)*, a member of phylogroup 3 within the *Pseudomonas syringae* species complex, causes canker disease in European ash (*Fraxinus excelsior*). Infections persist for years within the bark parenchyma, where bacteria are enclosed in cavities that contribute to the gradual expansion of host periderm. This pathosystem therefore provides an opportunity to examine pathogen evolution in a long-lived, largely unmanaged host. We combined population genomics and phenotypic analysis of 124 *Psf* strains collected from six sites across the UK. Phylogenetic analysis revealed a highly clonal population, with only 833 core genome SNPs across a 5.3 Mb genome, and a relatively small accessory genome largely shaped by gain and loss of large mobile genetic elements. Despite this limited genomic diversity, mutations were enriched in regulatory genes, including two-component systems, chemotaxis proteins, and cell envelope-associated loci. Notably, the global regulator *gacA/S* was independently mutated multiple times within the same clonal lineage. These mutations, typically small deletions, were associated with changes in motility, nutrient utilisation, stress tolerance, and virulence across genetic backgrounds. As a result, phenotypic heterogeneity was observed within otherwise clonal populations, including within individual lesions. These findings indicate that repeated mutation of regulatory systems represents a key mechanism of adaptation in this chronic plant–pathogen interaction, enabling phenotypic diversification despite limited sequence divergence. This study provides a microevolutionary perspective on *P. syringae* populations in the phyllosphere and highlights the role of regulatory variation in the evolution of low-virulence, ecologically restricted pathogens.

## Introduction

*Pseudomonas syringae* is a globally distributed, ecologically diverse species complex occupying multiple environmental and host-associated niches *(Morris et al., 2010).* It comprises at least 13 phylogroups defined by multilocus sequence analysis (Sarkar and Guttman, 2004, Hwang et al., 2005, Berge et al., 2014). These are grouped into primary phylogroups (PGs 1–6 and 10), which are largely plant-associated and encode canonical type III secretion systems, and secondary phylogroups (PGs 7–9 and 11–13), which include atypical species such as *P. viridiflava* and *P. cichorii* and are often isolated from non-agricultural environments (Berge et al., 2014, Dillon et al., 2019a).

While some lineages persist as harmless epiphytes, others are highly virulent pathogenic variants (pathovars) collectively capable of infecting nearly all major crop and woody plant species. Over 60 pathovars have been described, varying widely in their host range and tissue specificity (Berge et al., 2014). Some, such as *P. syringae* pv. *syringae*, exhibit broad host range, infecting more than 200 plant species and capable of infecting multiple tissue types within a single host (Kennelly et al., 2007). Others, including the horse chestnut (*Aesculus hippocastanum*) pathogen *P. syringae* pv. *aesculi*, are highly specialised, as distinct pathovars are restricted to woody tissue (European *Pae*) or leaves (Indian *Pae*) (Green et al., 2010).

*P. syringae* infection pathways can be divided into several key stages: epiphytic survival, entry into internal tissues, suppression of host defences, acquisition of host nutrients for proliferation, and subsequent dispersal (Xin et al., 2018). Many of the genes underlying pathogenicity and host range have been identified across the *P. syringae* species complex, and pathovars differ in their virulence gene content that enable progression through these stages (Baltrus et al., 2011, Nowell et al., 2016, Hulin et al., 2020, Lipps and Samac, 2022). However, gene content alone does not explain ecological success or long-term persistence. For example, many secreted virulence-associated traits, including extracellular enzymes, biosurfactants, toxins, and effectors, generate public goods and that can select for both cooperative and exploitative strategies within pathogen populations (Smith and Schuster, 2019, Ruiz-Bedoya et al., 2023). As a result, disease outcomes are shaped not only by host–microbe–environment interactions, but also by interactions among conspecifics within pathogen populations, especially during co-infection of the same host plant.

Microbial phenotypic diversity arises by multiple mechanisms, including mutation, gene gain and loss, epigenetic modulation, and regulatory change. Recombination and horizontal gene transfer (HGT) are central drivers of *P. syringae* adaptation (Nowell et al., 2014, Dillon et al., 2019a). HGT is often meditated by mobile genetic elements (MGEs) that results in the acquisition of virulence and fitness-associated genes, including T3SS effectors (Jackson et al., 2000, Hulin et al., 2023), copper resistance (Colombi et al., 2017), ultra-violet tolerance (Sundin and Murillo, 1999), and toxin biosynthesis (Watanabe et al., 1998). MGEs also promote within strain genome plasticity through structural rearrangement and inter-replicon transfer (Jackson et al., 1999, Bardaji et al., 2011). *De novo* mutations also contribute to host adaptation, including abolition of effector triggered cell death (e.g. deletions in *hopM1*) and altered host immune recognition (e.g. single nucleotide polymorphisms in *fliC*) (Cai et al., 2011, Clarke et al., 2013). Mutations in global regulators such as GacA/S can also drive rapid phenotypic shifts, altering exoenzyme production, secondary metabolism, and motility, although this has not been reported in phyllosphere pathogens (van den Broek et al., 2005, Lalaouna et al., 2012, Achouak et al., 2007). However, adaptive variants are not necessarily fixed within pathovars, and substantial diversity can be maintained at fine spatial and ecological scales through dynamics like resource partitioning and regulatory switching (Rufian et al., 2016, Rufian et al., 2018, Lopez-Pagan et al., 2025).

The extent and structure of this population-level diversity in natural *P. syringae* populations remains poorly characterised, despite evidence that pathogen diversity within a single host can strongly influence disease severity and transmission (Susi et al., 2015). Contrasts between agricultural and wild systems highlight the role of ecological context in shaping *P. syringae* population structure. Epidemic outbreaks in crops are frequently driven by the rapid expansion of clonal lineages, such as *P. syringae* pv. *tomato* T1 and *P. syringae* pv. *actinidiae* biovar 3 (*Psa-3*) (Cai et al., 2011, McCann et al., 2017). In contrast, wild *Arabidopsis thaliana* populations harbour diverse *Pseudomonas* communities, but are consistently dominated by a single pathogenic lineage, OTU5, which comprises numerous deeply divergent strains most closely related to *P. syringae* and *P. viridiflava* (Karasov et al., 2018). These patterns suggest that natural and managed ecosystems favour distinct evolutionary trajectories, with agriculture promoting clonal amplification while wild systems maintain diversity, although findings are likely influenced by sampling strategies (Zeng et al., 2026). Nonetheless, wild populations may act as reservoirs from which epidemic clones can emerge, as demonstrated by the origin of pandemic *Psa-3* lineages from a recombining source population prior to global dissemination via plant trade (McCann et al., 2017).

*Pseudomonas savastanoi* belongs to phylogroup 3 of the *P. syringae* species complex, including five woody-infecting pathovars which are generally recovered from a single host species: pv. *savastanoi* (olive), pv. *retacarpa* (Spanish broom), pv. *nerii* (oleander), pv. *mandevillae* (Dipladenia), and pv. *fraxini* (ash) (Gardan et al., 1992, Moreno-Perez et al., 2020, Caballo-Ponce et al., 2021). Cross-infection assays demonstrate that these pathovars are not strictly host-specific and display overlapping host ranges. For example, all pathovars cause symptoms on ash and olive (both *Oleaceae* family), as well as their main host of isolation (Ramos et al., 2012, Caballo-Ponce et al., 2021, Moreno-Perez et al., 2020). While these studies reported knot formation for most pathovars, *P. savastanoi pv. fraxini (Psf)* induces a distinctive “erumpent canker” in olive and ash.

*Psf* causes natural infections in European ash *(Fraxinus excelsior)* across the UK and Europe (Boa, 1978, Janse, 1981b, Janse, 1981a, Brown, 1932). Infection is initiated through wounds in woody tissue and proliferates intercellularly within the bark parenchyma, triggering host physiological changes that lead to callus and periderm formation around expanding, bacteria-filled cavities (Janse, 1982). *Psf* overwinters within bark tissue and renewed bacterial proliferation each growing season triggers periderm formation resulting in slowly enlarging cankers, which multiply due to secondary spread within a single host. This distinctive symptomology is attributed to negligible production of auxin (IAA) and cytokinin, which drive knot formation in other *P. savastanoi* pathovars (Aragon et al., 2014, Caballo-Ponce et al., 2017, Surico et al., 1985). Comparative genomics indicates that *Psf* strains lost several genomic regions carrying virulence-associated genes, possibly prior to dissemination in ash populations. Despite a reduced accessory genome, they retain an expanded effector repertoire (Moreno-Perez et al., 2020). This pattern is consistent with recent clonal expansion or compensatory evolution during niche specialisation.

*Psf* provides a unique opportunity to examine how pathogenic populations diversify within a long-lived, unmanaged host. European ash persists for decades and regenerates naturally, allowing host–pathogen interactions to unfold over extended timescales. *Psf* establishes chronic infections within woody tissues, where populations can persist within cankers for years. This system therefore raises key questions about how prolonged residence within a single host shapes pathogen evolution across the infection lifecycle. We hypothesised that long-term infection would favour small effective population sizes, minimal HGT, and diversification driven primarily by mutation, selection, and within-host competition. To test this, we conducted the first population genomic and phenotypic analysis of *Psf*, sampling isolates from six woodland sites across the UK. We show that *Psf* forms highly clonal, geographically structured populations with exceptionally low evolutionary rates, yet exhibits ongoing genotypic and phenotypic diversification, often involving regulatory mutations, with divergent genotypes co-existing within individual cankers.

## Results

### *Psf* populations are highly clonal and geographically structured across the UK

Ash (*F. excelsior*) is a dominant tree species in the UK landscape and is exposed to a range of biotic stresses, including infection by the bacterial pathogen *Psf*. *Psf* causes erumpent cankers on ash stems and branches, and these symptoms are now widely distributed across the UK. Between January 2022 and May 2023, 166 *Psf* isolates were recovered from symptomatic and non-symptomatic ash bark from 22 trees across the six UK sites (**Figure S1**): Wytham Woods (9 trees, n = 86), Matlock (2 trees, n = 6), Lathkill Dale (6 trees, n = 24), Marden Park Wood (1 tree, n = 1), Isle of Mull (1 tree, n = 25), and Cressbrook Dale (3 trees, n = 24). *Psf* was primarily isolated from symptomatic trees, although 14 isolates were recovered from healthy trees. Several canker types were observed, including erumpent, shot-hole and letterbox cankers, vertical cracks, and a rare bleeding lesion (**Figure S2**).

Whole-genome sequences from 122 strains collected in this study, together with the historical isolates NCPPB 1006 and CFPB 5062, were used to assess population structure (**Table S1**). Complete genome assemblies were generated for strains NCPPB 1006, W163a3b1, and L1928a1b6, with W163a3b1 used as the reference for phylogenetic analysis. Following removal of plasmid sequences, masking of mobile genetic elements (MGEs), and exclusion of recombinant regions, the resulting gap-free core chromosome comprised 5.27 Mb. A total of 351,732 bp were masked, thereby minimising the influence of horizontally acquired variation on phylogenetic inference (**Table S3**). Across this alignment, 833 non-recombinant, non-MGE single nucleotide polymorphisms (SNPs) were identified, indicating low overall sequence diversity.

The resulting SNP phylogeny exhibited a highly clonal population structure with limited divergence among strains, but 10 well-resolved clusters (**Figure 1a**). Terminal branches were short, with mean intra-clade pairwise distances of 13.2 SNPs (s.d. 11.3), whereas divergence between clades was an order of magnitude greater (mean 130 SNPs, s.d. 51.1; **Figure S3**). Clusters broadly corresponded to geographic origin. For instance, most strains from the geographically proximate sites of Lathkill Dale, Matlock, and Cressbrook Dale clustered closely together, consistent with likely recent shared ancestry and/or ongoing local connectivity. In contrast, a deeply divergent clade was also recovered from Lathkill Dale, indicating that multiple *Psf* lineages circulate within local ash populations and reflect both local dispersal and long-term lineage persistence. This pattern was supported by a Mantel test comparing pairwise genetic and geographic distances, which revealed a moderate positive correlation (r = 0.64, p < 0.001; **Figure S4**). Notably, the greatest range of genetic variation occurred at very small spatial scales (0–190 SNPs at ∼0 km), demonstrating that both clonal and highly divergent lineages can coexist locally. In contrast, genetic distances plateaued between 100 and 700 km, suggesting increased geographical distance does not necessarily reflect increased genetic divergence.

**Figure 1.**
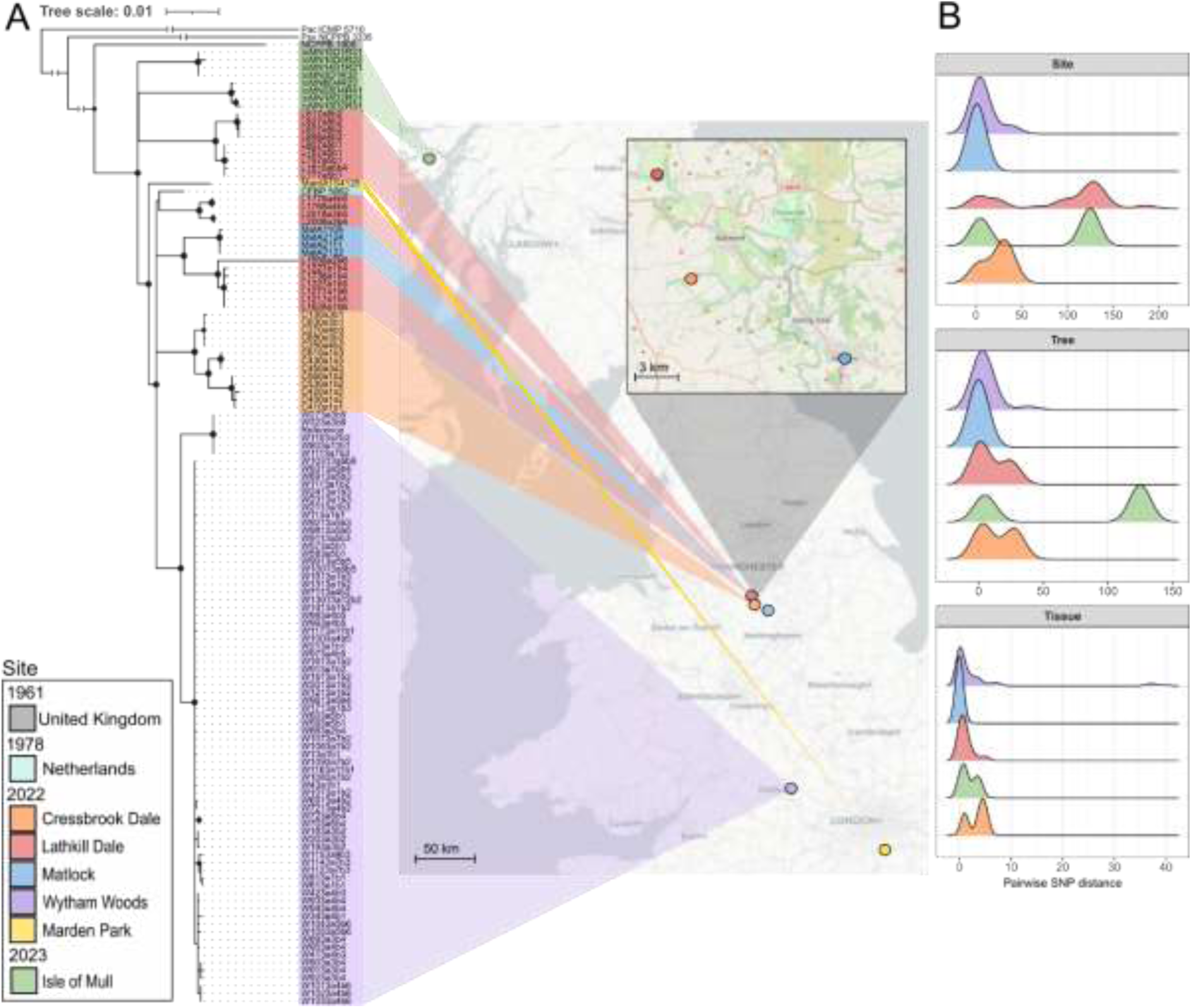
Core genome diversity of *Psf*. (A) Biogeographical distribution and core genome SNP phylogeny of 124 *Psf* strains isolated from *F. excelsior* bark tissue. A maximum-likelihood phylogeny was inferred using RAxML-NG from a recombination- and MGE-masked core SNP alignment and rooted with *P. amygdali* pv. *ciccaronei* ICMP 5710. All nodes have bootstrap support ≥70 (based on 1,000 Felsenstein bootstrap replicates); nodes with support ≥90 are indicated by black circles. Strains are colour-coded by geographic origin and mapped on the right: Isle of Mull (green), Lathkill Dale (red), Cressbrook Dale (orange), Matlock (blue), Wytham Woods (purple), and Marden Park Wood (yellow). Historical isolates CFBP 5062 (light blue; Netherlands, 1978), NCPPB 1006 (grey; UK, 1961), *P. savastanoi* pv. *savastanoi* NCPPB 3335 (France), and *P. amygdali* pv. *ciccaronei* ICMP 5170 (Italy), are included in the phylogeny but are not shown on the map. The branches for the latter three strains are truncated for tree visualisation. (B) Pairwise SNP distance distributions among strains at three nested sampling scales: site, tree, and tissue.

To further determine how this population structure is maintained across ecological scales, we quantified genetic variation at the site, tree, and tissue levels. Patterns of genetic structure varied markedly among sites (**Figure 1b; Table S4**). Lathkill Dale exhibited the greatest diversity, resolving into three distinct clades with a mean pairwise distance of 90.2 SNPs, while Isle of Mull contained two divergent clades (mean 73.6 SNPs). In contrast, Cressbrook Dale and Wytham Woods were dominated by one or two closely related clades (mean 23.1 and 8.4 SNPs, respectively), and Matlock was nearly clonal, with isolates differing by a single SNP. This suggests clonal expansion and localised dispersal of a single founding lineage.

At the tree scale, most hosts were dominated by a single genetic lineage. Limited diversification within hosts was observed at Cressbrook Dale and Lathkill Dale, where isolates from individual trees differed by a mean of 14.5 and 14.2 SNPs, respectively. In contrast, both major Isle of Mull clades were recovered from a single host, differing by 73.6 SNPs, indicating host colonisation by multiple divergent lineages.

At the tissue scale, lesion populations were predominantly clonal (**Figure 1b**). Of the 30 tissue samples with multiple isolates, six showed no detectable genetic variation, while the remainder differed by a mean of 3.7 ± 6.8 SNPs. However, three tissue samples from asymptomatic trees at Wytham Woods contained multiple genetic clades, with pairwise distances ranging from 15 to 27 SNPs, demonstrating that divergent lineages can coexist within individual tissues in the absence of visible disease symptoms.

Across all tissues, 95 within-tissue alleles were identified **(Figure S5**). Of these, 52 were unique, consistent with recent, local diversification. Forty alleles were shared among multiple trees within Wytham Woods, indicating the circulation of two closely related clonal populations across hosts. Three alleles were shared across multiple sites. These included a missense mutation in *aceA* (isocitrate lyase; *PPLAPI_16080*) in Lathkill Dale and Wytham Woods; an intergenic SNP between *hrpA* and *fadK* (*PPLAPI_21020* and *PPLAPI_21025)* in Cressbrook Dale and the Isle of Mull; and a missense mutation in the *hscC* hsp70 molecular chaperone (*PPLAPI_21900*) in Cressbrook Dale, Lathkill Dale, and Wytham Woods.

To assess whether mutations accumulated non-randomly across the genome, we examined their functional distribution. SNPs were only slightly depleted from coding regions relative to expectation (85.4% observed vs 88% expected; χ² = 5.54, *P* = 0.019), indicating weak purifying selection. However, mutations were unevenly distributed across loci, with 18% occurring in genes that carried multiple independent mutations (two to four per gene; **Table S5**). Two genes accumulated four mutations each, including a periplasmic oligopeptide-binding (PPLAPI_03825) and the global regulator *crp* (PPLAPI_17395), while four additional loci, the DNA polymerase I *polA* (PPLAPI_01185), *rpoB* (PPLAPI_23820), glycine-zipper protein (PPLAPI_12590), and a calcium-binding haemolysin (PPLAPI_05330), carried three mutations. Furthermore, several genes with recurrent mutation have established roles in virulence or host interaction, including methyl-accepting chemotaxis proteins (*tar*; PPLAPI_02645 and *ctpH;* PPLAPI_13005), *paaJ* (PPLAPI_05510), the T3SS effector *avrE1* (PPLAPI_07690), and the transcriptional regulator *lysR* (PPLAPI_13580), possibly indicating selection acting on host-associated functions.

### Genome stability and limited horizontal gene transfer

Despite high core genome clonality, variation in gene content was evident at the accessory genome level. The pangenome of *Psf* strains, including four additional publicly available genomes (n = 128), comprised 6,424 genes, of which 82% (n = 5,289) formed the soft-core genome, with 506 shell and 629 cloud genes (**Figures 2a–b**). Clustering strains based on the presence or absence of 1,459 variable genes revealed groups of isolates with divergent accessory gene content. These groups broadly corresponded to phylogenetic clades identified through concatenated core gene alignment, although clustering did not consistently mirror phylogeny. Notably, isolates from Wytham Woods and the Isle of Mull clustered together despite clear core gene divergence.

**Figure 2.**
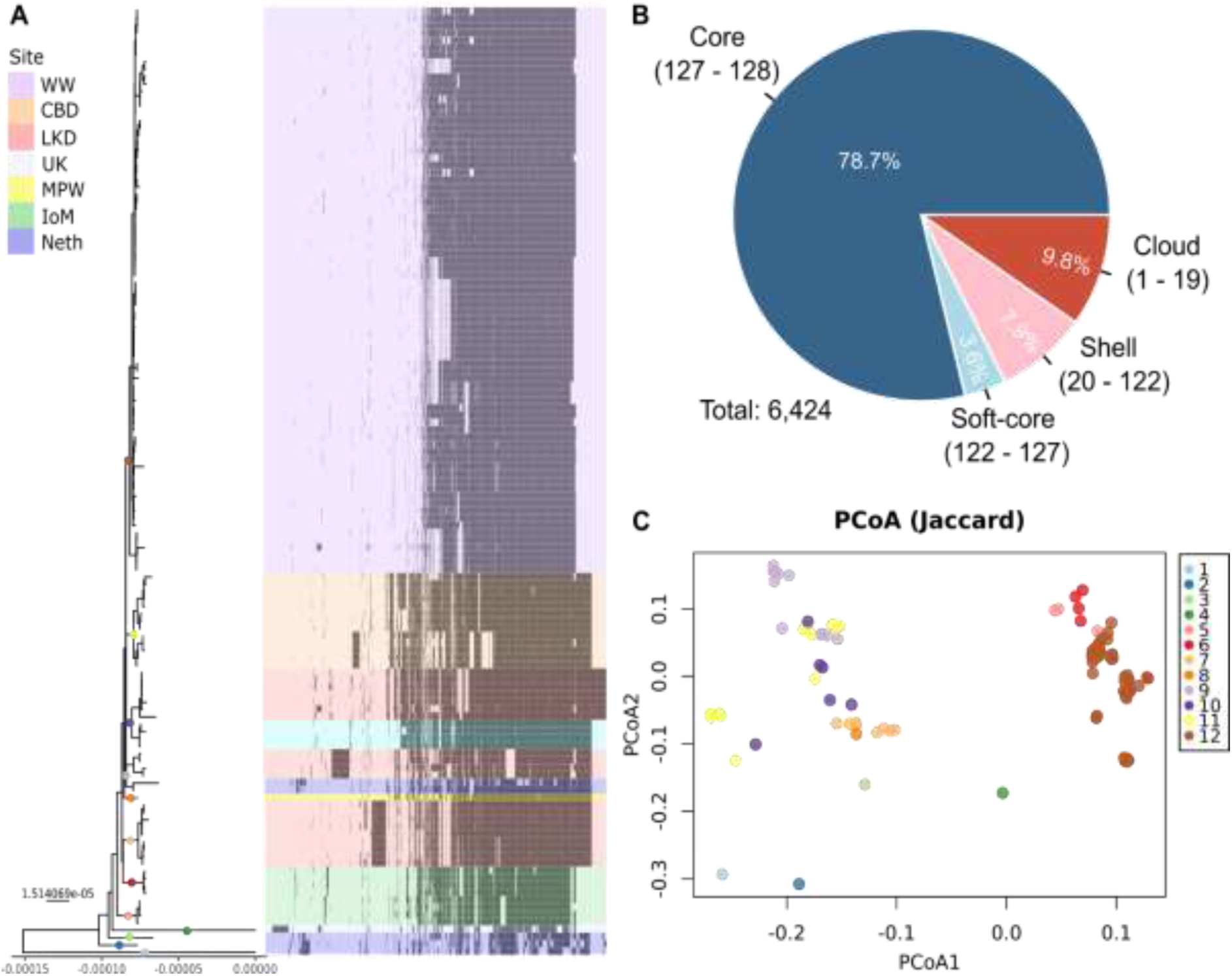
Pangenome diversity of *Psf*. (A) The presence/absence of variable genes (n = 1,459) across 128 *Psf* genomes, including 122 isolates collected in this study, plus NCPPB 1006, CFPB 5062, ICMP7711, ICMP7712, ICMP9129, ICMP9132 (Dillon *et al.,* 2019b). Strains are coloured by site: Wytham Woods (purple), Cressbrook Dale (orange), Lathkill Dale (red), Matlock (blue), Netherlands (dark blue), Marden Park (yellow), Isle of Mull (green), and UK (light grey). Clades used in the PCoA are denoted by coloured circles on the clade branches. The phylogenetic tree was generated from a concatenated alignment of core gene sequences (>99% of isolates) using IQ-TREE2 and midpoint-rooted for visualisation. (B) Pangenome composition, showing the proportion of genes classified as core (99–100%), soft-core (95–98%), shell (15–94%), or cloud (0–14%). (C) Clustering of strains based on accessory genome composition (genes present in <100% strains). Principal coordinate analysis (PCoA) of gene presence/absence is coloured by clade (1–12) which are also annotated on the branches of the tree.

To identify the genetic basis of accessory genome clustering, distance-based redundancy analysis (dbRDA) was used to resolve genes most associated with variation. The top 100 genes were overwhelmingly associated with DNA mobility, including recombination, conjugation, and transposition (n = 34), alongside a large proportion of poorly characterised proteins (14 hypothetical and 10 of unknown function; **Figure S6**). Other functional categories included DNA processing, phage-related functions, stability systems, virulence, DNA repair, and transport.

Variation along the first dbRDA axis was driven by a set of genes consistently absent from isolates sampled from Wytham Woods, the Isle of Mull, and the reference strain NCPPB 1006. As these genes were also absent from the reference genome *Psf* W163, we examined the complete genome of strain *Psf* L1928, which contained the full set of genes. This revealed that the genes are located within a ∼121 kb integrative and conjugative element (ICE), inserted at a tRNA-Lys locus and flanked by site-specific integrases. The ICE encodes multiple accessory functions, including metabolic genes (e.g. a sulfatase-like hydrolase/transferase and *cysC*) and both a pseudogenised and intact type III secretion system effector (*hopZ4*) (**Figures S7 and S8**).

The second dbRDA axis captured variation associated with a distinct set of mobile elements. Genes with negative loadings were largely absent from clade 7 and comprised predominantly phage-associated and regulatory functions. Manual inspection identified these as components of a 47 kb lysogenic phage present exclusively in clade 7 isolates, containing virulence-associated genes including CsrA transcriptional regulator, a T6SS effector, peptidoglycan hydrolase and chitinase (**Figure S9**). In contrast, genes with positive loadings were absent from clades 1–4, 7, and 8, and were enriched for plasmid-associated functions, including toxin–antitoxin systems, the surface exclusion protein ExcA, and transporters, which may suggest the presence or loss of a plasmid (**Figure S6**). Collectively, these results indicate that accessory genome structure is shaped by a small number of large MGEs, rather than widespread gene gain and loss across the pangenome.

### Mutational patterns reveal repeated targeting of environmental sensing and global regulatory pathways

We further assessed the functional distribution of *de novo* mutations by comparison of GO and KEGG terms for genes encoding *de novo* mutations, including SNPs, multiple nucleotide polymorphisms, insertions, and deletions (2,644), against all genes within the reference genome used for variant calling. Of the 6,052 annotated reference genes, 767 contained non-synonymous variants, 340 contained synonymous variants, and 114 harboured both. Functional grouping using KEGG (37% CDS annotated) and gene ontology (GO) (28% CDS annotated) categories enabled analysis of biological pathways enriched for genetic variation.

While no GO pathways were significantly enriched in mutations (**Figure S10**), KEGG pathway enrichment identified two pathways with significant overrepresentation after multiple-testing correction (FDR; q < 0.01): two-component systems (TCS) (68 genes) and beta-lactam resistance (17 genes), although nine genes were present across both pathways (**Figure 3a; Table S6**). The beta-lactam–associated genes encode proteins localised to the cell envelope (periplasmic, inner membrane or outer membrane), are implicated in cell-wall biogenesis, envelope integrity, compound sensing and export, or envelope stress responses.

**Figure 3.**
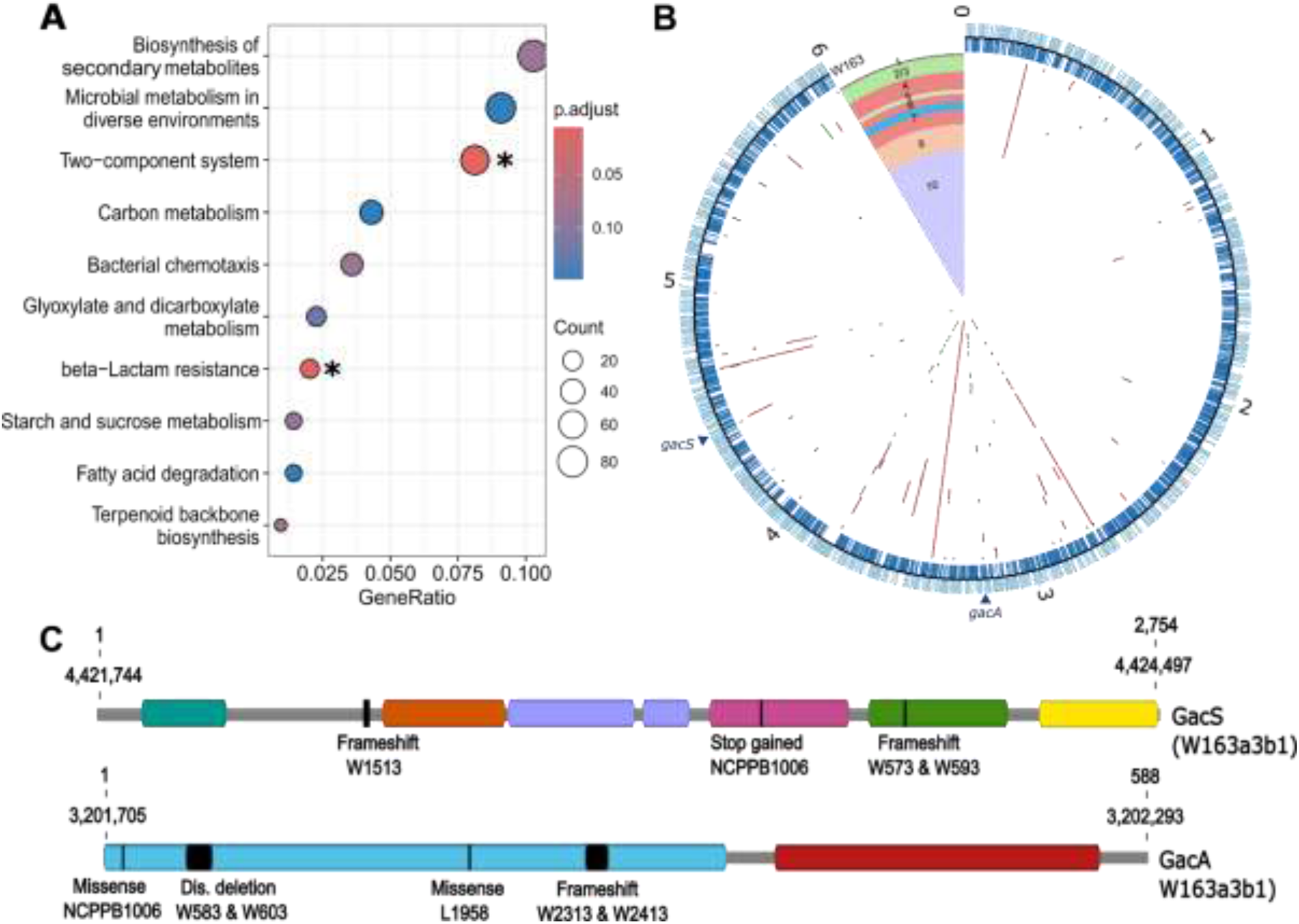
Ecologically significant pathways under selection. (A) KEGG pathways significantly enriched for mutations (SNPs, multiple nucleotide polymorphisms (MNPs), indels, and complex variants) across 124 *Psf* genomes. Asterisks indicate significant enrichment (Fisher’s exact test, p < 0.05; Benjamini–Hochberg FDR, q < 0.1). (B) Two-component system (TCS) mutations were mapped onto the W163 reference genome using CIRCOS (Krzywinski et al., 2009). Forward and reverse CDS are shown in the outer rings (blue), with genome coordinates marked at 1 Mb intervals (1–6 Mb). Inner rings display TCS mutations from 124 *Psf* genomes, ordered according to the phylogeny. Mutation types are coloured as follows: SNPs (red), complex variants (blue), insertions (purple), deletions (green), and MNPs (orange). The triangular section indicates strain isolation site: Isle of Mull (green), Lathkill Dale (red), Cressbrook Dale (orange), Matlock (blue), Wytham Woods (purple), Marden Park Wood (yellow), and historical isolates CFBP 5062 (light blue; Netherlands, 1978) and NCPPB 1006 (grey; UK, 1961). Numbers within coloured blocks denote phylogenetic groups. (C) Distribution of mutations in GacA and GacS proteins in strain W163. GacS (1-2,754bp, 917aa) contains six predicted domains (PAS, teal; helix hairpin, orange; histidine kinase ATPase, purple; two Rossmann/HAD-like domains, pink & green; histidine kinase sensor, yellow), with mutations occurring before domain 2 and within domains 4 and 5. GacA (1-588bp, 195aa) comprises a response regulator (blue) and a winged helix-like DNA-binding domain (red); all mutations occur within the response regulator domain. Domains were predicted using AlphaFold (Qscore > 80, data not shown).

TCS’s encompass signal transduction and environmental sensing pathways. Among the 88 unique TCS variants identified, 41 were strain-specific, including 20 specific to NCPPB 1006, and the remainder occurred in an average of 8 strains (±12 S.D.) (**Figure 3b**). Repeated independent mutations occurred in global regulators controlling bacterial lifestyle, virulence, stress tolerance, and community behaviour, including GacA/S (n = 7), CRP-cAMP global regulator (n = 4), EnvZ/OmpR (n = 1), AtoS/AtoC (n = 3), PleC/PleD (n = 1), PilS/PilR (n = 1) and PilS/PilG (n = 1) (**Table S6**). Nutrient-responsive TCS and chemotaxis also showed repeated variation (10 and 28 mutations, respectively), particularly in *dctB*-like histidine kinases and methyl-accepting chemotaxis proteins, suggesting alteration of environmental sensing and motility. Mutations were additionally observed in envelope stress and efflux genes (e.g. TolC) (8), central metabolism (8), transporters (3), respiration (1), replication (1), and unclassified signalling domains (8), reflecting widespread rewiring of core physiological and regulatory networks. Together, these patterns indicate adaptation of *Psf* in the ash niche through mutation of global regulators, nutrient sensing, and motility pathways.

### *gacA*/*S* Variants Introduce Phenotypic Heterogeneity within Lesions

The GacA/S TCS is a global regulator which controls diverse virulence-associated pathways. Of the seven independent *gacA*/*S* mutations identified through enrichment analysis, five were predicted to have high-impact effects on gene function (stop codon-gain, frameshift (fs), or disruptive deletion) and two had likely moderate effects (missense) (**Figure 3c**). Mapping the five strains with high-impact *gacA*/*S* mutations back to their origin of isolation revealed that two lesions from Wytham Woods ash tree 1 harboured mixed populations: *gac*A^Q91fs^ comprised 33% of isolates from lesion WA1B2, while *gacS^Y700fs^* comprised 12.5% of isolates from lesion WA1B3. Furthermore, a lesion from Wytham Woods ash tree 5 contained both *gacA^S16S20del^* and *gacS^I232fs^* mutants, with no reference (W163 alleles) *gacA*/*S* detected (**Table S7**). These results suggest divergent within-lesion population dynamics of recurrent *gacA*/*S* mutants.

We would predict that the high-impact mutations would have corresponding effects on bacterial behaviour and fitness. Based on previous studies, these could be related to motility, oxidative stress, T3SS, metabolism, and virulence (Lavado-Benito et al., 2024). We first focused on motility and observed a non-motile phenotype in all naturally occurring high-impact *gacA*/*S* mutants, while the reference strain W163 was highly motile (**Figure S11**). We expected that providing reference *gacA*/*S* alleles would restore motility to non-motile strains and conversely introducing mutant alleles to reference *gacA*/*S* would abolish motility. We complemented two natural *gacA* and *gacS* mutants, *gac*A^Q91fs^ in strain W2313 and *gacS^Y700fs^* in strain W1513, with reference *gacA* and *gacS* from strain W163, respectively. After complementation, both strains exhibited a change from being non-motile to motile in swarming agar assays (**Figure S11**). Moreover, we used an allele swap approach to introduce the W2313 *gac*A^Q91fs^ and W1513 *gacS^Y700fs^* alleles into motile W163. In both instances motility was abolished (**Figure S12**). Together these demonstrate that genotypic changes are occurring in natural populations that have consequences for bacterial behaviour.

We then expanded the phenotypic testing to investigate a broader range of behavioural changes, including swimming, biosurfactant production, antibiotic susceptibility, stress resistance, and carbon source utilisation. To assess the phenotypic impact of *gacA*/*S* mutations across diverse genetic backgrounds, we included both naturally occurring mutants and engineered mutants, alongside strains naturally carrying the reference alleles from the same genetic background or tissue. In total, nine strains were phenotyped, including: W2013 and W2313 *gac*A^Q91fs^ (ash 1 tissue 2), W1413 and W1513 *gacS^Y700fs^*(ash 1 tissue 3), W573 *gacS^I232fs^* and W593 *gacA^S16S20del^* (ash 5 tissue 1), and finally W163 (tree 3 tissue 1) and its engineered mutant derivatives W163 *gac*A^Q91fs^ and W163 *gacS^Y700fs^* (**Table 1**).

**Table 1.** Nomenclature of strains used in text. Strains include *gacA*/*S* mutants, non-mutant strains from the same tissue, and engineered *gacA*/*S* mutants.

| Tissue | Strains | Mutated gene name | Locus tag | DNA sequence change | Amino acid sequence change (three-letter code) | Standard nomenclature (used in text) |
| --- | --- | --- | --- | --- | --- | --- |
| Wytham Woods Ash 1 Tissue 2 | W2313 WT | *gacA* | 2898 | c.272_284del | p.Glu91fs | W2313 *gacA*<sup>Q91fs</sup> |
|  | W2013 WT | Non-mutant GacA/S (100 % homology to W163 WT) |  |  |  | W2013 |
| Wytham Woods Ash 1 Tissue 3 | W1513 WT | *gacS* | 4037 | c.2098delT | p.Tyr700fs | W1513 *gacS*<sup>Y700fs</sup> |
|  | W1413 WT | Non-mutant GacA (100 % homology to W163 WT) |  |  |  | W1413 |
| Wytham Woods Ash 5 Tissue 2 | W593a5b1 WT | *gacA* | 2898 | c.47_61del | p.Ser16_Ser20del | W593 *gacA*<sup>S16S20del</sup> |
|  | W573a5b1 WT | *gacS* | 4037 | c.693_705del | p.Ile232fs | W573 *gacS*<sup>I232fs</sup> |
| Wytham Woods Ash 1 Tissue 1 | W163 WT | Functional GacA/S (Reference alleles) |  |  |  | W163 WT |
|  | W163 | *gacA* | 2898 | c.272_284del | p.Glu91fs | W163 *gacA*<sup>Q91fs</sup> |
|  | W163 | *gacS* | 4037 | c.2098delT | p.Tyr700fs | W163 *gacS*<sup>Y700fs</sup> |

We used a linear mixed-effects model and post hoc comparison to test the effects of treatment (media supplement), *gacA*/*S* group (mutant or reference), and their interaction on bacterial growth after 72 hours, with experiment included as a random effect. Strains with reference *gacA*/*S* exhibited condition-dependent growth responses in minimal medium supplemented with different carbon sources (**Figure 4a**). Specifically, strains W2013, W1413, and W163, showed significantly reduced growth in glycerol (*p* < 0.05), glucose alone, and glucose supplemented with valine or serine (all *p* < 0.0001), but significantly increased growth in glucose supplemented with glutamate (*p* < 0.0001). Post hoc comparisons between strains with reference *gacA*/*S* revealed significantly reduced growth in glucose and valine (*p* < 0.005) and increased growth in glucose and glutamate (*p* < 0.0001) relative to glucose alone (**Table S8**). In contrast, mutant strains showed no significant variation across these treatments relative to glucose alone (all *p* > 0.05), indicating a loss of nutrient-responsive growth change (**Table S9**).

**Figure 4.**
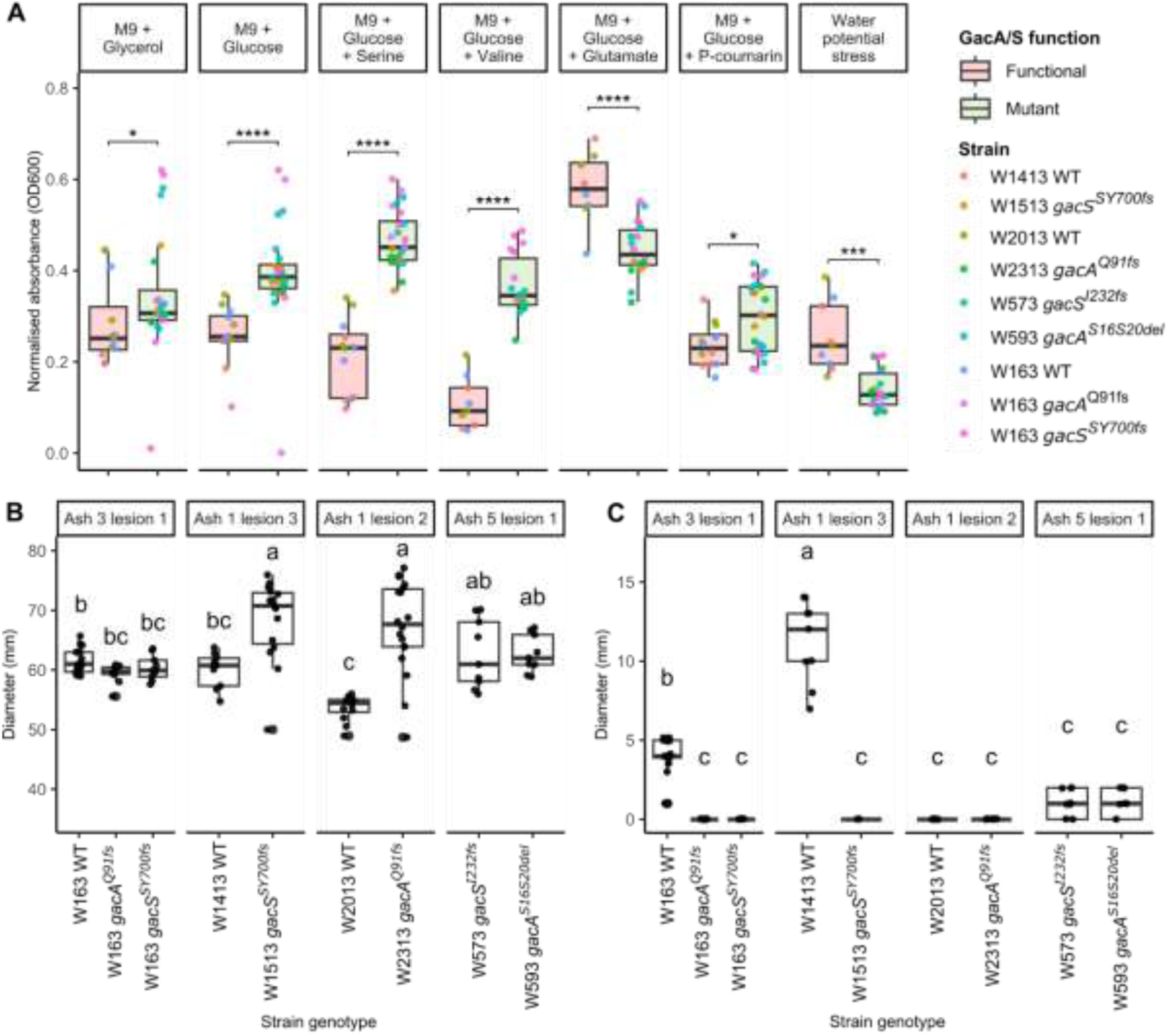
Phenotyping of *Psf* strains with functional or mutated GacA/S TCS. A) Growth in different media after 72 h. Points show means of three pseudreplicates, with three biological replicates per strain, coloured by strain and grouped by GacA/S status (mutant, green; functional, pink). A linear-mixed effects model was applied to examine the effects of treatment, GacA/S group (mutant or functional), and their interaction on bacterial growth after 72 hours, with experiment included as a random effect. Post hoc tests were conducted using Tukey-adjusted estimated marginal means. Significance is indicated (\**p* ≤ 0.05, \*\**p* ≤ 0.01, \*\*\**p* ≤ 0.001, \*\*\*\**p* ≤ 0.0001). Phenotypic assays of both WT strains (both functional and mutated *gacA*/*S*) as well as engineered W163 with *gacA* and *gacS* mutations for swimming motility and surfactant production across different trees (B and C): B) Swimming motility after 48 h in 0.25% KB agar. Points represent independent measurements. Biological replicate (n = 3) means from three technical replicates were analysed by one-way ANOVA with Tukey’s HSD (*p* < 0.05). Letters indicate significant differences. C) Biosurfactant production after 24 h on KBB. Points represent independent measurements. Biological replicate (n = 3) means from three technical replicates were analysed by one-way ANOVA with Tukey’s HSD (*p* < 0.05). Letters indicate significant differences.

Differences between groups were also observed under stress conditions. Strains with reference *gacA*/*S* maintained higher growth under osmotic stress (p < 0.001). p-coumaric acid caused a significant reduction in growth relative to glucose alone in mutant strains (p < 0.005), whereas functional strains showed no significant inhibition, indicating tolerance to p-coumaric acid (p = 0.726) (**Figure 4a; Table S9**). No significant differences were observed between groups in salt stress, quinic acid, rich medium, or antibiotic treatments (all p > 0.05) (**Table S8)**. Together, these results indicate that *gacA*/*S* function is associated with both nutrient-responsive growth and tolerance to specific environmental stresses.

While several phenotypes showed consistent associations with GacA/S functional status, others varied depending on genetic background. Within individual tissues, naturally occurring *gacA*/*S* mutants W1513 *gacS^Y700fs^* (p < 0.05) and W2313 *gac*A^Q91fs^ (p < 0.05) exhibited increased swimming motility relative to co-occurring functional strains (**Figure 4b**). However, this effect was not observed in engineered mutants in the W163 background, indicating that increased motility is not solely attributable to disruption of *gacA*/*S*.

Biosurfactant production also varied among strains. Most mutants showed no detectable production, whereas functional strains W163 and W1413 produced high levels (Figure 4c). This pattern was not consistent: mutant isolates W573 *gacS^I232fs^* and W593 *gacA^S16S20del^* produced low levels, while W2013 showed no production despite encoding a functional GacA/S system (**Figure 4c**). Given the established relationship between biosurfactant production, swarming motility, and GacA/S regulation (Burch et al., 2010, Song et al., 2016), we tested whether complementation with functional *gacA*/*S* alleles restored this phenotype. In all cases, complementation rescued biosurfactant production, including W2313 *gac*A^Q91fs^, isolated from the same tissue as W2013 (non-producer) (**Figure S13**). These results indicate that although GacA/S regulates biosurfactant production, its expression depends on genetic background, consistent with epistatic interactions that can be overridden by ectopic expression of the TCS.

When growth assays were considered together via hierarchical clustering, strains grouped primarily according to GacA*/*S functional status (**Figure S14**). Naturally evolved mutants from independent trees clustered with engineered W163 derivatives, indicating similar phenotypic reprogramming following disruption of Gac signalling in this population. Mutants further clustered by tree of isolation, suggesting additional variation is introduced by background mutations. Together these show that genotypic and phenotypic variation occurs in strains both in lesion (Ash 1 lesion 1 & 3), tree (Ash 1 & 5 vs other), and site levels (Wytham Woods vs other sites).

### Frequency-dependent maintenance of *gacA*/*S* mutants during *in planta* infection

To investigate whether the phenotype changes confer a fitness cost *in planta*, and to explore the maintenance of phenotypic heterogeneity within populations, ash tree stem pathogenicity assays were performed. Seven strain treatments were inoculated into the stem tissue of *F. excelsior* saplings: WT strains W163, W2313 *gac*A^Q91fs^, W1513 *gacS^Y700fs^*; W163 *gac*A^Q91fs^ and W163 *gacS^Y700fs^* mutants; and a mixed inoculation of W163 WT and W163 *gac*A^Q91fs^ in a 1:2 and 2:1 ratio, to test the outcome of a mixed genotype infection as seen for some lesions. After three months, lesions were scored for severity, and bacterial population size for each of the strains and strain combinations were quantified.

All lesions harboured population sizes above 3.5 log_10_ CFU mL⁻¹, 3 months post inoculation. WT W163 exhibited significantly higher population densities than both its Gac-mutated strains: W163 *gac*A^Q91fs^ (p < 0.0005) and W163 *gacS^Y700fs^* (p < 0.005) (**Figure 5a; Table S10**). This was also shown by a significant decrease in symptom development of W163 *gacS_W1513_* lesions (Dunn test; p < 0.05). A change in the symptom pattern was observed for W163 *gac*A^Q91fs^ but this was not significantly different to W163 (**Figure 5b; Table S11**). In contrast, no significant difference in symptom development or fitness was observed between WT W163 and either W2313 or W1513, suggesting other background mutations impact *in planta* fitness. Furthermore, W2313 exhibited a significantly higher population count than W163 *gac*A^Q91fs^ carrying the same mutation (p < 0.05); similarly, W1513 grew significantly better than W163 *gacS^Y700fs^*. Epistatic effects have previously been shown to affect phenotypic outcomes of *gacA*/*S* mutants (Li et al., 2021b). Notably, W1513 and W2313 strains each carry ∼70 genetic variations relative to W163, likely contributing to the differences in fitness.

**Figure 5:**
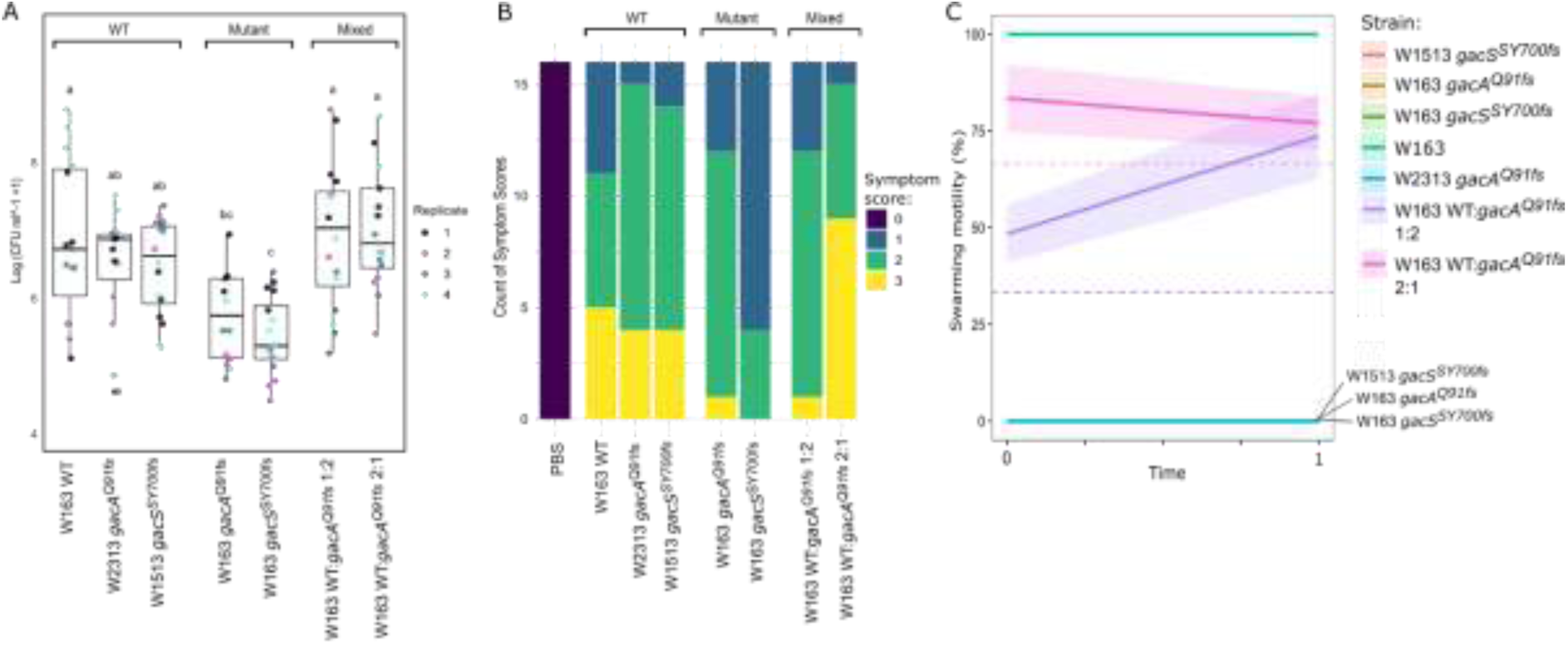
*In planta* fitness of WT and mutant *Psf* strains three months post infection. A) Bacterial population density was quantified three months post-inoculation. Mean log-transformed CFU mL⁻¹ values were calculated from three pseudoreplicates per lesion site, with four lesions per tree and four trees used as biological replicates for time 1. Differences were assessed using one-way ANOVA (significance p < 0.05). Pairwise comparisons were performed using Tukey’s Honest Significant Difference (HSD) test. Statistically distinct groups are denoted by letters a-c. B) Disease symptoms were scored on a 0-3 scale, where 0 = callused lesion, 1 = blackened lesion edges, 2 = tissue necrosis, and 3 = erumpent canker formation. C) Swarming motility of bacterial isolates from bark tissue at time 0 and after 3 months (time 1). Swarming motility was assayed for 56 isolates per lesion. At time 0, two lesions were sampled from one tree (n = 112), and at 3 months two lesions were sampled from four trees (n = 448) for each treatment. Lines show the change in the proportion of swarming isolates per treatment from time 0 to time 1, with standard deviations indicated by the shaded ribbons. Horizontal dashed lines represent the expected proportions of swarming strains in the 1:2 (purple) and 2:1 (pink) mixed treatments.

Mixed inoculations with W163 *gac*A^Q91fs^ and W163 WT strains showed *in planta* population sizes comparable to WT alone, despite the reduced fitness of W163 *gac*A^Q91fs^ inoculated in isolation. Both 1:2 and 2:1 mixtures (WT: W163 *gac*A^Q91fs^) reached significantly higher population densities than W163 *gac*A^Q91fs^ (p < 0.0001) and W163 *gacS^Y700fs^* (p < 0.001), indicating rescue effects of the WT W163 strains, independent of motile to non-motile proportions (**Figure 5a**). In contrast, only the 2:1 mixture showed significantly increased symptom development compared to W163 *gac*A^Q91fs^ alone (p < 0.05), while the 1:2 mixture was comparable (**Figure 5b; Table S11**). This suggests frequency-dependant effects of *gacA*/*S* mutants on plant physiological responses and subsequent canker formation.

We additionally tested this by quantifying an approximate ratio of swarming to non-swarming strains (**Figure 5c**). While no reversions or new *gacA*/*S* mutants arose in single-strain inoculations, mixed populations converged toward ∼75% swarming after 3 months. At time 0, swarming strains were over-represented relative to expectations in both the 1:2 (χ² = 3.7 × 10^⁻7^) and 2:1 mixtures (χ² = 3.5 × 10^⁻9^) (**Table S12**), likely due to faster growth of swarming strains on agar (data not shown). After 3 months, the proportion of W163 WT swarmer’s increased markedly in the 1:2 mixture (χ² = 4.8 × 10^⁻73^), while the 2:1 mixture showed a decline of W163 WT relative to time 0 (χ² = 3 × 10^⁻5^) (**Table S13**). These observations suggest that *gacA/S* mutants can be maintained *in planta* in single and mixed populations, but not become dominant within the tree when the wildtype is present.

## Discussion

This study examined *Pseudomonas savastanoi* pv. *fraxini* (*Psf*), a poorly understood pathogen causing erumpent cankers in European ash (*Fraxinus excelsior*). Unlike more aggressive *P. syringae* pathovars, *Psf* does not cause widespread tree mortality or major economic loss (Boa, 1978, Mansfield et al., 2012), yet it persists across the UK and Europe (EPPO, 2002). Lesions develop gradually, extending internally, establishing chronic infections which can persist for multiple years (Janse, 1982). Considering recent environmental changes, including the impact of the aggressive fungal disease causing ash dieback, shifts in host susceptibility could alter *Psf* population dynamics and selection on traits such as virulence and transmission. By integrating field sampling, comparative genomics, and phenotypic characterisation, this study investigated how *Psf* persists and diversifies within natural host populations.

### Population structure reflects local diversification and restricted ecological breadth

Whole-genome analyses revealed a highly clonal population structure, with only 833 SNPs across a 5.3 Mb core genome. Despite low overall diversity, genetic clustering was evident but did not consistently reflect geographic distance between clonal groups. Instead, diversity varied across spatial scales. In dense ash stands such as Wytham Woods, strains from nine trees were highly similar, consistent with a single clonal population present among hosts. In contrast, sparsely distributed sites such as Lathkill Dale showed a decline in diversity from site to tree to lesion, with lineages largely confined to individual hosts. These patterns indicate that even small spatial gaps can restrict dispersal. This contrasts with epidemic systems in which low diversity is maintained by frequent long-distance movement. For example, phylogroup 2d *P. syringae* on cantaloupe maintains near-identical genomes across hundreds of kilometres via rain- or irrigation-mediated dispersal (Monteil et al., 2016), while *Psa-3* populations in New Zealand accumulated minimal core variation over 12 years and did not form distinct subpopulations (Hemara et al., 2025). In *Psf*, high genetic similarity is more consistent with constrained dispersal and local transmission than with frequent gene flow.

At finer scales, lesions or clusters of trees were typically founded by single genotypes that subsequently diversified locally, generating variation without deep divergence. Repeated transmission bottlenecks may reduce effective population size within lesions, increasing the influence of genetic drift. While most *de novo* mutations are neutral or deleterious (Kimura, 1967; Lynch *et al.,* 2016), small populations limit the efficiency of purifying selection, allowing such variants to persist. Consistent with this, several members of phylogroup 3 exhibit genomic features associated with reduced effective population size. Notably, pathovars such as pv. *lachrymans* and pv. *aesculi* show expansions of IS5 family insertion sequences, which are frequently mildly deleterious (Baltrus et al., 2026). Unlike other *P. syringae* lineages frequently recovered from environmental reservoirs, phylogroup 3 members (including *Psf*) are rarely detected outside host-associated contexts (Berge et al., 2014, Morris et al., 2010). This restricted ecological range, together with reliance on host-to-host transmission, likely reduces population connectivity and opportunities for gene flow. As a result, evolutionary dynamics in *Psf* are expected to be dominated by mutation and drift rather than recombination.

Consistent with this expectation, analysis of the *Psf* pangenome revealed a large soft-core genome (∼81%), consistent with a previous analysis of six *Psf* strains (∼83%) (Moreno-Perez et al., 2020), and is substantially higher than the ∼48% species-wide core genome reported across *P. syringae* (Nowell et al., 2016). Although structural rearrangements and mobile elements contribute to some variation, clustering of the population by gene content was strongly influenced by a small number of large mobile elements. In contrast, the soft-core genome of the pandemic kiwifruit pathogen *Psa-3*, a phylogroup 2 species, accounts for 49.5% of the pangenome, with HGT shown to play a primary role in the emergence of pandemic lineages (Marcelletti et al., 2011, Butler et al., 2013, McCann et al., 2013, McCann et al., 2017). Similarly, high genome plasticity has driven the emergence and dominance of *P. syringae* pv. *tomato* race 1 on tomato crops globally, through prophages, plasmids, and genomic islands (Valenzuela and Herrera-Vásquez, 2026).

Together, these observations indicate that the *Psf* genome is relatively stable, with lower rates of HGT and recombination than in more epidemic lineages. As larger accessory genomes are typically associated with adaptation to variable environments and reduced host dependence (Dewer *et al.,* 2024), the structure of the *Psf* pangenome likely reflects both ecological restriction and potential physiological barriers to DNA uptake (Moreno-Perez et al., 2020, Perez-Martinez et al., 2007). However, as sampling here is limited to ash, broader environmental sampling will be required to determine whether recombination rates differ outside the host environment, where selective pressures may vary (Monteil et al., 2016, McCann et al., 2017).

### Regulatory mutations contribute to long term persistence

The combination of a conserved core genome and limited accessory variation suggests *Psf* encodes a large set of essential gene functions required for fitness, and mutations within existing gene networks may drive adaptation. In particular, we identified an enrichment of mutations in genes encoding two-component systems (TCSs), which regulate responses to environmental stimuli through histidine kinases and modulate downstream gene expression via response regulators, often controlling traits linked to virulence and environmental adaptation (Sultan et al., 2021). Functional validation of mutations in the GacA/S system linked genetic variation to phenotypic differences in virulence, motility, and carbon utilisation. Although not all mutations were experimentally tested, changes in amino-acid sequence can produce significant phenotypic consequences. For example, we identified a missense mutation in TolC, a membrane transport protein involved in multidrug efflux and secretion. Even small mutations in TolC can generate pronounced phenotypic effects, including altered antibiotic susceptibility (Augustus et al., 2004) and changes in bacteriophage susceptibility (German and Misra, 2001). Similarly, small mutations in the chemotaxis signalling kinase CheA, which we identified, disrupt phosphorylation signalling and lead to defects in chemotaxis and motility (Quax et al., 2018). These examples illustrate how relatively minor genetic changes in regulatory or signalling proteins can produce substantial phenotypic variation, thus these are good targets for further exploration of *Psf* behavioural changes.

Comparable patterns have been observed in *Pseudomonas aeruginosa* infections, where populations can transition from acute to chronic within the cystic fibrosis (CF) lung. During this shift, populations diversify phenotypically, including increased biofilm formation, loss of motility and altered antibiotic susceptibility (Flores-Vega et al., 2026). This is mediated by bacterial sensing of environmental signals, particularly through *de novo* mutations in regulatory pathways like GacA/S, while recombination plays a relatively minor role in within-host adaptation (Winstanley et al., 2016). Comparison of biological pathways under selection between chronic CF and acute isolates showed positive selection on quorum sensing and motility, stress resistance, regulation, efflux pumps, metabolism, and cell-surface proteins in CF isolates (Dettman and Kassen, 2021), reflecting results found in the *<u>Psf</u>* population. Notably, chronic infection in CF is not solely caused by weak immunity, but rather reflects strong and sustained selection from antibiotic exposure, oxidative and osmotic stress, and microbial competition (Winstanley et al., 2016).

In woody cankers, bacterial populations are likely exposed to external stressors such as antimicrobial phenolics, lignification responses, microbial antagonists, nutrient availability, osmotic stress, and seasonal dynamics (Baldrian, 2017, Tanase et al., 2019). In addition, a foundational histological study indicated that ash cankers are spatially heterogeneous environments (Janse, 1982). Progressive wound gum deposition, phenolic impregnation, suberisation, and repeated periderm formation generate a mosaic of colonised, necrotic, and sealed compartments that could result in divergent local selection pressures and promote persistence of distinct subpopulations. As such, the repeated and independent mutations observed in cell-surface proteins and environmental signalling pathways in highly clonal *Psf* populations may reflect adaptive fine-tuning to chronic persistence within ash cankers rather than neutral drift alone. However, this interpretation remains provisional, as direct longitudinal evidence for within-host adaptation in plant bacterial pathogens is still limited relative to *P. aeruginosa* literature.

### The role of GacA/S in within-host adaptation

The GacA/S TCS, which harboured the highest mutation count, is a master regulator that controls key behaviours underlying the pathogenicity and fitness of *Pseudomonas* (Ferreiro and Gallegos, 2021). Spontaneous *gacA/S* mutations commonly arise in biocontrol and rhizosphere strains and during growth in rich medium (Lalaouna et al., 2012, van den Broek et al., 2005), but have not been documented in phyllosphere pathogens. Independent mutations in the GacA/S system indicate positive selection on this regulatory pathway. Mutants displayed abolished swarming motility, altered amino-acid utilisation, reduced tolerance to certain stresses, and increased growth in glucose-supplemented media, whereas traits such as swimming motility, biosurfactant production, and virulence varied between strain backgrounds, reflecting epistatic interactions on some traits. These patterns were also observed in *P. protegens*, where the phenotypic outcome of *gacA* mutations was influenced by genomic context, causing either stress resistance or social traits depending on co-occurring alleles (Li et al., 2021). *gacA*/*S* mutations have been described in rhizosphere-associated *Pseudomonas*, where they are linked to plant-beneficial traits, mutualism, or potential “cheating” strategies (Li et al., 2021, Driscoll et al., 2011). In *Psf*, mutants persisted at intermediate frequencies (∼75%), consistent with frequency-dependent selection, suggesting trade-offs between individual fitness and population-level cooperation, although this requires further testing.

Selection for these mutations likely reflects the heterogeneous environment within ash lesions. Loss of GacA/S function may reduce the cost of cooperative traits while enhancing growth under nutrient limitation, providing a fitness advantage in spatially structured populations. In a system with limited recombination, such regulatory changes provide a rapid and flexible mechanism for adaptation.

### Conclusions/significance statement

*Psf* presents a model of a chronic, host-restricted pathogen in which limited dispersal, repeated bottlenecks, and small effective population size shape evolutionary dynamics. Population structure is weakly correlated to geographical distance, reflecting constrained transmission rather than widespread gene flow. Genome stability and limited HGT further restrict adaptive processes, placing greater emphasis on regulatory mutations as drivers of phenotypic diversification. Together, these findings highlight how local processes operating within hosts can generate diversity and facilitate persistence in clonal, low-virulence pathogens.

## Methods

### Strains and growth conditions

A complete list of *Psf* strains used in this study is provided in Table S1, and primers are listed in Table S2. Purified *Pseudomonas* strains were cultured at 27 °C in King’s Medium B (KB) broth or agar (KBA) (King et al., 1954). For selective isolation from plant tissue, Difco™ Pseudomonas Agar F (pancreatic digest of casein 10 g/L, proteose peptone No. 3 10 g/L, dipotassium phosphate 1.5 g/L, magnesium sulfate 1.5 g/L, agar 15 g/L) supplemented with cycloheximide and cephalexin was used. *Escherichia coli* was grown at 37 °C in lysogeny broth or agar (LB) (Bertani, 1951). Incubation times were 36–48 h for *Psf* and 16 h for *E. coli*.

Antibiotics were used at the following concentrations unless otherwise stated: cephalexin (40 mg L⁻¹), cycloheximide (100 mg L⁻¹), ampicillin (Ap, 100 µg mL⁻¹), gentamicin (Gm, 25 µg mL⁻¹), kanamycin (Km, 50 µg mL⁻¹), X-gal (40 µg mL⁻¹), nitrofurantoin (100 µg mL⁻¹), and streptomycin (100 µg mL⁻¹).

### Sampling and bacterial isolation

Bacteria were cultured from bark tissue of healthy and diseased *Fraxinus excelsior* trees from six UK woodland sites, selected for the presence of canker: Marden Park Wood, Surrey (51.2655577, -0.0550343); Wytham Woods, Oxford (51.7522, -1.25596); Lathkill Dale, Derbyshire (53.1881391, -1.7164465); Cressbrook Dale, Derbyshire (53.2550664, -1.7527726); Matlock, Derbyshire (53.1374514, -1.5544694); Isle of Mull, Scotland (56.4596986, -5.8620604); and Little Wittenham, Oxfordshire (51.63827503, -1.199820992). issue from healthy trees was classified as “Healthy,” while tissue from diseased trees was categorised as “Symptomatic” or “Non-symptomatic.” Bark samples were collected from 65 trees. Sections (2 × 4 cm) were excised using a sterilised chisel, handled with sterile gloves and bags, transported in a cool box, and processed the following day.

Two culturing methods were designed to isolate epiphytic and endophytic populations separately. However, variation in bark texture, particularly the crumbly nature of diseased tissue, led to growth from tissue particles in epiphytic washes and inconsistent sterilisation controls from the water washes. Consequently, isolates were not classified as epiphytic or endophytic in this study.

For epiphytic isolation, 2 × 1 cm bark pieces were washed in 5 mL phosphate-buffered saline (PBS; Oxoid, Hampshire, UK) for 1 h at 28 °C with shaking (200 rpm). The suspension was centrifuged (5 min, 10,000 rpm), the pellet resuspended, and briefly centrifuged to remove plant debris. From the supernatant, 100 μL was plated onto Pseudomonas isolation agar and incubated at 27 °C for 1–3 days. For endophytic isolation, bark tissue was surface-sterilised by three sequential 1 min washes in 98% ethanol, followed by sterile water rinses. Tissue was air-dried (30 min), sectioned into six pieces, and plated onto Pseudomonas agar, then incubated at 28 °C for 3–5 days and monitored daily for growth.

### Strain identification and sequencing

Isolates were initially screened using *Psf*-specific PCR primers targeting the *nicB* gene, encoding 6-hydroxy-3-succinoylpyridine 3-monooxygenase (**Table S2**). Putative *Psf*-positive strains were subsequently whole-genome sequenced (2 × 250 bp paired-end, Illumina; MicrobesNG). Long-read sequencing (Oxford Nanopore; MicrobesNG) was additionally performed for strains NCPPB1006, W163a3b1 (W163), and L1928a6b1 (L1928).

Reads were trimmed and filtered using Trimmomatic (v0.39) with parameters: ILLUMINACLIP:TruSeq2-PE.fa:2:30:10:2:True, LEADING:20, TRAILING:20, SLIDINGWINDOW:10:24, MINLEN:50 (Bolger et al., 2014). *De novo assemblies for short-read datasets were generated using SPAdes (v3.15.5) with the --careful option, while hybrid assemblies were produced using Unicycler (v0.4.0)* (Bankevich et al., 2012, Wick et al., 2017). Assembly quality was assessed using QUAST (v5.2.0) against the complete W163a3b1 reference genome(Gurevich et al., 2013). Genomes were annotated using Bakta (v1.9.4) with the --compliant flag (Schwengers et al., 2021). Poor quality genome assemblies were discarded (coverage <25 and contigs over 500bp >300 and N50 < 70,000) (**Table S1**).

All raw data and assemblies are available under BioProject PRJNA1344481.

### Recombination and mobile genetic element detection

Filtered reads from 123 *Psf* isolates were mapped to the W163a3b1 reference genome using Snippy (v4.6.0) with default parameters (Seeman, 2015). These strains included 121 isolates collected in this study, NCPPB 1006, and CFBP 5062 (Nowell et al., 2016) (**Table S1**). A core genome alignment was generated with *snippy-core*, and ambiguous sites were standardised using *snippy-clean*, with low-coverage, masked, or low-quality positions converted to “N”. The cleaned alignment was analysed with Gubbins (v3.3.1) to identify homologous recombination (default parameters), inferring recombination blocks from ≥3 SNPs within 100–10,000 bp windows (Croucher et al., 2015).

Mobile genetic elements (MGEs), including insertion sequences (IS), transposons, and phage-associated regions, were identified in the W163a3b1 reference using complementary approaches. IS elements were detected by BLASTn searches (Altschul et al., 1990) against the ISfinder database (downloaded 13/10/2024). Phage regions were identified using PHASTEST (downloaded 13/10/2024) (Wishart et al., 2023). Additionally, the Bakta GFF3 annotation was manually screened for MGE-associated terms (e.g., *“IS”*, *“transposon”*, *“conjugative”*, *“conjugation”*, and *“integrase”*).

Coordinates from all MGE detections and Gubbins-predicted recombination regions were converted to BED format, sorted, and merged using BEDTools (v2.31.1)(Quinlan and Hall, 2010), yielding a total masked region of 351,732 bp used in downstream phylogenetic analyses.

### Core genome SNP analyses

A recombination- and MGE-masked core chromosomal SNP alignment was generated using snippy-core with the --mask option, specifying the BED file derived from *Psf*-only recombination and MGE predictions. This alignment included our core 124 *Psf* isolates; *P. savastanoi* pv. *savastanoi* UPN12 (Biosample: SAMN15196490), a plasmidless derivative of *Psv* NCPPB 3335; and *P. amygdali* pv. *ciccaronei* ICMP 5710 (Biosample: SAMN03976288), which was used as the outgroup.

Phylogenetic relationships were inferred using RAxML-NG (v1.0.1) under a GTR+Γ model (Kozlov et al., 2019). Branch support was assessed with 1,000 bootstrap replicates, reporting both Felsenstein bootstrap proportions and Transfer Bootstrap Expectation (TBE) values. Nodes with support <70 were collapsed using GoTree (v0.4.5) (Lemoine and Gascuel, 2021), and trees were visualised with iTOL (v6) (Letunic and Bork, 2024).

The core SNP alignment was also used to assess genetic distance across spatial scales, within-tissue allelic variation, and isolation by distance using custom R scripts. Pairwise SNP distances were calculated with snp-dists (v0.8.2) and summarised by site, tree, and tissue; distributions were visualised with *ggridges* (Wilke, 2024). Within-tissue variation was assessed by grouping isolates by tissue ID, with SNPs classified as within-tissue alleles if present in a subset of isolates from the same sample, and further categorised as unique (single tissue) or shared across trees or sites. Geographic distances were calculated from latitude and longitude using the Haversine formula implemented in the *geosphere* package (v1.5-20) (Pebesma and Bivand, 2005, Bivand et al., 2013). Correlation between genetic and geographic distance matrices was tested using a Mantel test (Pearson method, 9,999 permutations) in *vegan* (v2.6-10) (Oksanen et al., 2025). To assess mutation distribution, SNPs were classified as coding or intergenic based on W163a3b1 annotations, and observed versus expected proportions were compared using a χ² test.

### Enrichment analysis

Genes with variants were identified by generating a binary presence–absence matrix of all mutation types (SNPs, multiple-nucleotide polymorphisms, insertions, deletions, and complex variants). The matrix was produced from Snippy per-sample variant calls across the full reference alignment (chromosome and plasmids), using custom R scripts. All reference genes were functionally annotated with all pathway types using eggNOG-mapper (Huerta-Cepas et al., 2019, Cantalapiedra et al., 2021). The eggNOG database was downloaded and installed, and emapper.py script was used to annotate input amino acid sequences using DIAMOND (Buchfink et al., 2015) with default settings, specifying “Bacteria” as the annotation taxa group. GO and KEGG annotations were saved in separate mapping files for downstream analysis.

GO and KEGG pathway enrichment analyses were performed to assess whether genes with mutations were significantly overrepresented in a specific biological pathway compared to a background set of all annotated genes in the W163a3b1 reference. Enrichment analyses were performed using the enricher function from the clusterProfile package (Wu et al., 2021). KEGG annotations were retrieved using the keggGet function from the KGGREST package . GO annotations were assigned using a custom gene-to-GO mappings, which were converted into TERM2GENE data structure splitting GO terms and assigning gene identifiers as names. Pathways were considered significantly enriched at p-value < 0.05 and a false discovery rate (q-value) < 0.1, using Benjamini-Hochberg (BH) multiple testing correction method.

A Circos plot was generated to visualise the distribution of TCS variants (Krzywinski et al., 2009). TCS variant coordinates from KEGG enrichment analysis were converted to BED format and used to extract per-strain TCS variants from VCF files using *bedtools intersect* (Quinlan and Hall, 2010). These were concatenated and formatted for Circos, including aesthetic parameters. Radius positions were assigned using a custom R script to ensure all variants from a given strain shared the same radius, with strains ordered by phylogeny. The W163 reference genome coordinates were used to generate the karyotype file, and annotated with forward and reverse CDS.

### Pangenome analysis

The pangenome dataset combined the 124 core *Psf* strains with four additional NCBI isolates: ICMP 7711 (Thakur et al., 2016), ICMP 9129, ICMP 9132, ICMP 7712 (Dillon et al., 2019b) (**Table S1**), totalling 128 genomes. These were not included in the phylogenetic analysis due to lack of SRA data. Bakta-annotated GFF3 and FNA files were analysed in Panaroo v.1.3.2 (Tonkin-Hill et al., 2020) using *strict* cleaning mode (--clean-mode strict), a pan-genome alignment (-a pan), MAFFT for alignment, and paralog merging (--aligner mafft --merge-paralogs). Gene clustering thresholds were set to --len_dif_percent 0.98 and --threshold 0.99. The resulting core gene alignment was filtered in Panaroo using the BMGE entropy filter (--core_entropy_filter) to remove poorly aligned, high-entropy core genes prior to phylogenetic analysis. A maximum-likelihood phylogeny was inferred from the filtered core gene alignment using IQ-TREE2 v.2.2.2.6 (Minh et al., 2020) with ModelFinder (-m MFP), 1,000 ultrafast bootstrap replicates (-B 1000) and SH-aLRT support (-alrt 1000), using 30 threads (-T 30) and a fixed random seed (-seed 12345).

Accessory genome clustering was examined by calculating pairwise pairwise Jaccard distances from the filtered (invariant genes removed) binary matrix using the *vegan* v2.6-10 package. A principal coordinates analysis (PCoA) was performed with the cmdscale function in base R. Distance-based redundancy analysis (dbRDA) was conducted using the capscale function with Jaccard distance, in order to obtain ordination values analogous to PCA loadings. Clade structure, manually annotated from the core gene phylogenetic tree, was included as a constraining factor to assess the proportion of genomic variation explained by phylogenetic grouping and to identify lineage-associated genes.

### *In vitro* growth assays

*In vitro* assays were performed following Li *et al.,* (2021) with minor modifications, including the use of alternative carbon sources. Overnight cultures were washed three times in PBS and adjusted to OD₆₀₀ = 0.05, then inoculated into 180 μL medium to a final OD₆₀₀ of 0.002. All assays were performed in 96-well plates (180 μL per well) at 27 °C without shaking. Growth was measured after 72 h at OD₆₀₀ using a Tecan Spark® multimode microplate reader (Thermo Fisher Scientific), with 15 s orbital shaking prior to measurement.

Stress resistance was assessed in 1 g L⁻¹ TSB supplemented with 15% polyethylene glycol (PEG-6000), 2% NaCl, or antibiotics: streptomycin (1 μg mL⁻¹), tetracycline (1 μg mL⁻¹), and penicillin (5 μg mL⁻¹). Carbon source utilisation was assessed in M9 minimal medium (M9; NaHPO₄ 34.0 g L⁻¹, KH₂PO₄ 15.0 g L⁻¹, NaCl 2.5 g L⁻¹, NH₄Cl 5.0 g L⁻¹) supplemented with individual carbon sources, either alone or with glucose (0.5 g L⁻¹). Serine, glutamic acid, valine, *p*-coumaric acid, and quinic acid were added at 0.5 g L⁻¹. Fraxetin was added at 0.1 g L⁻¹. Succinate and glycerol were tested individually at 0.5 g L⁻¹ only, with no glucose supplementation. Growth in rich medium was assessed in 1/3 King’s B broth (KBB).

Each phenotype was tested for seven strains in at least three independent experiments with three technical replicates. Growth values were normalised per experiment and phenotype by subtracting media blanks, and mean values per strain (technical replicates) were used for analysis. A linear mixed-effects model was fitted with phenotype (media) and GacA/S group (functional or mutant) as interacting fixed effects and experiment as a random effect. Post hoc comparisons were performed using Tukey-adjusted estimated marginal means. Significance was defined as *p ≤ 0.05, **p ≤ 0.01, ***p ≤ 0.001, ****p ≤ 0.0001.

### Motility and biosurfactant production

Motility was assessed on King’s B agar (KBA) plates containing 0.5% agar for swarming and 0.25% agar in 1/3-strength KBA for swimming. Plates were prepared by pouring 30 mL sterile agar (cooled to 50 °C) into 90 mm Petri dishes, then drying in a laminar flow hood for 30 min, rotating after 15 min to ensure even drying. For swimming assays, a single colony was stab-inoculated into the centre of the agar without contacting the plate base. For swarming assays, 3 μL of overnight culture (OD₆₀₀ = 1.0) was spotted onto the centre of each plate. Plates were incubated at 22 °C in the dark, in a single layer, for 72 h. Plates were imaged using a gel documentation system (Analytica Jena UVP GelSolo), and colony expansion was quantified in ImageJ by measuring the maximum colony diameter, using the 90 mm plate as a scale reference.

Biosurfactant detection was done via an assay with atomized oil performed following Burch *et al.,* 2010. Briefly, overnight cultures were adjusted to 0.7 OD_600_ in fresh KBB, and 5µl was spotted onto KBA plates and incubated overnight at 22°C. An airbrush (0.3 mm nozzle, 48 PSI) was used to apply a fine mist of mineral oil (Sigma-Aldrich, St. Louis, MO, USA) with an air pressure of between 15 and 20 lb/in^2^. Biosurfactant halos were measured with a ruler from the edge of the bacterial colony to the edge of halo and photographed.

Each assay was performed in three independent experiments with three technical replicates per experiment. The mean of technical replicates was used for analysis. Differences between strains were assessed using one-way ANOVA, followed by Tukey’s honestly significant difference (HSD) post hoc test.

### Pathogenicity assays

Pathogenicity of Psf strains was assessed on *Fraxinus excelsior* saplings collected from Norbury Park, Staffordshire, UK (January 2023) with land manager permission. Seedlings were sourced from a ∼50 × 50 m area surrounding a single ash tree and assumed to share a common maternal origin. Saplings were transferred to 5 L pots containing a peat-free soil–perlite mix and maintained under greenhouse conditions (natural photoperiod, 16 h light/8 h dark, 18–20 °C). At inoculation, trees were ∼2 years old, and ∼6-month-old stem tissue was used.

Overnight bacterial cultures were adjusted to 10⁸ CFU/mL (OD₆₀₀ = 0.5) in PBS (strains in Table 4.1). For mixed inocula, WT W163 and W163 *gacAQ91fs* were combined at 2:1 or 1:2 ratios. Saplings were surface-sterilised with 70% ethanol, and wounds (∼1 cm) were created by horizontal bark incisions. Four lesions per tree (A–D) were randomly assigned, inoculated with 20 μL, air-dried (15 min), and sealed with parafilm for 1 month. Five plants per strain were used (four inoculation points each). One plant was sampled immediately post-inoculation to determine initial CFU (n = 4), and the remaining four were analysed at 3 months (n = 16).

For CFU quantification, ∼1 cm tissue (0.1 ± 0.05 g) from each inoculation site was excised, homogenised in 1 mL PBS (4 m/s, 40 s), serially diluted (neat to 10⁻⁷), and plated (3 μL, three technical replicates) on KBA with cycloheximide and cephalexin. CFU/mL was calculated using standard dilution-based methods. Symptoms were photographed at 3 months and scored (0–3): 0, no disease; 1, blackened lesion edges; 2, blackening with necrosis and/or corky outgrowth; 3, erumpent canker formation.

To assess in vivo swarming ratios, 50 μL of 10⁻¹–10⁻³ dilutions from two lesions per tree (A, B) were plated on selective KBA and incubated at 27 °C (48 h). From each lesion, 56 colonies were randomly selected and stab-inoculated onto 0.5% swarming agar (500 × 500 mm plates, 300 mL medium), incubated at 22 °C, and scored after 24 h as swarming/non-swarming.

Differences in bacterial population size (log₁₀ CFU/mL) were analysed by one-way ANOVA on log-transformed data with Tukey’s HSD. Symptom severity was analysed using a Kruskal–Wallis test with Dunn’s post hoc test and Benjamini–Hochberg correction (rstatix). Observed vs expected swarming ratios (1:2, 2:1) were tested using Pearson’s chi-squared goodness-of-fit tests at both individual inoculation points and aggregated across replicates and timepoints (0 and 3 months).

### Preparation of chemically competent and electrocompetent cells

*Pseudomonas* strains were made electrocompetent by washing 1 mL overnight cultures (∼1.4 OD_600_) in 750 µL of ice-cold 0.5 M sucrose three times (16,200 g for 5 minutes) on ice. The final pellet was resuspended in 300 µL ice-cold 0.5 M sucrose, aliquoted in 50 µL and stored at -80°C.

To prepare chemically competent cells of S17-1 λpir *E. coli*, a single colony was inoculated into 50 mL LBB in a 300 mL flask and incubated at 37°C (60 rpm for 16 hours). One mL of overnight culture was inoculated into 100 mL LB in a 500mL flask and incubated again at 37°C at 300rpm until OD_600_ = 0.85 to 0.98 (mid-logarithmic phase). On ice, the bacterial suspension was transferred to four 50 mL falcon tubes. Cells were washed in a 2.5 mL mixture of ice cold 80 mM CaCl_2_ and 50 mM MgCl_2_ three times with centrifugation of 2,400 g for 10 min at 4°C. Cells were resuspended in ice-cold 0.1 M CaCl_2_ at 5 × 10^9^ cells/ml and glycerol added (50%). Aliquots of 50 µL were stored at -80°C.

### Two-step allelic exchange

Allelic exchange was performed following the method of Hmelo *et al.,* (2015), with the following modifications. Mutant alleles of *gacA* (588 bp) and *gacS* (2,754 bp) were PCR-amplified from the non-motile strains W2313 GacA^Q91fs^ and W1513 GacS*^Y700fs^*, respectively, including ∼450 bp of homologous flanking sequence (100% sequence identity) on either side of the mutant allele to facilitate homologous recombination in the recipient strain W163. PCR reactions were prepared with 1X Q5® High-Fidelity Master Mix (New England Biolabs, UK), 10 µM each of forward and reverse primers, 10 ng of template DNA, and nuclease free water to 50 µL total volume. PCR products were verified by agarose gel electrophoresis, excised, and purified using the Monarch® DNA Gel Extraction Kit (NEB, Hitchin, UK). Purified PCR products and the vector pK18mob*sacB* (Km^R^) were digested with High-Fidelity *Eco*RI and *Bam*HI (each were 20,000 units µL^-1^; NEB, UK) following the manufacturer’s protocol. Digestion reaction was prepared using 1µL DNA, 10X rCutSmart® Buffer, 20 units each of *Eco*RI or *Bam*HI, and NFW to a final volume of 25 µL (insert) or 50 µL (vector). Reactions were incubated at 37°C for 60 minutes followed by heat inactivation at 65°C for 20 minutes. Digested products were resolved by 1% agarose (Meridian Bioscience, Cincinnati, OH, USA) gel electrophoresis (250 V) and extracted using the Monarch® kit. DNA concentrations were quantified using a NanoDrop 1000 spectrophotometer (Thermo Scientific).

Ligation reactions were set up using T4 DNA Ligase (NEB), where the required mass insert was calculated using the NEBioCalculator tool, targeting 50 ng of plasmid in a 10 µL reaction. Final mixtures included 50 ng vector, the required mass of the insert (ng) for molar 3:1 ratio, 5X NEB ligase buffer and 5U/µL T4 DNA Ligase (NEB, UK), and NFW to final volume of 10 µL. Ligation reactions were incubated at 4°C overnight (∼19 hours)and then transformed into chemically competent *E. coli* DH5α (C2987H, NEB) via heat shock. Approximately 100 ng of the purified ligation reaction was incubated with 50 µL ice-thawed cells on ice for 30 minutes, heat-shocked at 42°C for 30 seconds, and placed on ice for 5 minutes. SOC medium (950 µL) was added, and cells were incubated at 37°C with shaking for 2 hours. Cells were pelleted (10,000 rpm, 7 minutes) and resuspended in 100 µL of SOC medium. Aliquots of 80 µL and 20 µL were plated onto LBA supplemented with X-gal and kanamycin (km). Plates were incubated at 37°C for 18 hours, and transformants were identified by blue/white screening. Positive clones were stored at −80°C in glycerol stocks. Recombinant plasmids were extracted using the PureLink™ Quick Plasmid Miniprep Kit (Invitrogen). Insert size and orientation were confirmed by PCR and Sanger sequencing using M13 primers. Verified constructs were transformed into *E. coli* S17-1 λpir by heat shock as described above, for subsequent biparental mating. Transformed S17-1 λpir cells were plated on LBA + Km (50 µL and 1:5 dilution) and incubated at 37°C for 16–24 hours. Colonies were verified by a second round of PCR using M13 primers.

### Biparental mating

Overnight cultures of donor cells S17-1 λpir (OD_600_ ∼0.6) containing the plasmid construct and the recipient strain W163 (OD_600_ >1.2) were separately washed three times in phosphate-buffered saline to remove any residual antibiotics. Donor and recipient cells were mixed in 1:1, 1:2 and 2:1 ratios, pelleted (9,600 x g, 5 minutes) and resuspended in 50 µL LBB. The mixture was spotted onto LBA plates in a droplet and air-dried before incubation at 30°C for 48 hours. After incubation, the entire spot was spread on LBA containing Km and Nitrofurantoin (NF) with a sterile loop and spread with a sterile L-shaped spreader. Plates were incubated at 27°C for 2-5 days. Resulting colonies were re-streaked onto LBA + Km + NF and incubated at 27°C for 2 days.

### Sucrose counterselection

Single colonies were streaked onto KBA plates supplemented with 10% sucrose plates, and in parallel, onto KBA + Km control plates. Plates were incubated at 27°C for two days. Colonies that grew on KBA + sucrose but not on KBA + Km were selected as putative double recombinants and confirmed by PCR and Sanger sequencing of the *gacA* or *gacS* gene.

### Deletion mutant complementation

Complementation of deletion mutants was performed using the broad-host-range plasmid pBBR1MCS-5 (Gm^R^). W163a3b1 *GacA* and *GacS* genes, including 300-400bp upstream to preserve native promotor control and <10bp downstream to avoid adjacent start sites, were amplified from W163 strain by PCR as previously described. PCR products and the pBBR1MCS-5 vector were digested with *Apa*I and *Spe*I restriction enzymes (Thermo Fisher Scientific), following the manufacturer’s protocol. Digestions were resolved by gel electrophoresis, gel-excised, purified, ligated and transformed into *E. coli* DH5α as previously described for allelic exchange. Ligated constructs were confirmed by sanger sequencing. Verified constructs were transformed into the relevant recipients by electroporation as described below. Specifically, pBBR1MCS-5 + G*acA_W163* was transformed into W163 WT, W163 g*ac*A^Q91fs^ and W2313 gacA^Q91fs^, whilst pBBR1MCS-5 + *gacS_W163* was transformed into W163 WT, W163 gacS*^Y700fs^* and W1513 gacS*^Y700fs^*. The empty pBBR1MCS-5 vector was transformed into each strain as a negative control.

### Transformation by electroporation

Electrocompetent cells of *Pseudomonas* (50 µL) were thawed on ice for 10 minutes. Approximately ∼100 ng plasmid DNA was added to each sample and incubated on ice for 30 minutes without vortexing. Cells were transferred to pre-chilled electroporation cuvettes (2 mm) (Bio-Rad, UK). Electroporation was carried out using a Bio-Rad Gene Pulser under the following conditions: 2.5 kV, 25 µF capacitance, and 400 Ω resistance. Immediately following the pulse, 1 mL of room-temperature KBB was added to each cuvette to resuspend the cells. Cells were transferred to a 1.5 mL Eppendorf and incubated at 28°C for 4 hours with shaking at 200rpm. Cultures were centrifuged, and pellets were resuspended in 100 µL KBB. Aliquots of 20 µL and 80 µL were plated onto KBA + Gm and incubated at 28°C for 48 hours. Transformants were screened by PCR using M13 primers and confirmed by Sanger sequencing.

## Supporting information

All supplementary figs and tables

## Acknowledgments

We thank the JABBS Foundation for funding to RWJ to support KGH, SD and DVV. We are especially thankful to Eric Boa providing key guidance on identifying woodlands and conducting preliminary site visits for sampling as well as shipping the sample from the Isle of Mull. We thank Cayo Ramos and his team for assistance with training in sapling inoculation assays. We thank James Brown for provision of ash trees, Keith Kirby and Ben Lebas for assistance in identifying and obtaining permissions for suitable sampling sites, and Vanja Milenkovic, Emily Grace, Amy Webster and Theophile Muller for help with fieldwork and bacterial isolations. Thanks also to Sophie Powell and Andrea Vadillo Dieguez for help with the mutant phenotyping assays and pathogenicity assays. We are very grateful to Natural England and Forest Research for providing sampling permissions.

