## Supplementary material for "Mutations in bacterial regulatory genes are linked with chronic ash tree infections": All supplementary figs and tables

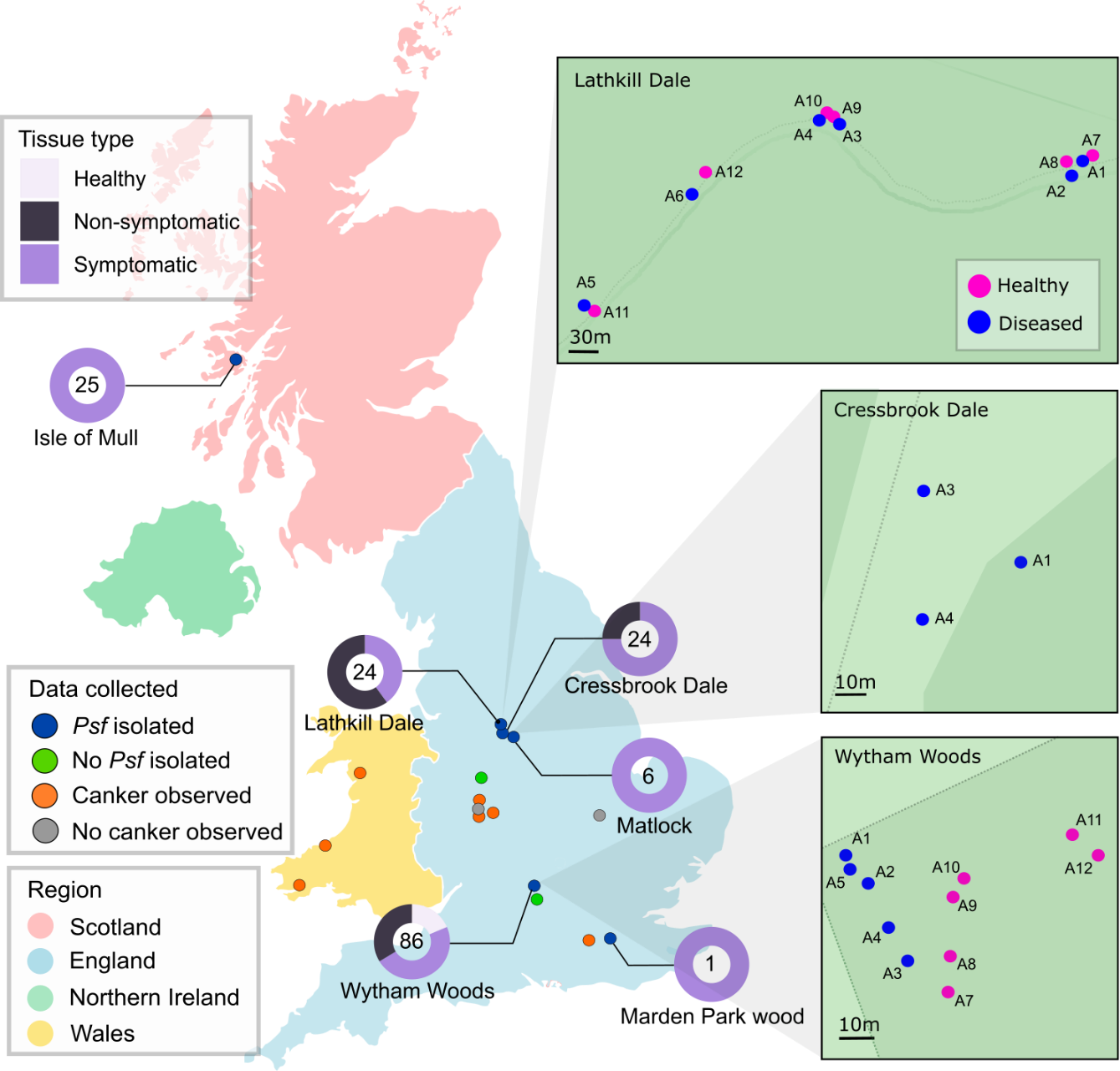

Figure S1 Locations of woodland sites visited and sampled in the UK. Location of UK woodland sites are annotated by circles. On the main map, blue and green circles denote sampled sites from which *Psf* was successfully or unsuccessfully isolated, respectively. Orange markers indicate sites where canker was observed but not sampled, whilst grey markers indicate sites which were visited but no cankers were observed. Pie charts show the number of *Psf* isolated per site, coloured by the tissue type from which they were isolated: Healthy (pale purple, Wytham Woods only), non-symptomatic tissue from diseased trees (dark purple) and symptomatic tissue from diseased trees (purple). In large green boxes, the location of trees within each site where *Psf* was successfully isolated from multiple trees. Healthy trees are indicated by pink circles, whilst cankered trees are shown in blue. Map data from OpenStreetMap, licenced under the Open Data Commons Open Database Licence (ODbL) by the OpenStreetMap Foundation (OSMF).

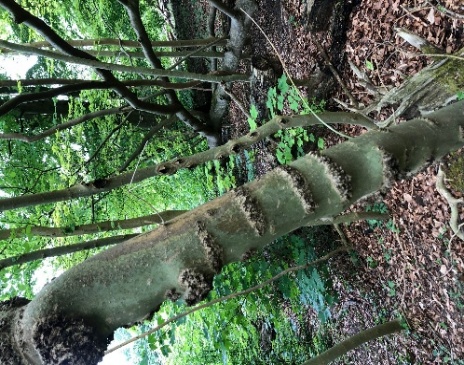

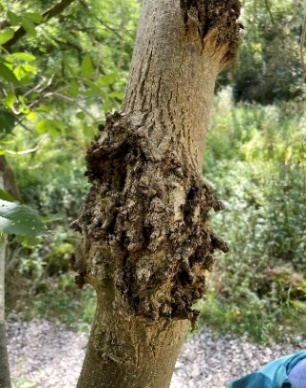

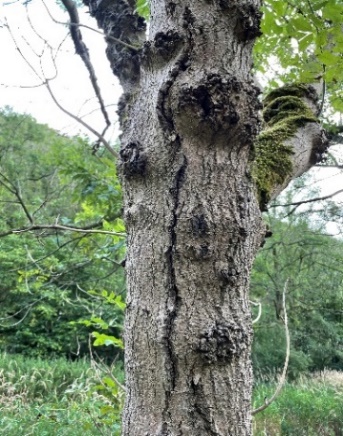

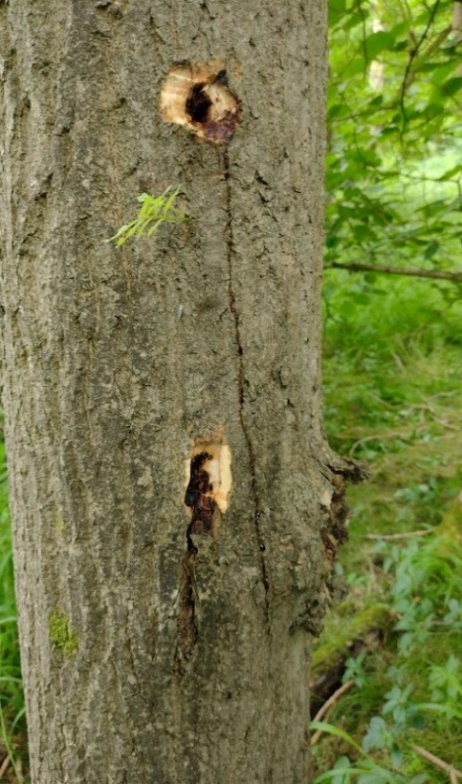

Figure S2 Symptoms of ash canker caused by *Psf*. Left to right) Letterbox cankers spread longitudinally up the main trunk with shothole cankers present on rear tree (Marden Park Wood); erumpent canker (Lathkill Dale); lateral cracks (Lathkill Dale); bleeding lesion (Wytham woods). Photographs taken by author.

**Table S1 Quality of *Psf* genome assembly (SPAdes) and annotation (Bakta).** Tissue health is abbreviated: Healthy (H), Symptomatic (S) and Non-symptomatic (NS). Strains which did not pass QC and were not included in the phylogeny are denoted with *. Assembled genomes were downloaded from NCBI: ICMP7711, ICMP7712, ICMP9129, ICMP9132 (Dillon *et al.,* 2019b). Raw fastQ files were downloaded for strain CFPB5062 (Nowell *et al.,* 2016).

| **Strain** | **Tissue health** | **Clade** | **Contigs >500bp** | **Total length >500bp** | **GC (%)** | **N50** | **Unaligned length** | **CDSs** | **NCBI Biosample** |
| --- | --- | --- | --- | --- | --- | --- | --- | --- | --- |
| C130a3b1 | S | 8a | 276 | 6225316 | 57.98 | 71636 | 206140 | 5755 | SAMN52817268 |
| C150a3b1 * | S |  | 279 | 6224110 | 57.97 | 71828 | 210761 | 5752 | SAMN52817269 |
| C170a3b3 * | S |  | 283 | 6227390 | 57.97 | 71407 | 201954 | 5745 | SAMN52817270 |
| C250a4b2 | S | 8 | 268 | 6231054 | 57.98 | 71913 | 203825 | 5756 | SAMN52817271 |
| C270a4b3 | S | 8a | 280 | 6226128 | 57.98 | 71828 | 203091 | 5749 | SAMN52817272 |
| C400a1s2 | NS | 8c | 282 | 6274659 | 57.91 | 71502 | 202414 | 5816 | SAMN52817273 |
| C410a1s1 | NS | 8c | 279 | 6233079 | 57.93 | 68795 | 172126 | 5763 | SAMN52817274 |
| C420a1s2 | NS | 8c | 293 | 6279210 | 57.9 | 71456 | 199434 | 5823 | SAMN52817275 |
| C430a1s3 | NS | 8b | 291 | 6258447 | 57.92 | 68484 | 171585 | 5800 | SAMN52817276 |
| C450a3s3 | NS | 8b | 300 | 6260809 | 57.92 | 68708 | 173355 | 5819 | SAMN52817277 |
| C50a1b2 * | S |  | 319 | 6284871 | 57.9 | 66425 | 201279 | 5859 | SAMN52817278 |
| C530a1b2 | S | 8c | 281 | 6290088 | 57.9 | 68708 | 200120 | 5835 | SAMN52817279 |
| C560a1b2 | S | 8c | 288 | 6281475 | 57.9 | 68795 | 199973 | 5825 | SAMN52817280 |
| C610a1b3 | S | 8b | 290 | 6254849 | 57.92 | 71537 | 175742 | 5797 | SAMN52817281 |
| C630a3b1 | S | 8a | 280 | 6227385 | 57.98 | 68708 | 203720 | 5759 | SAMN52817282 |
| C680a3b3 | S | 8a | 260 | 6176666 | 58.01 | 71828 | 188200 | 5689 | SAMN52817283 |
| CFBP5062 | NA | 6a | 330 | 6273967 | 58 | 68900 | 165797 | 5766 | SAMN03992201 |
| ICMP7711 | NA | NA | NA | 6023921 | 58 | 38704 | NA | 5576 | SAMN03976404 |
| ICMP7712 | NA | NA | NA | 5928827 | 58.1 | 39170 | NA | 5443 | SAMN03976405 |
| ICMP9129 | NA | NA | NA | 5998153 | 58.1 | 46711 | NA | 5496 | SAMN03976407 |
| ICMP9132 | NA | NA | NA | 6045945 | 58.1 | 46511 | NA | 5543 | SAMN03976406 |
| IoMN10D5R22 | S | 2 | 268 | 6080187 | 58 | 68717 | 54110 | 5592 | SAMN52817284 |
| IoMN14D1R21 | S | 2 | 268 | 6062157 | 58.01 | 68717 | 50553 | 5575 | SAMN52817285 |
| IoMN15D1R31 | S | 2 | 273 | 6110487 | 57.99 | 68794 | 75229 | 5644 | SAMN52817286 |
| IoMN18D2R21 | S | 3 | 267 | 6091973 | 58 | 71405 | 58520 | 5616 | SAMN52817287 |
| IoMN19D2R31 | S | 3 | 283 | 6116181 | 57.99 | 71641 | 74266 | 5632 | SAMN52817288 |
| IoMN28D4R41 | S | 3 | 292 | 6136329 | 57.99 | 71733 | 74563 | 5663 | SAMN52817289 |
| IoMN3D1R32 | S | 2 | 280 | 6108836 | 57.99 | 71405 | 75406 | 5635 | SAMN52817290 |
| IoMN8D4R22 | S | 3 | 263 | 6092489 | 58.02 | 71528 | 73992 | 5614 | SAMN52817291 |
| L1217a1b4 | NS | 7b | 312 | 6343501 | 57.89 | 69589 | 245701 | 5896 | SAMN52817292 |
| L1227a1b4 | NS | 7b | 310 | 6347676 | 57.89 | 68797 | 247401 | 5904 | SAMN52817293 |
| L1277a1b6 | NS | 7b | 325 | 6370861 | 57.89 | 69589 | 248122 | 5933 | SAMN52817294 |
| L1327a1b5 | NS | 7b | 335 | 6373608 | 57.89 | 68797 | 249055 | 5925 | SAMN52817295 |
| L1738a1b4 | NS | 7b | 309 | 6346519 | 57.89 | 68704 | 238969 | 5895 | SAMN52817296 |
| L1768a4b4 | NS | 6 | 291 | 6234392 | 57.87 | 71407 | 209494 | 5770 | SAMN52817297 |
| L1778a4b5 | NS | 6 | 296 | 6243169 | 57.87 | 68797 | 206138 | 5792 |  |
| L1818a6b4 | NS | 4b | 282 | 6265381 | 57.95 | 71821 | 193125 | 5802 | SAMN52817298 |
| L1928a1b6 | NS | 7b | 6 | 6699942 | 57.8 | 6.00E+06 |  | 6212 | SAMN52817299 |
| L1958a2b6 | NS | 7b | 330 | 6386294 | 57.88 | 68704 | 254505 | 5964 | SAMN52817300 |
| L2008a3b4 | NS | 6 | 257 | 6162598 | 57.91 | 70694 | 188857 | 5694 | SAMN52817301 |
| L2018a3b5 | NS | 6 | 280 | 6230428 | 57.87 | 71405 | 229522 | 5801 | SAMN52817302 |
| L777a5b1 | S | 4b | 259 | 6232692 | 57.97 | 71390 | 170973 | 5778 | SAMN52817303 |
| L787a5b1 | S | 4b | 275 | 6264495 | 57.96 | 71425 | 195421 | 5800 | SAMN52817304 |
| L797a5b1 | S | 4b | 286 | 6266933 | 57.96 | 71821 | 197935 | 5817 | SAMN52817305 |
| L807a5b1 | S | 4b | 304 | 6272173 | 57.95 | 71390 | 201584 | 5832 | SAMN52817306 |
| L817a5b2 | S | 4a | 286 | 6252257 | 57.97 | 68339 | 192030 | 5807 | SAMN52817307 |
| L827a5b2 | S | 4a | 266 | 6213953 | 57.99 | 71405 | 166021 | 5746 |  |
| L837a5b2 | S | 4a | 271 | 6218217 | 57.98 | 71405 | 172881 | 5762 | SAMN52817308 |
| L898a5b2 | S | 4a | 263 | 6217643 | 57.98 | 71821 | 172969 | 5756 | SAMN52817309 |
| MardA1S4125 | S | 5 | 275 | 6181720 | 57.99 | 71426 | 144794 | 5718 | SAMN52817310 |
| MatA1109 | S | 7a | 301 | 6263651 | 57.92 | 68795 | 172038 | 5824 | SAMN52817311 |
| MatA2111 | S | 7a | 294 | 6240265 | 57.94 | 71522 | 157534 | 5789 | SAMN52817312 |
| MatA2122 | S | 7a | 312 | 6278750 | 57.92 | 71522 | 172852 | 5824 | SAMN52817313 |
| MatA2124 | S | 7a | 292 | 6266369 | 57.92 | 71405 | 171285 | 5802 | SAMN52817314 |
| NCPPB1006 | S | 1 | 5 | 6298151 | 58 | 6.00E+06 | NA | 5785 | SAMN52648148 |
| W10013a5b5 | NS | 9 | 250 | 6055050 | 58.05 | 71871 | 18363 | 5548 | SAMN52817315 |
| W1003a4b5 | NS | 9 | 263 | 6140877 | 57.99 | 71830 | 53220 | 5676 | SAMN52817316 |
| W1013a1b2 | S | 9 | 283 | 6153012 | 57.98 | 71405 | 57185 | 5671 | SAMN52817317 |
| W1013a4b6 | NS | 9 | 274 | 6149036 | 57.98 | 68795 | 57372 | 5672 | SAMN52817318 |
| W10213a5b6 * | NS |  | 262 | 6054719 | 58.05 | 71875 | 18969 | 5546 | SAMN52817319 |
| W1023a4b6 | NS | 9 | 296 | 6167339 | 57.98 | 71830 | 58046 | 5699 | SAMN52817320 |
| W10313a5b6 | NS | 9 | 252 | 6059439 | 58.04 | 72398 | 20565 | 5548 | SAMN52817321 |
| W1033a4b6 | NS | 9 | 286 | 6155981 | 57.98 | 71733 | 57415 | 5682 | SAMN52817322 |
| W1043a5b6 | NS | 9 | 294 | 6149665 | 57.99 | 71830 | 47314 | 5668 | SAMN52817323 |
| W1053a5b6 | NS | 9 | 297 | 6161443 | 57.99 | 71830 | 51245 | 5674 | SAMN52817324 |
| W1063a7b2 | H | 9 | 284 | 6156185 | 57.98 | 71871 | 58003 | 5681 | SAMN52817325 |
| W1073a7b2 | H | 9 | 282 | 6155949 | 57.98 | 71871 | 58106 | 5679 | SAMN52817326 |
| W1083a7b2 | H | 9 | 286 | 6150370 | 57.98 | 71405 | 57644 | 5664 | SAMN52817327 |
| W1093a7b2 | H | 9 | 277 | 6149239 | 57.99 | 71830 | 58568 | 5667 | SAMN52817328 |
| W1103a7b2 | H | 9 | 263 | 6083335 | 58.02 | 72184 | 0 | 5580 | SAMN52817329 |
| W1113a1b2 | S | 9 | 283 | 6106366 | 58.01 | 71871 | 26798 | 5616 | SAMN52817330 |
| W1113a7b3 | H | 9 | 257 | 6080441 | 58.02 | 73835 | 0 | 5574 | SAMN52817331 |
| W1123a7b3 | H | 9 | 297 | 6157954 | 57.98 | 71830 | 56537 | 5675 | SAMN52817332 |
| W113a1b1 | S | 9 | 288 | 6116531 | 58.01 | 68795 | 27924 | 5626 | SAMN52817333 |
| W1143a7b3 | H | 9 | 295 | 6158205 | 57.98 | 71871 | 57119 | 5690 | SAMN52817334 |
| W1153a9b3 | H | 9 | 295 | 6163900 | 57.98 | 71830 | 57792 | 5693 | SAMN52817335 |
| W1163a11b1 | H | 9 | 289 | 6150446 | 57.99 | 71830 | 57439 | 5671 | SAMN52817336 |
| W1173a11b1 | H | 9 | 293 | 6160501 | 57.99 | 71871 | 55205 | 5695 | SAMN52817337 |
| W1213a1b2 | S | 9 | 279 | 6152049 | 57.98 | 71405 | 57712 | 5671 | SAMN52817338 |
| W12913a12b2 * | H |  | 285 | 6145791 | 57.99 | 71871 | 58091 | 5676 | SAMN52817339 |
| W13013a12b2 | H | 9 | 275 | 6147076 | 58 | 71871 | 56353 | ` | SAMN52817340 |
| W1313a1b2 | S | 9 | 258 | 6099204 | 58.03 | 71830 | 57855 | 5612 | SAMN52817341 |
| W13a1b1 | S | 9 | 294 | 6164718 | 57.98 | 71871 | 57500 | 5674 | SAMN52817342 |
| W1413a1b2 | S | 9 | 254 | 6097803 | 58.03 | 72228 | 58338 | 5610 | SAMN52817343 |
| W1513a1b2 | S | 9 | 280 | 6152743 | 57.98 | 71830 | 58730 | 5687 | SAMN52817344 |
| W1613a1b2 | S | 9 | 279 | 6107887 | 58.01 | 71871 | 27257 | 5613 | SAMN52817345 |
| W163a3b1 | S | 10 | 7 | 6525693 | 57.9 | NA | 0 | 6053 | SAMN52659773 |
| W1713a1b3 | S | 9 | 280 | 6150099 | 57.98 | 71837 | 58596 | 5674 | SAMN52817347 |
| W183a3b2 | S | 9 | 253 | 6058299 | 58.05 | 71875 | 22733 | 5561 | SAMN52817348 |
| W1913a1b3 | S | 9 | 267 | 6145826 | 57.98 | 71871 | 57803 | 5672 | SAMN52817349 |
| W193a3b2 | S | 9 | 266 | 6061188 | 58.04 | 68795 | 24531 | 5547 | SAMN52817350 |
| W2013a1b3 | S | 9 | 292 | 6162161 | 57.98 | 69690 | 58177 | 5686 | SAMN52817351 |
| W203a3b2 | S | 9 | 257 | 6060356 | 58.05 | 71405 | 23092 | 5552 | SAMN52817352 |
| W2113a1b3 | S | 9 | 283 | 6156153 | 57.98 | 71405 | 58322 | 5675 | SAMN52817353 |
| W213a1b1 | S | 9 | 273 | 6146910 | 58.01 | 71871 | 44942 | 5663 | SAMN52817354 |
| W2313a1b3 | S | 9 | 282 | 6158613 | 57.98 | 71871 | 58074 | 5682 | SAMN52817355 |
| W2413a1b3 | S | 9 | 276 | 6104929 | 58.01 | 71830 | 27624 | 5615 | SAMN52817356 |
| W273a3b5 | NS | 10 | 257 | 6081847 | 58.03 | 73834 | 0 | 5571 | SAMN52817357 |
| W313a1b1 * | S |  | 320 | 6172973 | 57.98 | 71919 | 59887 | 5695 | SAMN52817358 |
| W343a4b1 | S | 9 | 286 | 6153968 | 57.99 | 71830 | 50249 | 5677 | SAMN52817359 |
| W413a4b3 | S | 9 | 281 | 6132216 | 58 | 68669 | 54260 | 5647 | SAMN52817360 |
| W423a4b3 | S | 9 | 284 | 6136464 | 58 | 71405 | 53709 | 5660 | SAMN52817361 |
| W433a4b3 * | S |  | 308 | 6122214 | 58.01 | 60243 | 54384 | 5643 | SAMN52817362 |
| W43a1b1 | S | 9 | 275 | 6154835 | 57.98 | 71871 | 58309 | 5677 | SAMN52817363 |
| W513a1b1 * | S |  | 287 | 6148344 | 57.99 | 71830 | 54546 | 5664 | SAMN52817364 |
| W573a5b1 | S | 9 | 248 | 6056237 | 58.05 | 71871 | 18798 | 5548 | SAMN52817365 |
| W583a5b1 | S | 9 | 246 | 6056360 | 58.05 | 71875 | 18359 | 5559 | SAMN52817366 |
| W593a5b1 | S | 9 | 243 | 6054188 | 58.05 | 71871 | 18779 | 5552 | SAMN52817367 |
| W603a5b1 | S | 9 | 250 | 6053365 | 58.05 | 68795 | 18925 | 5552 | SAMN52817368 |
| W613a1b1 | S | 9 | 305 | 6169042 | 57.98 | 71871 | 56903 | 5696 | SAMN52817369 |
| W6913a4b2 | S | 9 | 287 | 6135365 | 58 | 71875 | 51778 | 5658 | SAMN52817370 |
| W703a5b4 | S | 9 | 282 | 6106737 | 58.01 | 71405 | 27804 | 5602 | SAMN52817371 |
| W7113a4b2 | S | 9 | 280 | 6090223 | 58.03 | 68795 | 28557 | 5589 | SAMN52817372 |
| W7213a4b2 | S | 9 | 278 | 6141804 | 57.99 | 71830 | 56898 | 5660 | SAMN52817373 |
| W723a5b4 | NS | 9 | 286 | 6105626 | 58.01 | 71830 | 27880 | 5607 | SAMN52817374 |
| W803a11b1 | H | 9 | 271 | 6086975 | 58.02 | 74124 | 0 | 5593 | SAMN52817375 |
| W813a1b1 | S | 9 | 288 | 6159767 | 57.98 | 71405 | 56541 | 5684 | SAMN52817376 |
| W883a2b4 | NS | 9 | 272 | 6100467 | 58.01 | 71830 | 29343 | 5616 | SAMN52817377 |
| W8913a5b3 | S | 9 | 251 | 6055898 | 58.05 | 71871 | 18927 | 5547 | SAMN52817378 |
| W893a3b4 | NS | 9 | 284 | 6153912 | 57.99 | 71830 | 52241 | 5680 | SAMN52817379 |
| W903a3b4 | NS | 9 | 290 | 6157957 | 57.99 | 71405 | 51059 | 5683 | SAMN52817380 |
| W9113a5b3 | S | 9 | 249 | 6057019 | 58.05 | 72398 | 18823 | 5550 | SAMN52817381 |
| W913a1b2 | S | 9 | 260 | 6098822 | 58.03 | 71405 | 60101 | 5620 | SAMN52817382 |
| W913a3b4 | NS | 9 | 285 | 6145373 | 57.99 | 71405 | 51495 | 5664 | SAMN52817383 |
| W9213a5b3 | S | 9 | 251 | 6053436 | 58.05 | 68795 | 18782 | 5550 | SAMN52817384 |
| W923a3b4 | NS | 9 | 307 | 6164518 | 57.99 | 71405 | 51698 | 5689 | SAMN52817385 |
| W9313a5b4 | NS | 9 | 241 | 6055888 | 58.05 | 71871 | 18359 | 5550 | SAMN52817386 |
| W933a4b4 | NS | 9 | 277 | 6128015 | 58 | 71733 | 54374 | 5655 | SAMN52817387 |
| W943a4b4 | NS | 9 | 275 | 6126873 | 58 | 68957 | 54625 | 5648 | SAMN52817388 |
| W953a4b4 | NS | 9 | 282 | 6150983 | 58 | 71405 | 53966 | 5669 | SAMN52817389 |
| W9613a5b4 | NS | 9 | 248 | 6054848 | 58.05 | 71871 | 19348 | 5544 | SAMN52817390 |
| W973a4b5 | NS | 9 | 283 | 6152131 | 57.99 | 71875 | 54251 | 5676 | SAMN52817391 |
| W9813a5b5 | NS | 9 | 252 | 6054453 | 58.05 | 71871 | 18711 | 5546 | SAMN52817392 |
| W983a4b5 | NS | 9 | 285 | 6158759 | 57.99 | 71871 | 53362 | 5694 | SAMN52817393 |
| W9913a5b5 | NS | 9 | 246 | 6055181 | 58.05 | 72303 | 18925 | 5546 | SAMN52817394 |
| W993a4b5 | NS | 9 | 293 | 6160241 | 57.99 | 71830 | 54035 | 5681 | SAMN52817395 |
| PccICMP5710 |  |  |  | 580000 | 58 | 39600 |  |  | SAMN03976288 |
| PsvNCPPPB3335 |  |  |  |  |  |  |  |  | SAMN02471367 |

| **Purpose** | **Name** | **Sequence (5' -> 3')** | **Length (bp)** | **Conditions** |
| --- | --- | --- | --- | --- |
| Final *Psf*-specific primers for characterisation of environmental isolates | nicB_6-F | CAGGTACGCCCTACAAGTGG | 386 | 95^o^C 5 mins,  35 cycles [95^o^C 30 secs, 59^o^C 30 secs, 72^o^C 30 secs]  72^o^C 5 mins |
|  | nicB_7-R | CGGAACCGTTCGTTTAGCAC |  |  |
| Allele swap of mutated GacA into mobile strain via pk18mobsac | gacA_23.13_EcoRI-F | CCGGAGAATTCAGACGCTTCTTGAGGTTC | 874 | 98°C 30 secs  35 cycles [98°C 10 secs, 69°C 30 secs, 72°C 20 secs/kb]  72^o^C 5 mins |
|  | gacA_23.13_BamHI-R | CCAGGATCCGTCAAGCTGTCGAGAGTC |  |  |
| Allele swap of mutated GacS into mobile strain via pk18mobsac | gacS_15.13_EcoRI-F | CCAGAATTCGGCCTACAAACCATCGTG | 924 |  |
|  | gacS_15.13_BamHI-R | CCAGGATCCCAACGCCTGTACATCCTG |  |  |
| Clone GacA ORF into pBBR1MSC-5 expression vector | GacA_16.13_SpeI-F | GGCCAAACTAGTGGTGTTGGGCAAAGATG | 887 |  |
|  | GacA_16.13_ApaI-R | GGCCAAGGGCCCCGAAAGTCTGGGTCATG |  |  |
| Clone GacS ORF into pBBR1MSC-5 expression vector | GacS_16.13_SpeI-F | GGCCAAACTAGTCTGCACAGAGAGTTAGG | 2,907 |  |
|  | GacS_16.13_ApaI-R | GGCCAAGGGCCCGAGTGTGAGGAGCGAAC |  |  |
| Quantifying plasmid insert size | M13F | TGTAAAACGACGGCCAGT | NA | 95^o^C 5 mins,  35 cycles [95^o^C 30 secs, 55^o^C 30 secs, 72^o^C 1 mins 10 secs]  72^o^C 5 mins |
|  | M13R | CAGGAAACAGCTATGACC |  |  |

**Table S2 PCR conditions and primers used in this study.**

**Table S3 Distribution of SNP types across core genomic regions and their effect on protein function.**

|  | Total | Intergenic | Missense | Synonymous | Stop gained | Stop retained | Stop lost | Start lost | Non coding transcript |
| --- | --- | --- | --- | --- | --- | --- | --- | --- | --- |
| Unmasked SNPs | 833 | 124 | 469 | 216 | 18 | 1 | 3 | 1 | 1 |
| MGE SNPs | 17 | 0 | 10 | 6 | 1 | 0 | 0 | 0 | 0 |
| Recombinant SNPs | 8 | 1 | 6 | 1 | 0 | 0 | 0 | 0 | 0 |
| All SNPs | 858 | 125 | 485 | 223 | 19 | 1 | 3 | 1 | 1 |

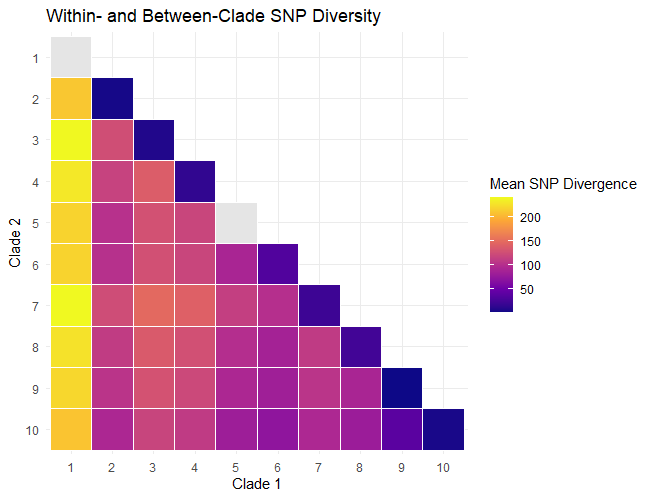

Mean SNP Distance

Figure S3 Within- and Between-Clade SNP diversity. A SNP matrix was generated from *Psf* genome alignments with reference W163. Strains were grouped into their respective clade and the mean pairwise distance between isolates was calculated using the upper triangle of the matrix, to avoid redundant pairwise comparisons. Between clade diversity was calculated between all possible pairs. For each clade pair, corresponding strains were extracted and used to subset the SNP distance matrix. Mean pairwise distance (clade divergence) between all isolates in the two clades were calculated.

Figure S4 Correlation between genetic distance and geographic separation. A) The Mantel test based on Pearson's product-moment correlation was applied to two distance matrices of genetic distance, calculated by pairwise SNP distance, and geographic distance, based on latitude and longitude metadata for individual trees. B) Distances were defined by bins of 100km. The Levene test was used to test whether there is equal variance in genetic distance between bins (p < 0.05).

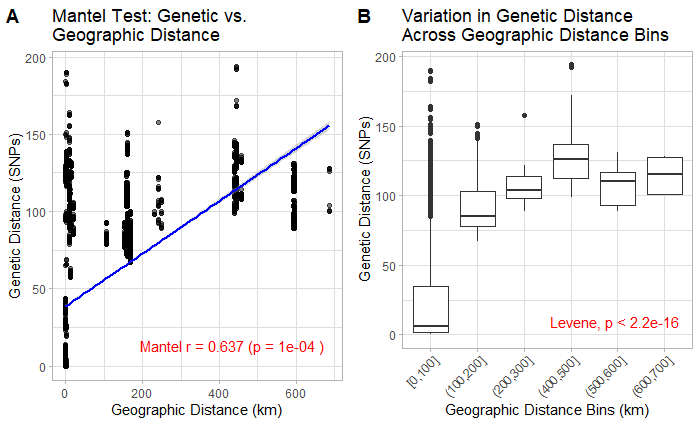

Table S4 SNP distance at three levels: Site, Tree, Tissue.

| Site | Tree | Tissue | Intra site diversity | Intra tree diversity | Intra tissue diversity |
| --- | --- | --- | --- | --- | --- |
| Cressbrook Dale | A1 | B2 | 23.128 | 14.476 | 1 |
| Cressbrook Dale | A1 | B2 | 23.128 | 14.476 | NA |
| Cressbrook Dale | A1 | B3 | 23.128 | 14.476 | NA |
| Cressbrook Dale | A1 | S1 | 23.128 | 14.476 | NA |
| Cressbrook Dale | A1 | S2 | 23.128 | 14.476 | 4 |
| Cressbrook Dale | A1 | S3 | 23.128 | 14.476 | NA |
| Cressbrook Dale | A3 | B1 | 23.128 | 16.667 | 5 |
| Cressbrook Dale | A3 | B3 | 23.128 | 16.667 | NA |
| Cressbrook Dale | A3 | S3 | 23.128 | 16.667 | NA |
| Cressbrook Dale | A4 | B2 | 23.128 | 1 | NA |
| Cressbrook Dale | A4 | B3 | 23.128 | 1 | NA |
| Isle of Mull | A1 | B1 | 73.571 | 73.571 | 0.6667 |
| Isle of Mull | A1 | B2 | 73.571 | 73.571 | 3 |
| Isle of Mull | A1 | B3 | 73.571 | 73.571 | NA |
| Isle of Mull | A1 | B4 | 73.571 | 73.571 | 4 |
| Isle of Mull | A1 | B5 | 73.571 | 73.571 | NA |
| Lathkill Dale | A1 | B3 | 90.189 | 2 | NA |
| Lathkill Dale | A1 | B4 | 90.189 | 2 | 3.3333 |
| Lathkill Dale | A1 | B5 | 90.189 | 2 | NA |
| Lathkill Dale | A1 | B6 | 90.189 | 2 | 1 |
| Lathkill Dale | A2 | B6 | 90.189 |  | NA |
| Lathkill Dale | A3 | B4 | 90.189 | 1 | NA |
| Lathkill Dale | A3 | B5 | 90.189 | 1 | NA |
| Lathkill Dale | A3 | B6 | 90.189 | 1 | NA |
| Lathkill Dale | A4 | B4 | 90.189 | 1 | NA |
| Lathkill Dale | A4 | B5 | 90.189 | 1 | NA |
| Lathkill Dale | A5 | 1B | 90.189 | 14.214 | NA |
| Lathkill Dale | A5 | B1 | 90.189 | 14.214 | 0.5 |
| Lathkill Dale | A5 | B2 | 90.189 | 14.214 | 0.5 |
| Lathkill Dale | A5 | B3 | 90.189 | 14.214 | NA |
| Lathkill Dale | A6 | B4 | 90.189 | NA | NA |
| Lathkill Dale | A6 | B5 | 90.189 | NA | NA |
| Lathkill Dale | A7 | B2 | 90.189 | NA | NA |
| Marden park | A1 | B1 | NA | NA | NA |
| Matlock | A1 | B1 | 1 | NA | NA |
| Matlock | A2 | B1 | 1 | 0 | 0 |
| Wytham Woods | A1 | B1 | 8.3758 | 1.7368 | 4.8667 |
| Wytham Woods | A1 | B2 | 8.3758 | 1.7368 | 0.25 |
| Wytham Woods | A1 | B3 | 8.3758 | 1.7368 | 0 |
| Wytham Woods | A2 | B4 | 8.3758 | NA | NA |
| Wytham Woods | A3 | B1 | 8.3758 | 18.036 | NA |
| Wytham Woods | A3 | B2 | 8.3758 | 18.036 | 0.6667 |
| Wytham Woods | A3 | B4 | 8.3758 | 18.036 | 2.5 |
| Wytham Woods | A3 | B5 | 8.3758 | 18.036 | NA |
| Wytham Woods | A3 | B6 | 8.3758 | 18.036 | NA |
| Wytham Woods | A4 | B1 | 8.3758 | 5.6417 | NA |
| Wytham Woods | A4 | B2 | 8.3758 | 5.6417 | 2 |
| Wytham Woods | A4 | B3 | 8.3758 | 5.6417 | 0 |
| Wytham Woods | A4 | B4 | 8.3758 | 5.6417 | 1.3333 |
| Wytham Woods | A4 | B5 | 8.3758 | 5.6417 | 0 |
| Wytham Woods | A4 | B6 | 8.3758 | 5.6417 | 0 |
| Wytham Woods | A5 | B1 | 8.3758 | 2.5735 | 1.8333 |
| Wytham Woods | A5 | B3 | 8.3758 | 2.5735 | 0 |
| Wytham Woods | A5 | B4 | 8.3758 | 2.5735 | 2.6667 |
| Wytham Woods | A5 | B5 | 8.3758 | 2.5735 | 0 |
| Wytham Woods | A5 | B6 | 8.3758 | 2.5735 | 4.6667 |
| Wytham Woods | A7 | B2 | 8.3758 | 18.036 | 15.2 |
| Wytham Woods | A7 | B3 | 8.3758 | 18.036 | 26.667 |
| Wytham Woods | A9 | B3 | 8.3758 | NA | NA |
| Wytham Woods | A11 | B1 | 8.3758 | 26 | 26 |
| Wytham Woods | A12 | B2 | 8.3758 | NA | NA |

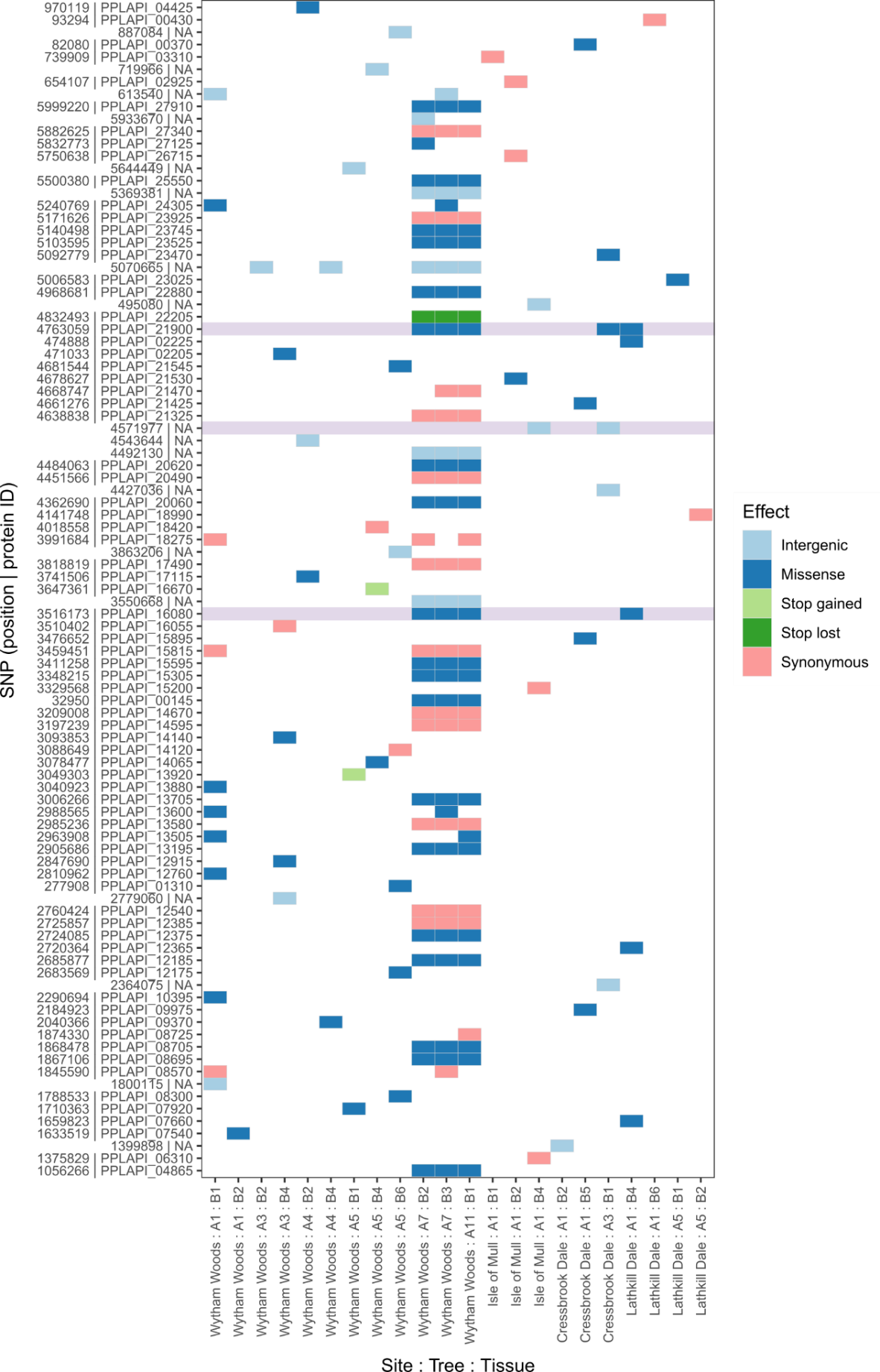

**Figure S5 Within-tissue allele distribution.** SNPs occurring between strains isolated from the same tissue sample were extracted. Within-tissue SNPs are shown by coloured blocks, including intergenic (light blue), missense (dark blue), stop gained (light green), stop lost (dark green) and synonymous (pink). Alleles shared across multiple sites are highlighted light purple. Protein IDs correspond to annotated CDS from reference W163 (SAMN52659773).

**Table S5 Genes with multiple non-synonymous and synonymous mutations**

| W163 bakta | Count | Mutation type | Effect | SNP information | Location | Gene function | Gene |
| --- | --- | --- | --- | --- | --- | --- | --- |
| PPLAPI_00805 | 2 | synonymous_variant | LOW | c.1356G>A | 1356/2805 | DUF2339 domain-containing protein | NA |
| PPLAPI_00805 | 2 | missense_variant | MODERATE | c.203G>T | 203/2805 | DUF2339 domain-containing protein | NA |
| PPLAPI_01150 | 2 | synonymous_variant | LOW | c.720G>A | 720/789 | zinc ABC transporter permease subunit ZnuB | znuB |
| PPLAPI_01150 | 2 | missense_variant | MODERATE | c.190T>C | 190/789 | zinc ABC transporter permease subunit ZnuB | znuB |
| PPLAPI_01185 | 3 | missense_variant | MODERATE | c.61G>T | 61/2766 | DNA polymerase I | polA |
| PPLAPI_01185 | 3 | missense_variant | MODERATE | c.893A>G | 893/2766 | DNA polymerase I | polA |
| PPLAPI_01185 | 3 | missense_variant | MODERATE | c.2285G>A | 2285/2766 | DNA polymerase I | polA |
| PPLAPI_01600 | 2 | missense_variant | MODERATE | c.388G>T | 388/801 | twin-arginine translocase subunit TatC | tatC |
| PPLAPI_01600 | 2 | missense_variant | MODERATE | c.136A>G | 136/801 | twin-arginine translocase subunit TatC | tatC |
| PPLAPI_01630 | 2 | missense_variant | MODERATE | c.258G>T | 258/624 | Ubiquinone biosynthesis protein UbiJ%2C contains SCP2 domain | ubiJ |
| PPLAPI_01630 | 2 | missense_variant | MODERATE | c.58C>A | 58/624 | Ubiquinone biosynthesis protein UbiJ%2C contains SCP2 domain | ubiJ |
| PPLAPI_02280 | 2 | synonymous_variant | LOW | c.1029G>A | 1029/3954 | trifunctional transcriptional regulator/proline dehydrogenase/L-glutamate gamma-semialdehyde dehydrogenase | putA |
| PPLAPI_02280 | 2 | missense_variant | MODERATE | c.20G>C | 20/3954 | trifunctional transcriptional regulator/proline dehydrogenase/L-glutamate gamma-semialdehyde dehydrogenase | putA |
| PPLAPI_02645 | 2 | stop_gained | HIGH | c.14T>G | 14/1935 | Methyl-accepting chemotaxis protein (MCP) | tar |
| PPLAPI_02645 | 2 | synonymous_variant | LOW | c.1239T>C | 1239/1935 | Methyl-accepting chemotaxis protein (MCP) | tar |
| PPLAPI_02700 | 2 | missense_variant | MODERATE | c.2398G>A | 2398/2745 | Permease of the major facilitator superfamily | NA |
| PPLAPI_02700 | 2 | missense_variant | MODERATE | c.239T>A | 239/2745 | Permease of the major facilitator superfamily | NA |
| PPLAPI_03130-PPLAPI_03135 | 2 | intergenic_region | MODIFIER | n.695247A>G | NA | NA | NA |
| PPLAPI_03130-PPLAPI_03135 | 2 | intergenic_region | MODIFIER | n.695546T>C | NA | NA | NA |
| PPLAPI_03370 | 2 | missense_variant | MODERATE | c.559C>A | 559/948 | 4-hydroxy-3-methylbut-2-enyl diphosphate reductase | ispH |
| PPLAPI_03370 | 2 | synonymous_variant | LOW | c.861C>A | 861/948 | 4-hydroxy-3-methylbut-2-enyl diphosphate reductase | ispH |
| PPLAPI_03710 | 2 | missense_variant | MODERATE | c.310G>C | 310/2436 | putative ATP-dependent protease | lonB |
| PPLAPI_03710 | 2 | missense_variant | MODERATE | c.311A>G | 311/2436 | putative ATP-dependent protease | lonB |
| PPLAPI_03825 | 4 | missense_variant | MODERATE | c.785A>C | 785/1596 | Solute-binding protein family 5 domain-containing protein | NA |
| PPLAPI_03825 | 4 | missense_variant | MODERATE | c.503C>T | 503/1596 | Solute-binding protein family 5 domain-containing protein | NA |
| PPLAPI_03825 | 4 | missense_variant | MODERATE | c.419G>A | 419/1596 | Solute-binding protein family 5 domain-containing protein | NA |
| PPLAPI_03825 | 4 | missense_variant | MODERATE | c.92A>T | 92/1596 | Solute-binding protein family 5 domain-containing protein | NA |
| PPLAPI_04360 | 2 | synonymous_variant | LOW | c.2439C>A | 2439/3816 | YhdP family protein | yhdR |
| PPLAPI_04360 | 2 | missense_variant | MODERATE | c.2452G>C | 2452/3816 | YhdP family protein | yhdR |
| PPLAPI_04520 | 2 | synonymous_variant | LOW | c.159T>C | 159/759 | Nif3-like dinuclear metal center hexameric protein | nIF3 |
| PPLAPI_04520 | 2 | synonymous_variant | LOW | c.393G>T | 393/759 | Nif3-like dinuclear metal center hexameric protein | nIF3 |
| PPLAPI_04540 | 2 | missense_variant | MODERATE | c.227G>C | 227/636 | Alpha/beta superfamily hydrolase | NA |
| PPLAPI_04540 | 2 | synonymous_variant | LOW | c.228T>C | 228/636 | Alpha/beta superfamily hydrolase | NA |
| PPLAPI_04865 | 2 | missense_variant | MODERATE | c.1618A>G | 1618/2481 | Toxin VasX N-terminal region domain-containing protein | vasX |
| PPLAPI_04865 | 2 | missense_variant | MODERATE | c.1489C>A | 1489/2481 | Toxin VasX N-terminal region domain-containing protein | vasX |
| PPLAPI_05330 | 3 | synonymous_variant | LOW | c.630G>T | 630/16536 | Calcium binding haemolysin protein | NA |
| PPLAPI_05330 | 3 | missense_variant | MODERATE | c.7273T>A | 7273/16536 | Calcium binding haemolysin protein | NA |
| PPLAPI_05330 | 3 | synonymous_variant | LOW | c.9948G>T | 9948/16536 | Calcium binding haemolysin protein | NA |
| PPLAPI_05510 | 2 | synonymous_variant | LOW | c.802T>C | 802/1179 | Acetyl-CoA acetyltransferase | paaJ |
| PPLAPI_05510 | 2 | missense_variant | MODERATE | c.501G>T | 501/1179 | Acetyl-CoA acetyltransferase | paaJ |
| PPLAPI_05510-PPLAPI_05515 | 2 | intergenic_region | MODIFIER | n.1202690C>A | NA | NA | NA |
| PPLAPI_05510-PPLAPI_05515 | 2 | intergenic_region | MODIFIER | n.1202695G>T | NA | NA | NA |
| PPLAPI_05690 | 2 | stop_gained | HIGH | c.1174G>T | 1174/1248 | mannuronan 5-epimerase | NA |
| PPLAPI_05690 | 2 | missense_variant | MODERATE | c.109G>A | 109/1248 | mannuronan 5-epimerase | NA |
| PPLAPI_05765 | 2 | synonymous_variant | LOW | c.1803C>T | 1803/2010 | PhoD-like phosphatase metallophosphatase domain-containing protein | NA |
| PPLAPI_05765 | 2 | synonymous_variant | LOW | c.1212C>A | 1212/2010 | PhoD-like phosphatase metallophosphatase domain-containing protein | NA |
| PPLAPI_05990 | 2 | missense_variant | MODERATE | c.901T>A | 901/1026 | TRAP-type transport system%2C periplasmic component%2C putative N-acetylneuraminate-binding protein | NA |
| PPLAPI_05990 | 2 | synonymous_variant | LOW | c.39G>T | 39/1026 | TRAP-type transport system%2C periplasmic component%2C putative N-acetylneuraminate-binding protein | NA |
| PPLAPI_07245 | 2 | missense_variant | MODERATE | c.913C>T | 913/1167 | ABC-type sugar transport system%2C ATPase component MalK | malK |
| PPLAPI_07245 | 2 | missense_variant | MODERATE | c.1127G>C | 1127/1167 | ABC-type sugar transport system%2C ATPase component MalK | malK |
| PPLAPI_07690 | 2 | missense_variant | MODERATE | c.3409G>A | 3409/5157 | Type III effector AvrE1 | avrE1 |
| PPLAPI_07690 | 2 | missense_variant | MODERATE | c.2392T>G | 2392/5157 | Type III effector AvrE1 | avrE1 |
| PPLAPI_07825-PPLAPI_07830 | 2 | intergenic_region | MODIFIER | n.1693589T>C | NA | NA | NA |
| PPLAPI_07825-PPLAPI_07830 | 2 | intergenic_region | MODIFIER | n.1693706A>G | NA | NA | NA |
| PPLAPI_08555 | 2 | missense_variant | MODERATE | c.670G>A | 670/2217 | DNA internalization-related competence protein ComEC/Rec2 | NA |
| PPLAPI_08555 | 2 | missense_variant | MODERATE | c.1537A>G | 1537/2217 | DNA internalization-related competence protein ComEC/Rec2 | NA |
| PPLAPI_09055 | 2 | missense_variant | MODERATE | c.956C>A | 956/1158 | DSBA oxidoreductase | NA |
| PPLAPI_09055 | 2 | missense_variant | MODERATE | c.1080A>T | 1080/1158 | DSBA oxidoreductase | NA |
| PPLAPI_09195 | 2 | missense_variant | MODERATE | c.1805C>G | 1805/6648 | Non-ribosomal peptide synthetase module | NA |
| PPLAPI_09195 | 2 | missense_variant | MODERATE | c.5077G>T | 5077/6648 | Non-ribosomal peptide synthetase module | NA |
| PPLAPI_09370 | 2 | missense_variant | MODERATE | c.850C>T | 850/1686 | Phospholipase C | acpA |
| PPLAPI_09370 | 2 | stop_gained | HIGH | c.566G>A | 566/1686 | Phospholipase C | acpA |
| PPLAPI_09975 | 2 | missense_variant | MODERATE | c.1840C>A | 1840/8628 | Carrier domain-containing protein | NA |
| PPLAPI_09975 | 2 | missense_variant | MODERATE | c.2602C>A | 2602/8628 | Carrier domain-containing protein | NA |
| PPLAPI_09985 | 2 | synonymous_variant | LOW | c.525G>T | 525/2232 | Outer membrane receptor for ferric coprogen and ferric-rhodotorulic acid | fhuE |
| PPLAPI_09985 | 2 | synonymous_variant | LOW | c.354T>A | 354/2232 | Outer membrane receptor for ferric coprogen and ferric-rhodotorulic acid | fhuE |
| PPLAPI_10595 | 2 | missense_variant | MODERATE | c.102T>G | 102/660 | bifunctional phosphoserine phosphatase/homoserine phosphotransferase ThrH | thrH |
| PPLAPI_10595 | 2 | missense_variant | MODERATE | c.103G>C | 103/660 | bifunctional phosphoserine phosphatase/homoserine phosphotransferase ThrH | thrH |
| PPLAPI_10945 | 2 | missense_variant | MODERATE | c.773A>C | 773/1029 | Fe2+-dicitrate sensor%2C membrane component | NA |
| PPLAPI_10945 | 2 | missense_variant | MODERATE | c.649G>C | 649/1029 | Fe2+-dicitrate sensor%2C membrane component | NA |
| PPLAPI_11020 | 2 | synonymous_variant | LOW | c.447C>T | 447/1026 | Inosine-uridine nucleoside N-ribohydrolase | uRH1 |
| PPLAPI_11020 | 2 | missense_variant | MODERATE | c.604C>T | 604/1026 | Inosine-uridine nucleoside N-ribohydrolase | uRH1 |
| PPLAPI_11240-PPLAPI_11245 | 2 | intergenic_region | MODIFIER | n.2464665C>T | NA | NA | NA |
| PPLAPI_11240-PPLAPI_11245 | 2 | intergenic_region | MODIFIER | n.2464666G>T | NA | NA | NA |
| PPLAPI_11860 | 2 | missense_variant | MODERATE | c.613T>C | 613/3096 | Error-prone DNA polymerase | dnaE2 |
| PPLAPI_11860 | 2 | missense_variant | MODERATE | c.1033T>A | 1033/3096 | Error-prone DNA polymerase | dnaE2 |
| PPLAPI_12590 | 3 | missense_variant | MODERATE | c.971A>T | 971/1371 | Glycine zipper family protein | NA |
| PPLAPI_12590 | 3 | missense_variant | MODERATE | c.794G>T | 794/1371 | Glycine zipper family protein | NA |
| PPLAPI_12590 | 3 | missense_variant | MODERATE | c.541T>A | 541/1371 | Glycine zipper family protein | NA |
| PPLAPI_12840 | 2 | missense_variant | MODERATE | c.811G>T | 811/1419 | Major facilitator family transporter | NA |
| PPLAPI_12840 | 2 | missense_variant | MODERATE | c.668C>A | 668/1419 | Major facilitator family transporter | NA |
| PPLAPI_12895 | 2 | missense_variant | MODERATE | c.410A>T | 410/1890 | penicillin-binding protein 2 | mrdA |
| PPLAPI_12895 | 2 | synonymous_variant | LOW | c.969A>C | 969/1890 | penicillin-binding protein 2 | mrdA |
| PPLAPI_13005 | 2 | missense_variant | MODERATE | c.43G>A | 43/1623 | Methyl-accepting chemotaxis protein CtpH | ctpH |
| PPLAPI_13005 | 2 | synonymous_variant | LOW | c.996G>T | 996/1623 | Methyl-accepting chemotaxis protein CtpH | ctpH |
| PPLAPI_13160 | 2 | synonymous_variant | LOW | c.789T>G | 789/2175 | type VI secretion system ATPase TssH | tssH |
| PPLAPI_13160 | 2 | missense_variant | MODERATE | c.1979G>T | 1979/2175 | type VI secretion system ATPase TssH | tssH |
| PPLAPI_13580 | 2 | missense_variant | MODERATE | c.592T>C | 592/879 | DNA-binding transcriptional regulator%2C LysR family | lysR |
| PPLAPI_13580 | 2 | synonymous_variant | LOW | c.723T>C | 723/879 | DNA-binding transcriptional regulator%2C LysR family | lysR |
| PPLAPI_13620 | 2 | missense_variant | MODERATE | c.944A>T | 944/1110 | bifunctional 3%2C4-dihydroxy-2-butanone-4-phosphate synthase/GTP cyclohydrolase II | ribBA |
| PPLAPI_13620 | 2 | missense_variant | MODERATE | c.92G>T | 92/1110 | bifunctional 3%2C4-dihydroxy-2-butanone-4-phosphate synthase/GTP cyclohydrolase II | ribBA |
| PPLAPI_13935 | 2 | missense_variant | MODERATE | c.490A>T | 490/693 | Chromosome segregation ATPase | NA |
| PPLAPI_13935 | 2 | synonymous_variant | LOW | c.478C>A | 478/693 | Chromosome segregation ATPase | NA |
| PPLAPI_14400 | 2 | missense_variant | MODERATE | c.490T>G | 490/1215 | SfnB family sulfur acquisition oxidoreductase | sfnB |
| PPLAPI_14400 | 2 | synonymous_variant | LOW | c.181C>T | 181/1215 | SfnB family sulfur acquisition oxidoreductase | sfnB |
| PPLAPI_14625 | 2 | missense_variant | MODERATE | c.206C>A | 206/588 | UvrY/SirA/GacA family response regulator transcription factor | uvrY |
| PPLAPI_14625 | 2 | missense_variant | MODERATE | c.11A>G | 11/588 | UvrY/SirA/GacA family response regulator transcription factor | uvrY |
| PPLAPI_14670 | 2 | missense_variant | MODERATE | c.44A>G | 44/1026 | ABC-type microcin C transport system%2C permease component YejE | yejE |
| PPLAPI_14670 | 2 | synonymous_variant | LOW | c.531G>C | 531/1026 | ABC-type microcin C transport system%2C permease component YejE | yejE |
| PPLAPI_14975 | 2 | missense_variant | MODERATE | c.1822G>A | 1822/2784 | malto-oligosyltrehalose synthase | treY |
| PPLAPI_14975 | 2 | missense_variant | MODERATE | c.1835C>A | 1835/2784 | malto-oligosyltrehalose synthase | treY |
| PPLAPI_15925 | 2 | missense_variant | MODERATE | c.848A>T | 848/2250 | DEAD/DEAH box helicase | NA |
| PPLAPI_15925 | 2 | missense_variant | MODERATE | c.1825G>A | 1825/2250 | DEAD/DEAH box helicase | NA |
| PPLAPI_17120 | 2 | synonymous_variant | LOW | c.30C>A | 30/615 | transporter | NA |
| PPLAPI_17120 | 2 | missense_variant | MODERATE | c.29C>A | 29/615 | transporter | NA |
| PPLAPI_17395 | 4 | missense_variant | MODERATE | c.53G>T | 53/735 | cAMP-binding domain of CRP or a regulatory subunit of cAMP-dependent protein kinases | crp |
| PPLAPI_17395 | 4 | synonymous_variant | LOW | c.381G>A | 381/735 | cAMP-binding domain of CRP or a regulatory subunit of cAMP-dependent protein kinases | crp |
| PPLAPI_17395 | 4 | missense_variant | MODERATE | c.482C>T | 482/735 | cAMP-binding domain of CRP or a regulatory subunit of cAMP-dependent protein kinases | crp |
| PPLAPI_17395 | 4 | synonymous_variant | LOW | c.498C>A | 498/735 | cAMP-binding domain of CRP or a regulatory subunit of cAMP-dependent protein kinases | crp |
| PPLAPI_20830 | 2 | missense_variant | MODERATE | c.344G>T | 344/801 | Putative NADPH-quinone reductase (modulator of drug activity B) | mdaB |
| PPLAPI_20830 | 2 | missense_variant | MODERATE | c.475C>A | 475/801 | Putative NADPH-quinone reductase (modulator of drug activity B) | mdaB |
| PPLAPI_21000 | 2 | missense_variant | MODERATE | c.322C>G | 322/1833 | ABC-type multidrug transport system%2C ATPase and permease component | mdlB |
| PPLAPI_21000 | 2 | missense_variant | MODERATE | c.776G>T | 776/1833 | ABC-type multidrug transport system%2C ATPase and permease component | mdlB |
| PPLAPI_21020 | 2 | missense_variant | MODERATE | c.3388T>C | 3388/3909 | ATP-dependent RNA helicase HrpA | hrpA |
| PPLAPI_21020 | 2 | missense_variant | MODERATE | c.1375T>C | 1375/3909 | ATP-dependent RNA helicase HrpA | hrpA |
| PPLAPI_21405 | 2 | missense_variant | MODERATE | c.92G>T | 92/747 | ABC-type amino acid transport system%2C permease component | hisM |
| PPLAPI_21405 | 2 | missense_variant | MODERATE | c.93G>T | 93/747 | ABC-type amino acid transport system%2C permease component | hisM |
| PPLAPI_21530 | 2 | missense_variant | MODERATE | c.284A>G | 284/768 | NADPH-dependent ferric siderophore reductase%2C contains FAD-binding and SIP domains | viuB |
| PPLAPI_21530 | 2 | missense_variant | MODERATE | c.718G>A | 718/768 | NADPH-dependent ferric siderophore reductase%2C contains FAD-binding and SIP domains | viuB |
| PPLAPI_21690 | 2 | missense_variant | MODERATE | c.727T>C | 727/1617 | L-aspartate oxidase | nadB |
| PPLAPI_21690 | 2 | missense_variant | MODERATE | c.862G>A | 862/1617 | L-aspartate oxidase | nadB |
| PPLAPI_22145 | 2 | missense_variant | MODERATE | c.725A>G | 725/978 | GTPase | yjiA |
| PPLAPI_22145 | 2 | missense_variant | MODERATE | c.166G>C | 166/978 | GTPase | yjiA |
| PPLAPI_22200-PPLAPI_22205 | 2 | intergenic_region | MODIFIER | n.4832144C>T | NA | NA | NA |
| PPLAPI_22200-PPLAPI_22205 | 2 | intergenic_region | MODIFIER | n.4832286T>A | NA | NA | NA |
| PPLAPI_22655 | 2 | missense_variant | MODERATE | c.118C>T | 118/924 | Acetyl esterase/lipase | aes |
| PPLAPI_22655 | 2 | missense_variant | MODERATE | c.28C>G | 28/924 | Acetyl esterase/lipase | aes |
| PPLAPI_22955 | 2 | synonymous_variant | LOW | c.3324G>A | 3324/4950 | putative conserved protein YfaS%2C alpha-2-macroglobulin family | yfaS |
| PPLAPI_22955 | 2 | missense_variant | MODERATE | c.3958G>A | 3958/4950 | putative conserved protein YfaS%2C alpha-2-macroglobulin family | yfaS |
| PPLAPI_23820 | 3 | missense_variant | MODERATE | c.3313G>A | 3313/4074 | DNA-directed RNA polymerase subunit beta | rpoB |
| PPLAPI_23820 | 3 | missense_variant | MODERATE | c.1414G>T | 1414/4074 | DNA-directed RNA polymerase subunit beta | rpoB |
| PPLAPI_23820 | 3 | missense_variant | MODERATE | c.463C>T | 463/4074 | DNA-directed RNA polymerase subunit beta | rpoB |
| PPLAPI_24970-PPLAPI_24975 | 2 | intergenic_region | MODIFIER | n.5369381C>T | NA | NA | NA |
| PPLAPI_24970-PPLAPI_24975 | 2 | intergenic_region | MODIFIER | n.5369806G>T | NA | NA | NA |
| PPLAPI_25315 | 2 | missense_variant | MODERATE | c.1804C>A | 1804/2694 | Cyclic di-GMP metabolism protein%2C combines GGDEF and EAL domains with a 6TM membrane domain | NA |
| PPLAPI_25315 | 2 | stop_gained | HIGH | c.2034G>A | 2034/2694 | Cyclic di-GMP metabolism protein%2C combines GGDEF and EAL domains with a 6TM membrane domain | NA |
| PPLAPI_25365 | 2 | missense_variant | MODERATE | c.767C>A | 767/1986 | hypothetical protein | NA |
| PPLAPI_25365 | 2 | synonymous_variant | LOW | c.966C>T | 966/1986 | hypothetical protein | NA |
| PPLAPI_25550 | 2 | missense_variant | MODERATE | c.152C>T | 152/453 | NTF2 fold immunity protein domain-containing protein | NA |
| PPLAPI_25550 | 2 | missense_variant | MODERATE | c.315A>C | 315/453 | NTF2 fold immunity protein domain-containing protein | NA |
| PPLAPI_25595 | 2 | missense_variant | MODERATE | c.1293G>T | 1293/1314 | Outer membrane porin | NA |
| PPLAPI_25595 | 2 | missense_variant | MODERATE | c.353C>G | 353/1314 | Outer membrane porin | NA |
| PPLAPI_26275 | 2 | missense_variant | MODERATE | c.122G>A | 122/1818 | Signal transduction histidine kinase regulating C4-dicarboxylate transport system | NA |
| PPLAPI_26275 | 2 | missense_variant | MODERATE | c.1385G>A | 1385/1818 | Signal transduction histidine kinase regulating C4-dicarboxylate transport system | NA |
| PPLAPI_26515 | 2 | synonymous_variant | LOW | c.429G>A | 429/555 | (p)ppGpp synthase/hydrolase%2C HD superfamily | spoT |
| PPLAPI_26515 | 2 | missense_variant | MODERATE | c.508A>G | 508/555 | (p)ppGpp synthase/hydrolase%2C HD superfamily | spoT |
| PPLAPI_26650 | 2 | synonymous_variant | LOW | c.93A>G | 93/777 | Heme oxygenase-like protein | NA |
| PPLAPI_26650 | 2 | missense_variant | MODERATE | c.716T>G | 716/777 | Heme oxygenase-like protein | NA |

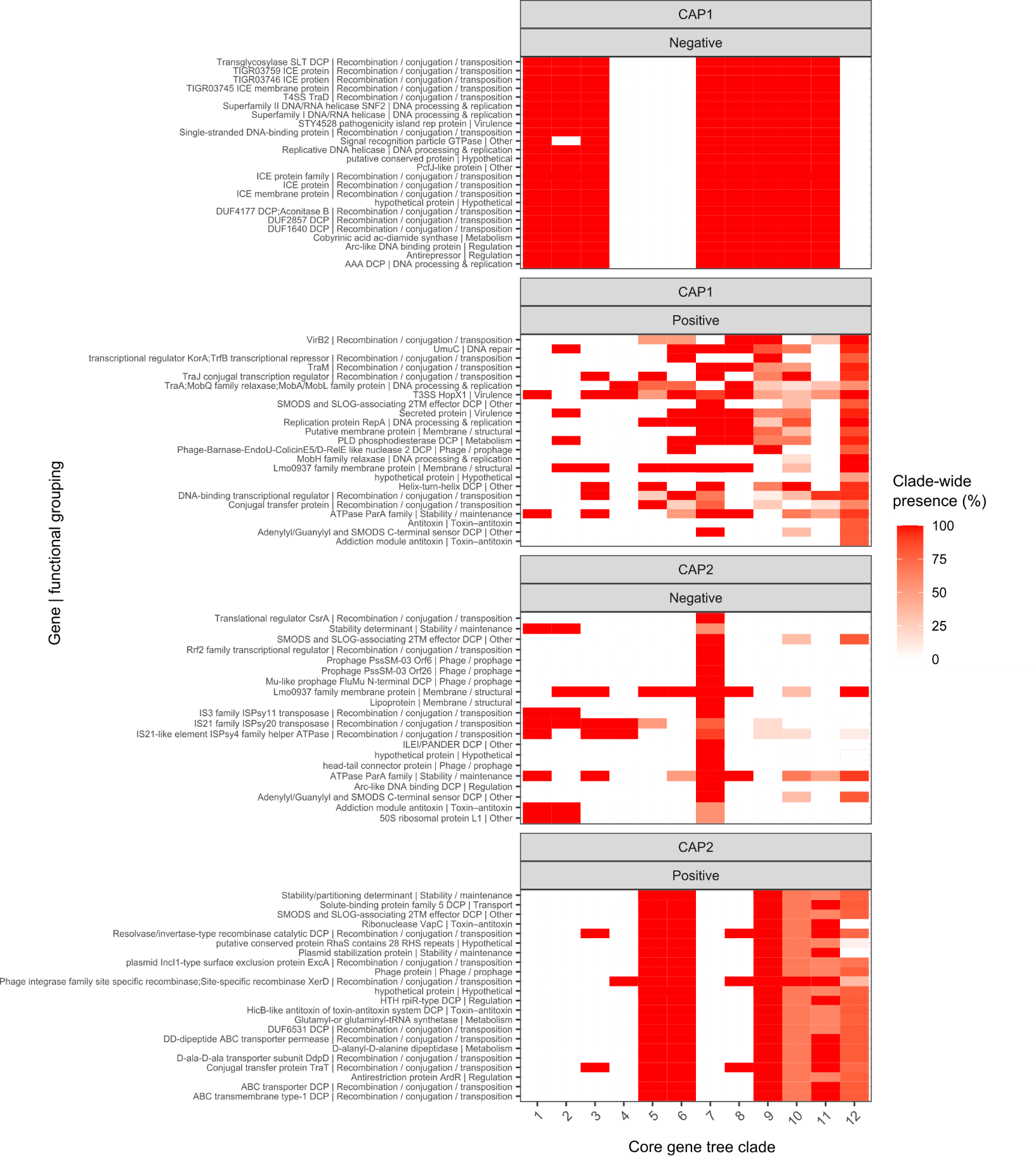

**Figure S6 Top 100 genes associated with accessory genome clustering.** Distance-based redundancy analysis (dbRDA) was performed on a binary gene presence/absence matrix to identify genes contributing most strongly to accessory genome variation. Gene scores (analogous to loadings in PCA) represent the contribution of each gene to variation along the first two dbRDA axes derived from a Jaccard distance matrix. Strains are grouped by phylogenetic clade (1–12), defined from the concatenated core gene phylogeny. Heatmaps show the proportion of strains within each clade containing a given gene (red, high prevalence; white, low prevalence). Genes shown correspond to the top positively loading genes for each axis. Functional categories were assigned manually based on annotation. Abbreviations: DCP, domain-containing protein; ICE, integrative and conjugative element.

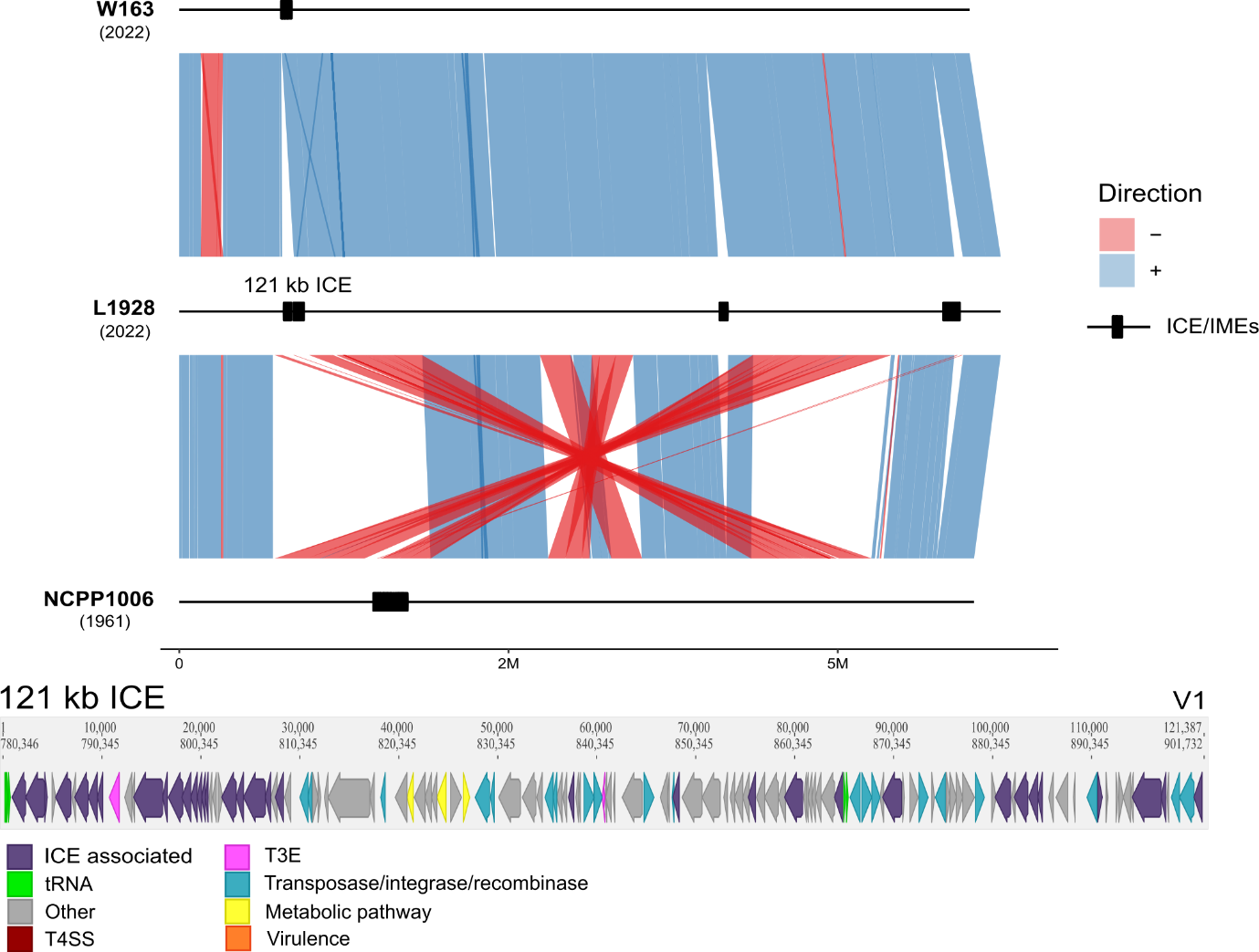

**Figure S7. Synteny plot of three *Psf* chromosomes and associated ICEs.** Three complete genomes—W163 (2022), L1928 (2022), and NCPPB 1006 (1961)—were aligned pairwise. Alignments were filtered to regions >10,000 bp to highlight structural variation. Genome backbones are shown as black lines, with putative integrative conjugative elements (ICEs) indicated by black squares, predicted by ICEfinder. Syntenic regions are shown as blue blocks (forward orientation) and red blocks (reverse orientation). Gene content within the 121 kb ICE is coloured by functional category: ICE-associated genes (purple; e.g. Tra, PilL, ParB, MobH, helicases, toxin–antitoxin systems), tRNA (green), hypothetical/unknown (grey), type III effectors (pink), mobile genetic elements (blue; transposases/integrases/recombinases), type IV secretion system (red), virulence genes (orange), and metabolic genes (yellow).

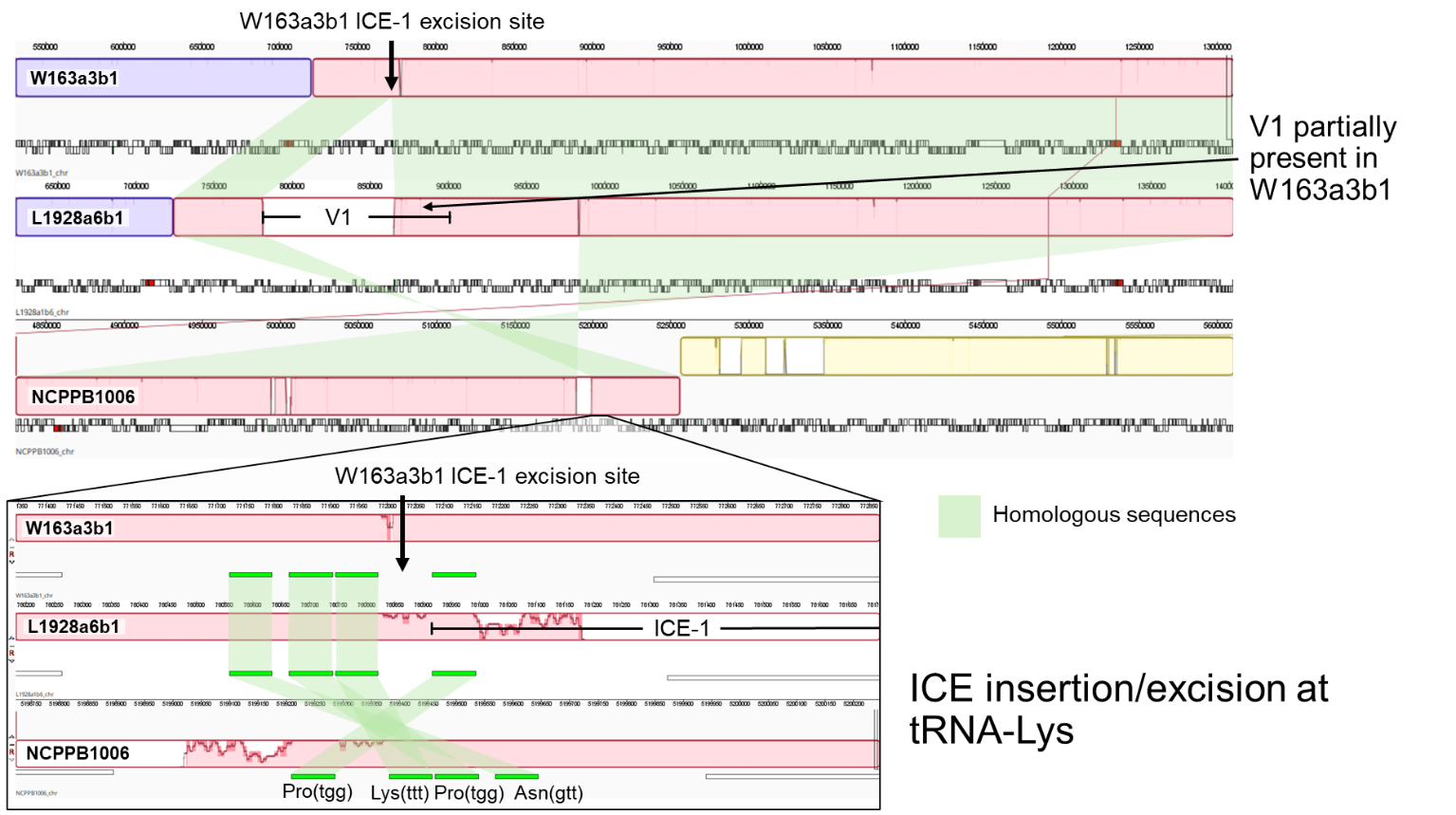

**Figure S8. Variable presence of an integrative conjugative element (ICE-V1) drives pangenome diversification.** Mauve alignment of genomes NCPPB 1006 (UK; 1961), L1928a6b1 (Lathkill Dale; 2022), and W163a3b1 (Wytham Woods; 2022). Syntenic chromosomal blocks are shown in different colours, with strain-specific regions in white and homologous regions indicated by green translucent connectors between genomes. L1928a6b1 contains a 121,387 bp putative integrative conjugative element (ICE-V1; labelled within a red chromosomal block). In contrast, W163a3b1 shows partial presence of this element, while NCPPB 1006 lacks it entirely.

A zoomed region highlights the ICE integration site at a tRNA-Lys locus (arrow). In L1928a6b1, the ICE is located adjacent to this site, whereas both W163a3b1 and NCPPB 1006 retain the tRNA-Lys locus but lack the full ICE sequence. The ICE also contains an internal tRNA-Lys site, which likely mediates excision in W163a3b1, resulting in its partial retention.

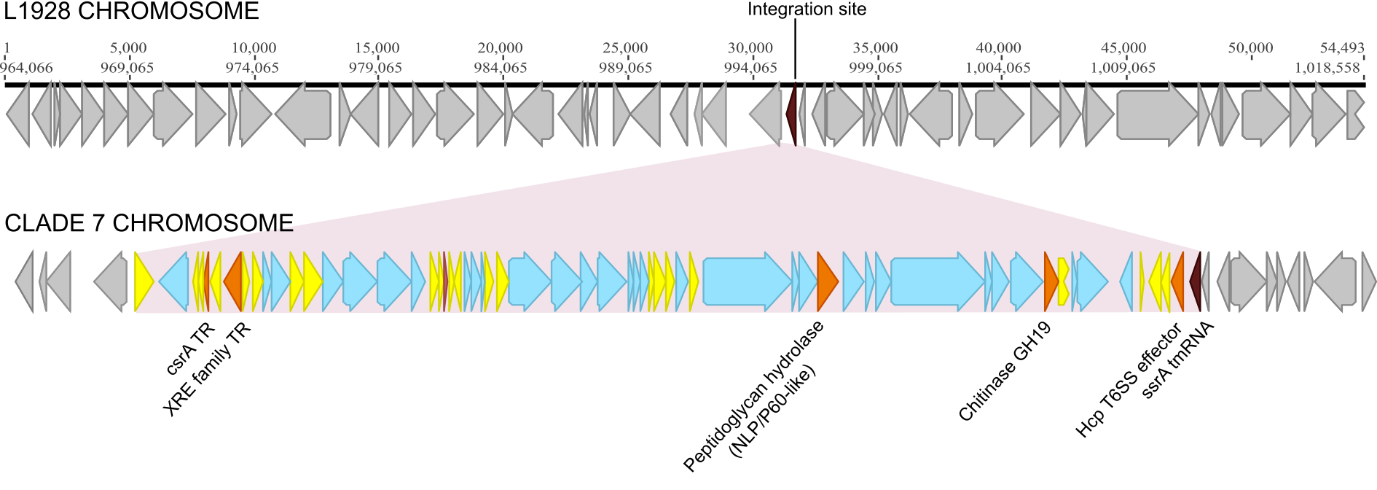

**Figure S9 Chromosomally integrated lysogenic prophage is exclusively present in clade 7 Lathkill Dale strains.** A lysogenic prophage (highlighted in pink) was integrated at a srrA tmRNA site (dark red) in reference genome L1928 (clade 9, Lathkill Dale). The prophage includes 31 prophage-associated genes (blue), five virulence factors (orange), and 22 with other functions (yellow). The virulence factors include two transcriptional regulators, CsrA and XRE; denoted TR. Grey genes represent those present in the chromosome of reference L1928 (top) and the chromosome of the clade 7 strain L777 (bottom).

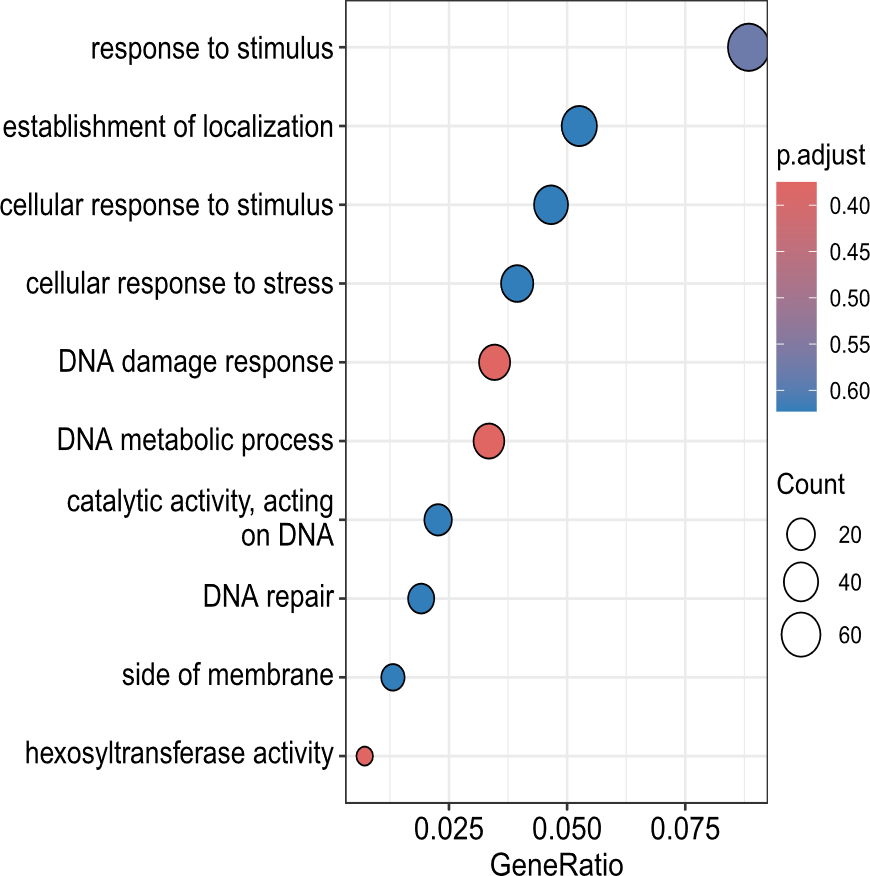
**Figure S10** GO pathways disproportionately affected by mutations, including SNPs, MNPs, indels, and complex variants*,* across 124 *Psf g*enomes. Significant enrichment is indicated by an asterisk (*), based on Fisher’s exact test (p < 0.05) and Benjamini–Hochberg false discovery rate (FDR) correction (q < 0.1).

**Table S6 KEGG enrichment genes**

| **KEGG pathway** | **Annotation** | **Gene** | **Gene Function** | **Type** | **Effect** | **Strain count** |
| --- | --- | --- | --- | --- | --- | --- |
| beta-Lactam resistance | PPLAPI_11990 | *dppD* | ABC-type dipeptide/oligopeptide/nickel transport system%2C ATPase component | snp | missense_variant c.422G>T p.Arg141Leu | 4 |
| beta-Lactam resistance | PPLAPI_12615 | *dppD* | ABC-type dipeptide/oligopeptide/nickel transport system%2C ATPase component | snp | missense_variant c.875G>A p.Cys292Tyr | 1 |
| beta-Lactam resistance | PPLAPI_12895 | *mrdA* | penicillin-binding protein 2 | snp | missense_variant c.410A>T p.Tyr137Phe | 1 |
| beta-Lactam resistance | PPLAPI_12895 | *mrdA* | penicillin-binding protein 2 | snp | synonymous_variant c.969A>C p.Ala323Ala | 1 |
| beta-Lactam resistance | PPLAPI_14365 | *adeC* | Outer membrane protein | snp | missense_variant c.1045G>A p.Val349Met | 4 |
| beta-Lactam resistance | PPLAPI_14370 | *adeB* | Multidrug efflux pump subunit AcrB | del | conservative_inframe_deletion c.1654_1659delCTGGTG p.Leu552_Val553del | 1 |
| beta-Lactam resistance | PPLAPI_14375 | *adeA* | Multidrug efflux pump subunit AcrA (membrane-fusion protein) | snp | missense_variant c.520G>A p.Ala174Thr | 1 |
| beta-Lactam resistance | PPLAPI_15340 | *tolC* | Outer membrane protein TolC | snp | missense_variant c.92C>A p.Pro31Gln | 1 |
| beta-Lactam resistance | PPLAPI_15365 | NA | Acriflavin resistance protein | snp | synonymous_variant c.1713A>G p.Thr571Thr | 1 |
| beta-Lactam resistance | PPLAPI_15380 | *nodT* | RND efflux system%2C outer membrane lipoprotein%2C NodT | snp | missense_variant c.766C>T p.His256Tyr | 6 |
| beta-Lactam resistance | PPLAPI_15385 | *ompR* | DNA-binding response regulator%2C OmpR family%2C contains REC and winged-helix (wHTH) domain | snp | synonymous_variant c.652T>C p.Leu218Leu | 1 |
| beta-Lactam resistance | PPLAPI_15395 | *baeS* | Signal transduction histidine kinase | snp | missense_variant c.451T>C p.Trp151Arg | 1 |
| beta-Lactam resistance | PPLAPI_15740 | *acrB* | Multidrug efflux pump subunit AcrB | snp | synonymous_variant c.1743G>A p.Lys581Lys | 3 |
| beta-Lactam resistance | PPLAPI_01720 | *mrcA* | Membrane carboxypeptidase/penicillin-binding protein | snp | missense_variant c.794T>C p.Ile265Thr | 4 |
| beta-Lactam resistance | PPLAPI_21520 | *mrcB* | Penicillin-binding protein 1B/1F%2C peptidoglycan transglycosylase/transpeptidase | del | frameshift_variant c.114_153delTTGGGTGATTGTGGTGCTGGCCTGCATTGCGCTGGGTGTT p.Phe38fs | 1 |
| beta-Lactam resistance | PPLAPI_21965 | *adeC* | outer membrane channel protein that acts as part of the AdeABC multidrug efflux pump | del | disruptive_inframe_deletion c.924_932delACGCGCAGC p.Arg309_Ala311del | 1 |
| beta-Lactam resistance | PPLAPI_25490 | NA | Sensor protein | snp | missense_variant c.994T>G p.Ser332Ala | 4 |
| beta-Lactam resistance | PPLAPI_00430 | *lysA* | diaminopimelate decarboxylase | snp | synonymous_variant c.909C>T p.Val303Val | 5 |
| Two-component system | PPLAPI_05390 | *terZ* | Stress response protein SCP2 | del | frameshift_variant c.758_759delAG p.Glu253fs | 4 |
| Two-component system | PPLAPI_05510 | *paaJ* | Acetyl-CoA acetyltransferase | snp | missense_variant c.501G>T p.Gln167His | 4 |
| Two-component system | PPLAPI_05795 | *pAS* | PAS domain | snp | missense_variant c.35C>T p.Ala12Val | 1 |
| Two-component system | PPLAPI_06015 | *tar* | Methyl-accepting chemotaxis protein (MCP) | snp | missense_variant c.967G>A p.Val323Met | 5 |
| Two-component system | PPLAPI_09540 | *citB* | DNA-binding response regulator%2C NarL/FixJ family%2C contains REC and HTH domains | snp | synonymous_variant c.264G>A p.Pro88Pro | 5 |
| Two-component system | PPLAPI_10490 | *tar* | Methyl-accepting chemotaxis protein (MCP) | snp | missense_variant c.231C>G p.His77Gln | 1 |
| Two-component system | PPLAPI_10795 | *hPtr* | HPt (histidine-containing phosphotransfer) domain | snp | missense_variant c.542A>T p.Asn181Ile | 4 |
| Two-component system | PPLAPI_11620 | *ompA* | Outer membrane protein OmpA and related peptidoglycan-associated (lipo)proteins | snp | synonymous_variant c.261G>A p.Gly87Gly | 4 |
| Two-component system | PPLAPI_01220 | *walK* | Sensor histidine kinase WalK | snp | missense_variant c.830C>A p.Pro277Gln | 48 |
| Two-component system | PPLAPI_12335 | *cheW* | Chemotaxis signal transduction protein CheW | snp | missense_variant c.961C>G p.Pro321Ala | 1 |
| Two-component system | PPLAPI_12365 | *tar* | Methyl-accepting chemotaxis protein (MCP) | del | conservative_inframe_deletion c.1309_1311delCAG p.Gln437del | 1 |
| Two-component system | PPLAPI_12365 | *tar* | Methyl-accepting chemotaxis protein (MCP) | snp | missense_variant c.505C>G p.Leu169Val | 32 |
| Two-component system | PPLAPI_12365 | *tar* | Methyl-accepting chemotaxis protein (MCP) | snp | missense_variant c.452T>C p.Val151Ala | 47 |
| Two-component system | PPLAPI_12470 | NA | Histidine kinase%2C HAMP region: chemotaxis sensory transducer | snp | missense_variant c.509A>T p.Lys170Met | 1 |
| Two-component system | PPLAPI_12475 | *tar* | Methyl-accepting chemotaxis protein (MCP) | snp | synonymous_variant c.1677G>A p.Gln559Gln | 7 |
| Two-component system | PPLAPI_12830 | *norR* | nitric oxide reductase transcriptional regulator NorR | snp | missense_variant c.848T>A p.Leu283Gln | 4 |
| Two-component system | PPLAPI_12905 | NA | Histidine kinase%2C HAMP region: chemotaxis sensory transducer | snp | synonymous_variant c.1557G>A p.Val519Val | 11 |
| Two-component system | PPLAPI_13005 | *ctpH* | Methyl-accepting chemotaxis protein CtpH | snp | missense_variant c.43G>A p.Gly15Ser | 4 |
| Two-component system | PPLAPI_13005 | *ctpH* | Methyl-accepting chemotaxis protein CtpH | snp | synonymous_variant c.996G>T p.Ala332Ala | 1 |
| Two-component system | PPLAPI_13170 | *pilR* | Type 4 fimbriae expression regulatory protein pilR | snp | splice_region_variant&stop_retained_variant c.957G>A p.Ter319Ter | 1 |
| Two-component system | PPLAPI_13785 | *arnB* | UDP-4-amino-4-deoxy-L-arabinose aminotransferase | del | frameshift_variant c.444_511delCGGGATTACGGTGATAGAAGACGCTGCACATGCGACCGGAACCGCTTACAAGGGTCGTCCGATCGGCA p.His148fs | 1 |
| Two-component system | PPLAPI_14120 | NA | 3-ketoacyl-CoA thiolase | snp | synonymous_variant c.192A>T p.Ser64Ser | 1 |
| Two-component system | PPLAPI_14365 | *adeC* | outer membrane channel protein that acts as part of the AdeABC multidrug efflux pump | snp | missense_variant c.1045G>A p.Val349Met | 4 |
| Two-component system | PPLAPI_14370 | *adeB* | Multidrug transporter component of the AdeABC efflux system | del | conservative_inframe_deletion c.1654_1659delCTGGTG p.Leu552_Val553del | 1 |
| Two-component system | PPLAPI_14375 | *adeA* | Multidrug efflux pump subunit AcrA (membrane-fusion protein) | snp | missense_variant c.520G>A p.Ala174Thr | 1 |
| Two-component system | PPLAPI_14580 | NA | HTH lacI-type domain-containing protein | del | frameshift_variant c.438_520delTCGCCAGGCTGGCGTCCCGGTGATAGCGCTGCTTTCAGACATACATGAGAAGGCCGATGAGCCTTACGTCGGTCAGGACAATT p.Arg147fs | 13 |
| Two-component system | PPLAPI_14585 | NA | Sigma-54 factor interaction domain-containing protein | snp | synonymous_variant c.1555C>T p.Leu519Leu | 4 |
| Two-component system | PPLAPI_14625 | *uvrY* | UvrY/SirA/GacA family response regulator transcription factor | del | frameshift_variant c.272_284delAAATGGTCCAGGC p.Glu91fs | 2 |
| Two-component system | PPLAPI_14625 | *uvrY* | UvrY/SirA/GacA family response regulator transcription factor | snp | missense_variant c.206C>A p.Pro69Gln | 1 |
| Two-component system | PPLAPI_14625 | *uvrY* | UvrY/SirA/GacA family response regulator transcription factor | del | disruptive_inframe_deletion c.47_61delCCGGCGAGGAATCAC p.Ser16_Ser20del | 2 |
| Two-component system | PPLAPI_14625 | *uvrY* | UvrY/SirA/GacA family response regulator transcription factor | snp | missense_variant c.11A>G p.Asp4Gly | 1 |
| Two-component system | PPLAPI_15165 | *paaJ* | Acetyl-CoA acetyltransferase | snp | stop_gained c.1195G>T p.Glu399* | 1 |
| Two-component system | PPLAPI_01585 | *tar* | Methyl-accepting chemotaxis protein (MCP) | snp | synonymous_variant c.933C>T p.Ala311Ala | 1 |
| Two-component system | PPLAPI_15340 | *tolC* | Outer membrane protein TolC | snp | missense_variant c.92C>A p.Pro31Gln | 1 |
| Two-component system | PPLAPI_15380 | *nodT* | RND efflux system%2C outer membrane lipoprotein%2C NodT | snp | missense_variant c.766C>T p.His256Tyr | 6 |
| Two-component system | PPLAPI_15595 | *pstS* | phosphate ABC transporter substrate-binding protein PstS | snp | missense_variant c.998T>G p.Leu333Trp | 119 |
| Two-component system | PPLAPI_16155 | NA | methyl-accepting chemotaxis protein | snp | synonymous_variant c.258G>A p.Leu86Leu | 29 |
| Two-component system | PPLAPI_16170 | NA | Methyl-accepting chemotaxis protein | snp | missense_variant c.218C>T p.Ala73Val | 5 |
| Two-component system | PPLAPI_16450 | NA | Methyl-accepting chemotaxis protein | snp | missense_variant c.701G>T p.Arg234Leu | 11 |
| Two-component system | PPLAPI_17035 | *ompR* | DNA-binding response regulator%2C OmpR family%2C contains REC and winged-helix (wHTH) domain | snp | missense_variant c.547G>T p.Asp183Tyr | 1 |
| Two-component system | PPLAPI_17115 | NA | Aerotaxis receptor Aer | snp | missense_variant c.644G>C p.Gly215Ala | 1 |
| Two-component system | PPLAPI_17330 | *ccoN* | cytochrome-c oxidase%2C cbb3-type subunit I | snp | missense_variant c.647C>G p.Thr216Ser | 5 |
| Two-component system | PPLAPI_17395 | *crp* | cAMP-binding domain of CRP or a regulatory subunit of cAMP-dependent protein kinases | snp | missense_variant c.53G>T p.Cys18Phe | 5 |
| Two-component system | PPLAPI_17395 | *crp* | cAMP-binding domain of CRP or a regulatory subunit of cAMP-dependent protein kinases | snp | synonymous_variant c.381G>A p.Leu127Leu | 1 |
| Two-component system | PPLAPI_17395 | *crp* | cAMP-binding domain of CRP or a regulatory subunit of cAMP-dependent protein kinases | snp | missense_variant c.482C>T p.Ala161Val | 4 |
| Two-component system | PPLAPI_17395 | *crp* | cAMP-binding domain of CRP or a regulatory subunit of cAMP-dependent protein kinases | snp | synonymous_variant c.498C>A p.Ala166Ala | 13 |
| Two-component system | PPLAPI_17440 | *cheA* | Chemotaxis protein CheA | del | disruptive_inframe_deletion c.831_836delACCTGC p.Pro278_Ala279del | 19 |
| Two-component system | PPLAPI_17440 | *cheA* | Chemotaxis protein CheA | ins | disruptive_inframe_insertion c.831_836dupACCTGC p.Ala279_Ala280insProAla | 5 |
| Two-component system | PPLAPI_17450 | *cheY* | chemotaxis protein CheY | snp | missense_variant c.361C>T p.Arg121Cys | 1 |
| Two-component system | PPLAPI_18435 | *mprF* | bifunctional lysylphosphatidylglycerol flippase/synthetase MprF | snp | missense_variant c.1984C>A p.Leu662Met | 1 |
| Two-component system | PPLAPI_18930 | *cheB3* | Protein-glutamate methylesterase/protein-glutamine glutaminase 3 | snp | missense_variant c.412C>G p.His138Asp | 2 |
| Two-component system | PPLAPI_18935 | *cheY* | CheY-like REC (receiver) domain%2C includes chemotaxis protein CheY and sporulation regulator Spo0F | del | frameshift_variant c.1922delT p.Met641fs | 1 |
| Two-component system | PPLAPI_19170 | NA | Anaerobic nitric oxide reductase transcription regulator | del | frameshift_variant c.542_552delAACGGAAACGG p.Glu181fs | 1 |
| Two-component system | PPLAPI_19870 | *cheY* | CheY-like REC (receiver) domain%2C includes chemotaxis protein CheY and sporulation regulator Spo0F | snp | synonymous_variant c.1755C>T p.Cys585Cys | 2 |
| Two-component system | PPLAPI_20345 | *gacS* | Sensor protein GacS | del | frameshift_variant c.693_705delCATCAATCGCATG p.Ile232fs | 2 |
| Two-component system | PPLAPI_20345 | *gacS* | Sensor protein GacS | snp | stop_gained c.1724T>A p.Leu575* | 1 |
| Two-component system | PPLAPI_20345 | *gacS* | Sensor protein GacS | del | frameshift_variant c.2098delT p.Tyr700fs | 1 |
| Two-component system | PPLAPI_20400 | *dctA-1* | C4-dicarboxylate transport protein | snp | missense_variant c.424A>G p.Thr142Ala | 13 |
| Two-component system | PPLAPI_02165 | *pilG* | twitching motility response regulator PilG | snp | missense_variant c.67G>T p.Ala23Ser | 1 |
| Two-component system | PPLAPI_02185 | *cheY* | CheY-like REC (receiver) domain%2C includes chemotaxis protein CheY and sporulation regulator Spo0F | snp | missense_variant c.1177G>T p.Asp393Tyr | 3 |
| Two-component system | PPLAPI_02185 | *cheY* | CheY-like REC (receiver) domain%2C includes chemotaxis protein CheY and sporulation regulator Spo0F | snp | missense_variant c.5914A>G p.Thr1972Ala | 1 |
| Two-component system | PPLAPI_21405 | *hisM* | ABC-type amino acid transport system%2C permease component | complex | missense_variant c.92_93delGGinsTT p.Trp31Phe | 4 |
| Two-component system | PPLAPI_21415 | NA | Aspartate/glutamate ABC-type transport system | snp | missense_variant c.556G>A p.Gly186Ser | 1 |
| Two-component system | PPLAPI_21425 | *atoC* | DNA-binding transcriptional response regulator%2C NtrC family%2C contains REC%2C AAA-type ATPase%2C and a Fis-type DNA-binding domains | ins | disruptive_inframe_insertion c.80_85dupCGCTGG p.Ala27_Leu28dup | 1 |
| Two-component system | PPLAPI_21425 | *atoC* | DNA-binding transcriptional response regulator%2C NtrC family%2C contains REC%2C AAA-type ATPase%2C and a Fis-type DNA-binding domains | snp | missense_variant c.470G>A p.Ser157Asn | 46 |
| Two-component system | PPLAPI_21690 | *nadB* | L-aspartate oxidase | snp | missense_variant c.727T>C p.Phe243Leu | 13 |
| Two-component system | PPLAPI_21690 | *nadB* | L-aspartate oxidase | snp | missense_variant c.862G>A p.Ala288Thr | 1 |
| Two-component system | PPLAPI_21710 | *hDOD* | HD-like signal output (HDOD) domain%2C no enzymatic activity | del | disruptive_inframe_deletion c.137_148delCCCTGAGCAAGG p.Ala46_Lys49del | 3 |
| Two-component system | PPLAPI_21715 | *baeS* | Signal transduction histidine kinase | snp | missense_variant c.1378C>G p.Pro460Ala | 1 |
| Two-component system | PPLAPI_21715 | *baeS* | Signal transduction histidine kinase | ins | conservative_inframe_insertion c.1004_1036dupCGTTGGCTCATTCACGCGGCGTCGCGCTGGCGC p.Pro335_Ala345dup | 4 |
| Two-component system | PPLAPI_21885 | NA | Signal transduction histidine kinase regulating C4-dicarboxylate transport system | ins | conservative_inframe_insertion c.784_786dupCGT p.Arg262dup | 1 |
| Two-component system | PPLAPI_21885 | NA | Signal transduction histidine kinase regulating C4-dicarboxylate transport system | snp | missense_variant c.1021G>C p.Val341Leu | 1 |
| Two-component system | PPLAPI_22915 | *pleD* | Two-component response regulator%2C PleD family%2C consists of two REC domains and a diguanylate cyclase (GGDEF) domain | ins | disruptive_inframe_insertion c.593_594insGCTGTTGTTTTTCAG p.Pro198_Leu199insLeuLeuPhePheSer | 1 |
| Two-component system | PPLAPI_24200 | NA | Two-component sensor | del | disruptive_inframe_deletion c.926_937delCGCAGGAGCAGG p.Ala309_Gln312del | 1 |
| Two-component system | PPLAPI_24620 | NA | Methyl-accepting chemotaxis protein | snp | missense_variant c.354C>A p.Asn118Lys | 4 |
| Two-component system | PPLAPI_25490 | NA | Sensor protein | snp | missense_variant c.994T>G p.Ser332Ala | 4 |
| Two-component system | PPLAPI_26275 | NA | Signal transduction histidine kinase regulating C4-dicarboxylate transport system | snp | missense_variant c.122G>A p.Ser41Asn | 1 |
| Two-component system | PPLAPI_26275 | NA | Signal transduction histidine kinase regulating C4-dicarboxylate transport system | snp | missense_variant c.1385G>A p.Arg462His | 9 |
| Two-component system | PPLAPI_26280 | *atoC* | DNA-binding transcriptional response regulator%2C NtrC family%2C contains REC%2C AAA-type ATPase%2C and a Fis-type DNA-binding domains | ins | conservative_inframe_insertion c.1159_1164dupCTGGAG p.Leu387_Glu388dup | 1 |
| Two-component system | PPLAPI_27230 | *ugd* | UDP-glucose 6-dehydrogenase | del | disruptive_inframe_deletion c.752_769delCCGACCCTCGCATTGGCT p.Ser251_Gly256del | 9 |
| Two-component system | PPLAPI_02645 | *tar* | Methyl-accepting chemotaxis protein (MCP) | snp | stop_gained c.14T>G p.Leu5* | 1 |
| Two-component system | PPLAPI_02645 | *tar* | Methyl-accepting chemotaxis protein (MCP) | snp | synonymous_variant c.1239T>C p.Asp413Asp | 1 |
| Two-component system | PPLAPI_27500 | *dnaA* | chromosomal replication initiator protein DnaA | snp | synonymous_variant c.999T>G p.Gly333Gly | 5 |
| Two-component system | PPLAPI_27890 | *tar* | Methyl-accepting chemotaxis protein (MCP) | snp | missense_variant c.1915A>G p.Thr639Ala | 1 |
| Two-component system | PPLAPI_27890 | *tar* | Methyl-accepting chemotaxis protein (MCP) | ins | conservative_inframe_insertion c.1871_1876dupTGGCCA p.Met624_Ala625dup | 1 |
| Two-component system | PPLAPI_03000 | *paaJ* | acetyl-CoA C-acetyltransferase | snp | missense_variant c.616G>A p.Ala206Thr | 2 |
| Two-component system | PPLAPI_03860 | NA | histidine kinase | snp | missense_variant c.727G>T p.Gly243Cys | 4 |

Table S7 Chromosomal mutations exclusive to Wytham Woods non-swarming strains. SNPs, indels and complex mutations present only in non-swarming strains. The table details each mutation, including its position on the reference genome (W163), affected locus, mutation type, and predicted functional consequence. For hypothetical proteins, AlphaFold structural predictions were used to infer potential functions. Only high-confidence predictions (pLDDT > 80) are included and are marked with (*).

| **Gene – function *alpha-fold prediction** | **Position** | **Type** | **Contig** | **Locus** | **Effect** | **Strains** |
| --- | --- | --- | --- | --- | --- | --- |
| *gntR* family transcriptional regulator * | 3049303 | snp | 1 | 2757 | Stop gained | W593a5b1, W603a5b1 |
|  |  |  |  |  | c.271C>T p.Gln91* |  |
| *gacA* - Response regulator GacA | 3202008 | del | 1 | 2898 | Frameshift variant c.272_284delAAATGGTCCAGGC p.Glu91fs | W2313a1b3, W2413a1b3 |
| *gacA* - Response regulator GacA | 3202231 | del | 1 | 2898 | Disruptive inframe deletion c.47_61delCCGGCGAGGAATCAC p.Ser16_Ser20del | W593a5b1, W603a5b1 |
| *gacS* - Signal transduction histidine-protein kinase gacS | 4422434 | del | 1 | 4037 | Frameshift variant c.693_705delCATCAATCGCATG p.Ile232fs | W573a5b1, W583a5b1 |
| *gacS* - Signal transduction histidine-protein kinase gacS | 4423840 | del | 1 | 4037 | Frameshift variant | W1513a1b2 |
|  |  |  |  |  | c.2098delT p.Tyr700fs |  |
| Hypothetical protein | 1633519 | snp | 1 | 1492 | Missense variant | W1513a1b2 |
|  |  |  |  |  | c.722C>T p.Pro241Leu |  |
| *hscB* - Co-chaperone protein HscB | 1710363 | snp | 1 | 1568 | Missense variant | W573a5b1, W583a5b1 |
|  |  |  |  |  | c.8C>T p.Thr3Ile |  |
| NA | 18531 | mnp | 4 | NA | NA | W1313a1b2, W1413a1b2, W593a5b1, W603a5b1 |
| NA | 18570 | complex | 4 | NA | NA | W1313a1b2, W1413a1b2, W593a5b1, W603a5b1 |
| Hypothetical protein | 3807753 | del | 1 | 3458 | Disruptive inframe deletion c.831_836dupACCTGC p.Ala279_Ala280insProAla | L1227a1b4, L1277a1b6, L1327a1b5, L1738a1b4, W1143a7b3 |
| NA | 4741558 | del | 1 | NA | NA | ICMP9132, ICMP7711, W1513a1b2 |
| NA | 5644449 | snp | 1 | NA | NA | W593a5b1 |

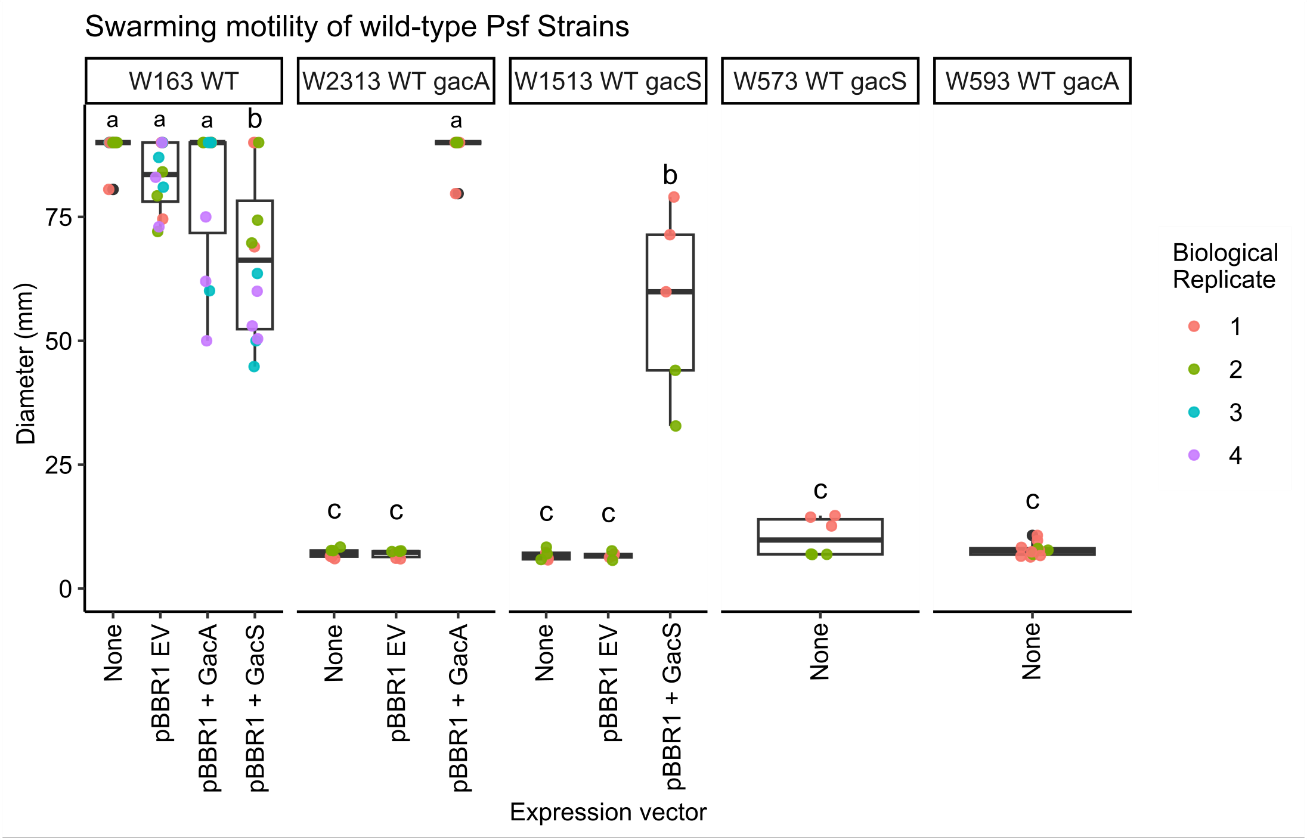
**Figure S11: Swarming motility of complemented wildtype *Psf* strains.** Diameter of bacterial growth on swarming agar after 72 hours incubation. Plots is facetted by strain: W163 WT swarming strain, W2313 WT non-swarming strain with a 1bp deletion mutation in *gacA*, W1513 WT non-swarming strain with a 13bp deletion in *gacS*, W573 non-swarming strain with a 13bp deletion in *gacS*, W593 non-swarming strain with a 15bp deletion in *gacA*. The presence of an expression vector (none, empty vector, *gacA* expression, or *gacS* expression), is indicated on the x-axis. Data points represent independent measurements, coloured by biological replicate. For statistical analysis, the mean halo diameter of three technical replicates was calculated for each biological replicate, and one-way ANOVA was performed on these biological replicate means. At least two independent biological replicates were analysed per strain. Letters above the boxplots indicate statistically significant differences in halo diameter, determined by Tukey’s HSD post hoc test (p < 0.05).

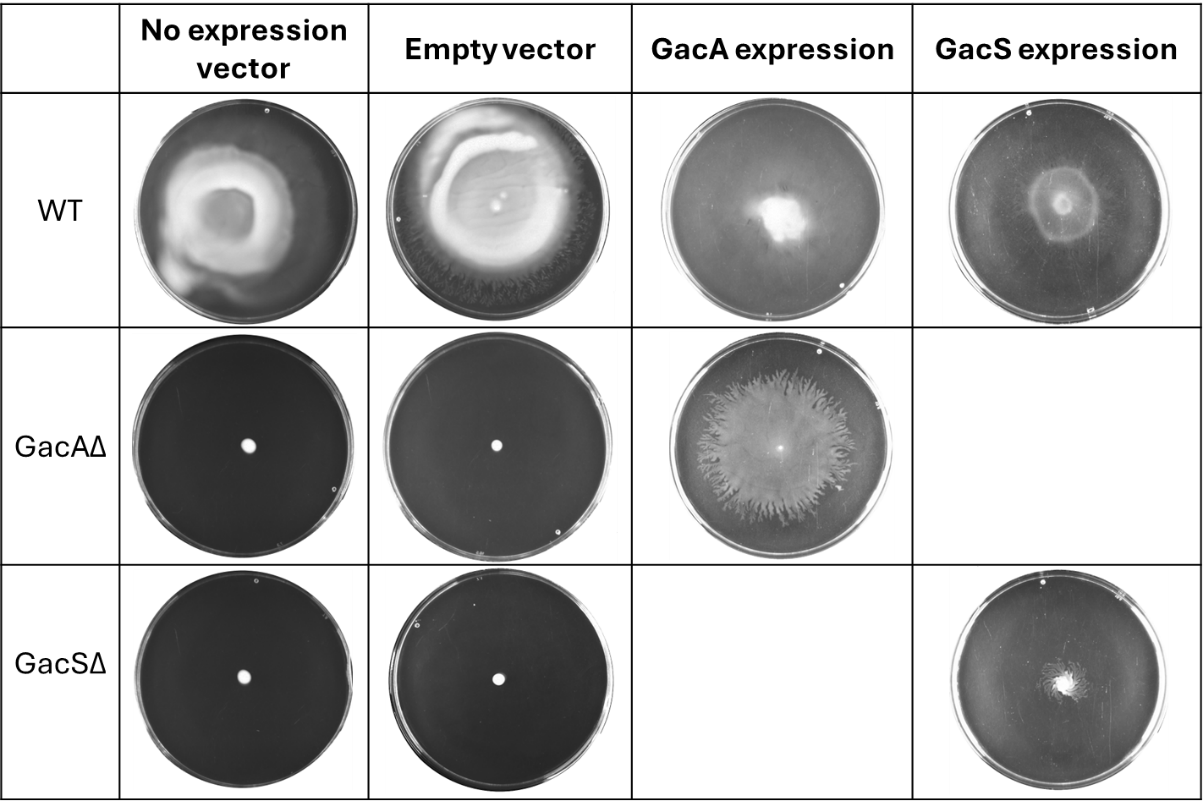

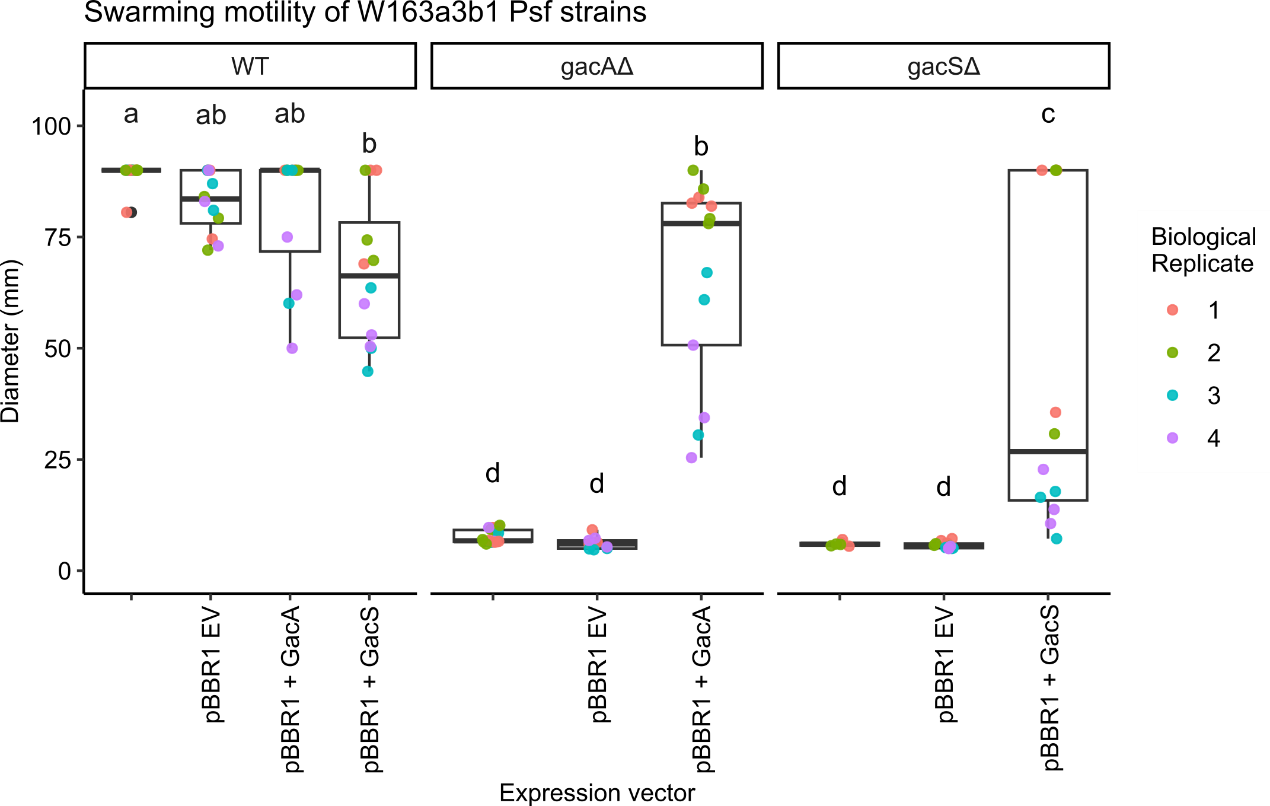
**Figure S12: Swarming motility of W163 wild-type and allele-swap introduced *gacA*/*S* mutant strains.** Bacterial growth was measured on 0.5% swarming agar after 72 hours incubation. W163 strains are grouped according to mutant background (wild-type, *gacA* mutant, or *gacS* mutant) and the presence of an expression vector (none, empty vector, *gacA* complementation, or *gacS* complementation), as indicated on the x-axis. Data points represent independent measurements, coloured by biological replicate number. For statistical analysis, the mean halo diameter of three technical replicates was calculated for each biological replicate, and one-way ANOVA was performed on these biological replicate means. At least two independent biological replicates were analysed per strain. Letters above the boxplots indicate statistically significant differences in halo diameter, determined by Tukey’s HSD post hoc test (p < 0.05). (Bottom) **:** Example of swarming motility phenotypes exhibited by different *Psf* mutant strains. Strains are categorised by their mutant background (wild-type, *gacA* mutant, *gacS* mutant) indicated in the first column, and the presence of an expression vector (none, empty vector, *gacA* expression, or *gacS* expression), as indicated in the top row.

**Table S8** **Pairwise comparisons of functional versus mutants across phenotypes.** A linear mixed-effects model was used to assess the effects of phenotype, GacA/S group (functional or mutant), and their interaction on growth (absorbance OD_600_) after 72 hours static growth, with experiment included as a random effect. Differences were estimated using Tukey-adjusted post hoc comparisons from the linear mixed-effects model. Estimates represent differences in mean response (functional − mutant) for each phenotype, with associated standard errors (SE), degrees of freedom (D.F.), t-ratios, and adjusted *P*-values.

| Contrast | Phenotype | Estimate | SE | D.F. | t.ratio | P-value |
| --- | --- | --- | --- | --- | --- | --- |
| Functional - Mutant | KBB | -0.01696 | 0.031679 | 702.0068 | -0.53552 | 0.592462 |
| Functional - Mutant | M9 | 4.80E-04 | 0.027435 | 702.0068 | 0.017503 | 0.98604 |
| Functional - Mutant | M9_EToH100 | -0.08094 | 0.031679 | 702.0068 | -2.55503 | 0.010828 |
| Functional - Mutant | M9_EToH50 | 0.00791 | 0.031679 | 702.0068 | 0.249681 | 0.802907 |
| Functional - Mutant | M9_Frax | -0.00202 | 0.05487 | 702.0068 | -0.03675 | 0.970692 |
| Functional - Mutant | M9_Frax_Gluc | 0.001136 | 0.024539 | 702.0068 | 0.046276 | 0.963103 |
| Functional - Mutant | M9_Gluc | -0.1377 | 0.027435 | 702.0068 | -5.01919 | 6.58E-07 |
| Functional - Mutant | M9_Glut | 0.006456 | 0.031679 | 702.0068 | 0.203778 | 0.838586 |
| Functional - Mutant | M9_Glut_Gluc | 0.134133 | 0.032978 | 702.1121 | 4.067332 | 5.29E-05 |
| Functional - Mutant | M9_Glyc | -0.06382 | 0.027435 | 702.0068 | -2.32607 | 0.020299 |
| Functional - Mutant | M9_Pcoum | -0.00276 | 0.027435 | 702.0068 | -0.10076 | 0.919773 |
| Functional - Mutant | M9_Pcoum_Gluc | -0.06163 | 0.027435 | 702.0068 | -2.2464 | 0.024989 |
| Functional - Mutant | M9_Quin | -0.01965 | 0.027435 | 702.0068 | -0.71632 | 0.47403 |
| Functional - Mutant | M9_Quin_Gluc | 0.043382 | 0.027435 | 702.0068 | 1.581269 | 0.114267 |
| Functional - Mutant | M9_Ser | -0.00288 | 0.027435 | 702.0068 | -0.10487 | 0.91651 |
| Functional - Mutant | M9_Ser_Gluc | -0.25651 | 0.027435 | 702.0068 | -9.34966 | 1.15E-19 |
| Functional - Mutant | M9_Suc | -0.02521 | 0.031679 | 702.0068 | -0.79569 | 0.426481 |
| Functional - Mutant | M9_Val | -0.00341 | 0.031679 | 702.0068 | -0.10766 | 0.914295 |
| Functional - Mutant | M9_Val_Gluc | -0.25911 | 0.031679 | 702.0068 | -8.17921 | 1.34E-15 |
| Functional - Mutant | TSB | 0.007051 | 0.031679 | 702.0068 | 0.222586 | 0.823922 |
| Functional - Mutant | TSB_NaCl | 0.003213 | 0.031679 | 702.0068 | 0.101436 | 0.919233 |
| Functional - Mutant | TSB_PEG | 0.115225 | 0.031679 | 702.0068 | 3.637213 | 2.96E-04 |
| Functional - Mutant | TSB_Pen | 0.010761 | 0.031679 | 702.0068 | 0.339674 | 0.734204 |
| Functional - Mutant | TSB_Strep | -0.00221 | 0.031679 | 702.0068 | -0.06975 | 0.94441 |
| Functional - Mutant | TSB_Tet | -0.00346 | 0.031679 | 702.0068 | -0.10912 | 0.913136 |

**Table S9 Pairwise comparisons of M9-media phenotypes within GacA/S groups (functional and mutant).** A linear mixed-effects model was used to assess the effects of phenotype, GacA/S group (functional or mutant), and their interaction on growth (absorbance OD_600_) after 72 hours static growth, with experiment included as a random effect. Differences were estimated using Tukey-adjusted post hoc comparisons from a linear mixed-effects model. Estimates represent differences in mean response between phenotypes, separated by functional grouping, with associated standard errors (SE), degrees of freedom (D.F.), t-ratios, and adjusted *P*-values.

| **Contrast** | **Genotype** | **Estimate** | **SE** | **D.F.** | **t.ratio** | **P-value** |
| --- | --- | --- | --- | --- | --- | --- |
| M9 - M9_EToH100 | Functional | -0.15543 | 0.034267 | 702.9885 | -4.53588 | 0.001767 |
| M9 - M9_EToH50 | Functional | -0.0979 | 0.034267 | 702.9885 | -2.85683 | 0.414304 |
| M9 - M9_Frax | Functional | 0.005769 | 0.050393 | 704.8194 | 0.114473 | 1 |
| M9 - M9_Frax_Gluc | Functional | -0.00142 | 0.030122 | 703.4071 | -0.0473 | 1 |
| M9 - M9_Gluc | Functional | -0.25699 | 0.031679 | 702.0068 | -8.11223 | 0 |
| M9 - M9_Glut | Functional | -0.33208 | 0.034267 | 702.9885 | -9.69089 | 0 |
| M9 - M9_Glut_Gluc | Functional | -0.58115 | 0.035474 | 703.0691 | -16.3825 | 0 |
| M9 - M9_Glyc | Functional | -0.28368 | 0.031679 | 702.0068 | -8.95464 | 0 |
| M9 - M9_Pcoum | Functional | -4.19E-04 | 0.031679 | 702.0068 | -0.01324 | 1 |
| M9 - M9_Pcoum_Gluc | Functional | -0.234 | 0.031679 | 702.0068 | -7.38649 | 0 |
| M9 - M9_Quin | Functional | -0.22117 | 0.031679 | 702.0068 | -6.98155 | 0 |
| M9 - M9_Quin_Gluc | Functional | -0.33172 | 0.031679 | 702.0068 | -10.4712 | 0 |
| M9 - M9_Ser | Functional | -0.00608 | 0.031679 | 702.0068 | -0.19187 | 1 |
| M9 - M9_Ser_Gluc | Functional | -0.21069 | 0.031679 | 702.0068 | -6.65058 | 1.48E-08 |
| M9 - M9_Suc | Functional | -0.222 | 0.034267 | 702.9885 | -6.47862 | 4.91E-08 |
| M9 - M9_Val | Functional | -0.09866 | 0.034272 | 703.1441 | -2.87869 | 0.397926 |
| M9 - M9_Val_Gluc | Functional | -0.10897 | 0.034272 | 703.1441 | -3.17957 | 0.207365 |
| M9_EToH100 - M9_EToH50 | Functional | 0.057536 | 0.03658 | 702.0068 | 1.572878 | 0.997826 |
| M9_EToH100 - M9_Frax | Functional | 0.1612 | 0.052253 | 704.8099 | 3.084969 | 0.259396 |
| M9_EToH100 - M9_Frax_Gluc | Functional | 0.154006 | 0.032808 | 703.6049 | 4.6942 | 8.67E-04 |
| M9_EToH100 - M9_Gluc | Functional | -0.10156 | 0.034267 | 702.9885 | -2.96375 | 0.337 |
| M9_EToH100 - M9_Glut | Functional | -0.17665 | 0.03658 | 702.0068 | -4.82904 | 4.63E-04 |
| M9_EToH100 - M9_Glut_Gluc | Functional | -0.42572 | 0.037711 | 702.0875 | -11.289 | 0 |
| M9_EToH100 - M9_Glyc | Functional | -0.12825 | 0.034267 | 702.9885 | -3.74254 | 0.03962 |
| M9_EToH100 - M9_Pcoum | Functional | 0.155012 | 0.034267 | 702.9885 | 4.523637 | 0.001865 |
| M9_EToH100 - M9_Pcoum_Gluc | Functional | -0.07857 | 0.034267 | 702.9885 | -2.29282 | 0.83448 |
| M9_EToH100 - M9_Quin | Functional | -0.06574 | 0.034267 | 702.9885 | -1.91845 | 0.970503 |
| M9_EToH100 - M9_Quin_Gluc | Functional | -0.17629 | 0.034267 | 702.9885 | -5.14456 | 9.91E-05 |
| M9_EToH100 - M9_Ser | Functional | 0.149353 | 0.034267 | 702.9885 | 4.358492 | 0.003793 |
| M9_EToH100 - M9_Ser_Gluc | Functional | -0.05525 | 0.034267 | 702.9885 | -1.61248 | 0.996878 |
| M9_EToH100 - M9_Suc | Functional | -0.06657 | 0.03658 | 702.0068 | -1.8199 | 0.98423 |
| M9_EToH100 - M9_Val | Functional | 0.056772 | 0.036717 | 704.2535 | 1.54618 | 0.998316 |
| M9_EToH100 - M9_Val_Gluc | Functional | 0.04646 | 0.036717 | 704.2535 | 1.265332 | 0.999937 |
| M9_EToH50 - M9_Frax | Functional | 0.103664 | 0.052253 | 704.8099 | 1.983869 | 0.957236 |
| M9_EToH50 - M9_Frax_Gluc | Functional | 0.09647 | 0.032808 | 703.6049 | 2.940469 | 0.353193 |
| M9_EToH50 - M9_Gluc | Functional | -0.1591 | 0.034267 | 702.9885 | -4.6428 | 0.001096 |
| M9_EToH50 - M9_Glut | Functional | -0.23418 | 0.03658 | 702.0068 | -6.40192 | 8.10E-08 |
| M9_EToH50 - M9_Glut_Gluc | Functional | -0.48325 | 0.037711 | 702.0875 | -12.8147 | 0 |
| M9_EToH50 - M9_Glyc | Functional | -0.18578 | 0.034267 | 702.9885 | -5.42159 | 2.36E-05 |
| M9_EToH50 - M9_Pcoum | Functional | 0.097476 | 0.034267 | 702.9885 | 2.844587 | 0.423585 |
| M9_EToH50 - M9_Pcoum_Gluc | Functional | -0.1361 | 0.034267 | 702.9885 | -3.97187 | 0.017544 |
| M9_EToH50 - M9_Quin | Functional | -0.12328 | 0.034267 | 702.9885 | -3.5975 | 0.063752 |
| M9_EToH50 - M9_Quin_Gluc | Functional | -0.23383 | 0.034267 | 702.9885 | -6.82361 | 2.92E-09 |
| M9_EToH50 - M9_Ser | Functional | 0.091817 | 0.034267 | 702.9885 | 2.679443 | 0.55408 |
| M9_EToH50 - M9_Ser_Gluc | Functional | -0.11279 | 0.034267 | 702.9885 | -3.29153 | 0.155729 |
| M9_EToH50 - M9_Suc | Functional | -0.12411 | 0.03658 | 702.0068 | -3.39278 | 0.117919 |
| M9_EToH50 - M9_Val | Functional | -7.64E-04 | 0.036717 | 704.2535 | -0.02081 | 1 |
| M9_EToH50 - M9_Val_Gluc | Functional | -0.01108 | 0.036717 | 704.2535 | -0.30166 | 1 |
| M9_Frax - M9_Frax_Gluc | Functional | -0.00719 | 0.049478 | 704.9976 | -0.14538 | 1 |
| M9_Frax - M9_Gluc | Functional | -0.26276 | 0.050393 | 704.8194 | -5.21421 | 6.95E-05 |
| M9_Frax - M9_Glut | Functional | -0.33785 | 0.052253 | 704.8099 | -6.46556 | 5.34E-08 |
| M9_Frax - M9_Glut_Gluc | Functional | -0.58692 | 0.053057 | 704.8202 | -11.062 | 0 |
| M9_Frax - M9_Glyc | Functional | -0.28945 | 0.050393 | 704.8194 | -5.74379 | 4.05E-06 |
| M9_Frax - M9_Pcoum | Functional | -0.00619 | 0.050393 | 704.8194 | -0.1228 | 1 |
| M9_Frax - M9_Pcoum_Gluc | Functional | -0.23977 | 0.050393 | 704.8194 | -4.75798 | 6.46E-04 |
| M9_Frax - M9_Quin | Functional | -0.22694 | 0.050393 | 704.8194 | -4.50341 | 0.002037 |
| M9_Frax - M9_Quin_Gluc | Functional | -0.33749 | 0.050393 | 704.8194 | -6.69716 | 1.01E-08 |
| M9_Frax - M9_Ser | Functional | -0.01185 | 0.050393 | 704.8194 | -0.2351 | 1 |
| M9_Frax - M9_Ser_Gluc | Functional | -0.21645 | 0.050393 | 704.8194 | -4.29535 | 0.004932 |
| M9_Frax - M9_Suc | Functional | -0.22777 | 0.052253 | 704.8099 | -4.359 | 0.003784 |
| M9_Frax - M9_Val | Functional | -0.10443 | 0.051977 | 704.4332 | -2.0091 | 0.951079 |
| M9_Frax - M9_Val_Gluc | Functional | -0.11474 | 0.051977 | 704.4332 | -2.2075 | 0.879146 |
| M9_Frax_Gluc - M9_Gluc | Functional | -0.25557 | 0.030122 | 703.4071 | -8.48436 | 0 |
| M9_Frax_Gluc - M9_Glut | Functional | -0.33065 | 0.032808 | 703.6049 | -10.0785 | 0 |
| M9_Frax_Gluc - M9_Glut_Gluc | Functional | -0.57972 | 0.034079 | 703.7711 | -17.011 | 0 |
| M9_Frax_Gluc - M9_Glyc | Functional | -0.28225 | 0.030122 | 703.4071 | -9.37031 | 0 |
| M9_Frax_Gluc - M9_Pcoum | Functional | 0.001005 | 0.030122 | 703.4071 | 0.033374 | 1 |
| M9_Frax_Gluc - M9_Pcoum_Gluc | Functional | -0.23257 | 0.030122 | 703.4071 | -7.72109 | 0 |
| M9_Frax_Gluc - M9_Quin | Functional | -0.21975 | 0.030122 | 703.4071 | -7.29521 | 0 |
| M9_Frax_Gluc - M9_Quin_Gluc | Functional | -0.3303 | 0.030122 | 703.4071 | -10.9653 | 0 |
| M9_Frax_Gluc - M9_Ser | Functional | -0.00465 | 0.030122 | 703.4071 | -0.1545 | 1 |
| M9_Frax_Gluc - M9_Ser_Gluc | Functional | -0.20926 | 0.030122 | 703.4071 | -6.94713 | 0 |
| M9_Frax_Gluc - M9_Suc | Functional | -0.22058 | 0.032808 | 703.6049 | -6.72336 | 8.13E-09 |
| M9_Frax_Gluc - M9_Val | Functional | -0.09723 | 0.03291 | 704.8265 | -2.95454 | 0.343352 |
| M9_Frax_Gluc - M9_Val_Gluc | Functional | -0.10755 | 0.03291 | 704.8265 | -3.26788 | 0.165742 |
| M9_Gluc - M9_Glut | Functional | -0.07509 | 0.034267 | 702.9885 | -2.19126 | 0.886699 |
| M9_Gluc - M9_Glut_Gluc | Functional | -0.32416 | 0.035474 | 703.0691 | -9.13795 | 0 |
| M9_Gluc - M9_Glyc | Functional | -0.02669 | 0.031679 | 702.0068 | -0.8424 | 1 |
| M9_Gluc - M9_Pcoum | Functional | 0.256571 | 0.031679 | 702.0068 | 8.098994 | 0 |
| M9_Gluc - M9_Pcoum_Gluc | Functional | 0.022991 | 0.031679 | 702.0068 | 0.72574 | 1 |
| M9_Gluc - M9_Quin | Functional | 0.035819 | 0.031679 | 702.0068 | 1.130688 | 0.999992 |
| M9_Gluc - M9_Quin_Gluc | Functional | -0.07473 | 0.031679 | 702.0068 | -2.35895 | 0.794395 |
| M9_Gluc - M9_Ser | Functional | 0.250912 | 0.031679 | 702.0068 | 7.920359 | 0 |
| M9_Gluc - M9_Ser_Gluc | Functional | 0.046304 | 0.031679 | 702.0068 | 1.461652 | 0.999292 |
| M9_Gluc - M9_Suc | Functional | 0.034987 | 0.034267 | 702.9885 | 1.021005 | 0.999999 |
| M9_Gluc - M9_Val | Functional | 0.158331 | 0.034272 | 703.1441 | 4.619789 | 0.001216 |
| M9_Gluc - M9_Val_Gluc | Functional | 0.148019 | 0.034272 | 703.1441 | 4.318904 | 0.004475 |
| M9_Glut - M9_Glut_Gluc | Functional | -0.24907 | 0.037711 | 702.0875 | -6.60472 | 2.07E-08 |
| M9_Glut - M9_Glyc | Functional | 0.048401 | 0.034267 | 702.9885 | 1.412474 | 0.999591 |
| M9_Glut - M9_Pcoum | Functional | 0.331659 | 0.034267 | 702.9885 | 9.678652 | 0 |
| M9_Glut - M9_Pcoum_Gluc | Functional | 0.098079 | 0.034267 | 702.9885 | 2.862198 | 0.410256 |
| M9_Glut - M9_Quin | Functional | 0.110908 | 0.034267 | 702.9885 | 3.236566 | 0.179746 |
| M9_Glut - M9_Quin_Gluc | Functional | 3.58E-04 | 0.034267 | 702.9885 | 0.010456 | 1 |
| M9_Glut - M9_Ser | Functional | 0.326 | 0.034267 | 702.9885 | 9.513507 | 0 |
| M9_Glut - M9_Ser_Gluc | Functional | 0.121392 | 0.034267 | 702.9885 | 3.542536 | 0.075699 |
| M9_Glut - M9_Suc | Functional | 0.110075 | 0.03658 | 702.0068 | 3.009145 | 0.306581 |
| M9_Glut - M9_Val | Functional | 0.233419 | 0.036717 | 704.2535 | 6.357163 | 1.07E-07 |
| M9_Glut - M9_Val_Gluc | Functional | 0.223107 | 0.036717 | 704.2535 | 6.076315 | 5.95E-07 |
| M9_Glut_Gluc - M9_Glyc | Functional | 0.297469 | 0.035474 | 703.0691 | 8.385646 | 0 |
| M9_Glut_Gluc - M9_Pcoum | Functional | 0.580727 | 0.035474 | 703.0691 | 16.37066 | 0 |
| M9_Glut_Gluc - M9_Pcoum_Gluc | Functional | 0.347147 | 0.035474 | 703.0691 | 9.78606 | 0 |
| M9_Glut_Gluc - M9_Quin | Functional | 0.359976 | 0.035474 | 703.0691 | 10.14769 | 0 |
| M9_Glut_Gluc - M9_Quin_Gluc | Functional | 0.249426 | 0.035474 | 703.0691 | 7.031315 | 0 |
| M9_Glut_Gluc - M9_Ser | Functional | 0.575068 | 0.035474 | 703.0691 | 16.21114 | 0 |
| M9_Glut_Gluc - M9_Ser_Gluc | Functional | 0.37046 | 0.035474 | 703.0691 | 10.44326 | 0 |
| M9_Glut_Gluc - M9_Suc | Functional | 0.359143 | 0.037711 | 702.0875 | 9.523663 | 0 |
| M9_Glut_Gluc - M9_Val | Functional | 0.482487 | 0.037861 | 704.3869 | 12.74358 | 0 |
| M9_Glut_Gluc - M9_Val_Gluc | Functional | 0.472175 | 0.037861 | 704.3869 | 12.47122 | 0 |
| M9_Glyc - M9_Pcoum | Functional | 0.283258 | 0.031679 | 702.0068 | 8.941397 | 0 |
| M9_Glyc - M9_Pcoum_Gluc | Functional | 0.049678 | 0.031679 | 702.0068 | 1.568144 | 0.997921 |
| M9_Glyc - M9_Quin | Functional | 0.062506 | 0.031679 | 702.0068 | 1.973091 | 0.959678 |
| M9_Glyc - M9_Quin_Gluc | Functional | -0.04804 | 0.031679 | 702.0068 | -1.51654 | 0.998744 |
| M9_Glyc - M9_Ser | Functional | 0.277599 | 0.031679 | 702.0068 | 8.762763 | 0 |
| M9_Glyc - M9_Ser_Gluc | Functional | 0.072991 | 0.031679 | 702.0068 | 2.304055 | 0.82799 |
| M9_Glyc - M9_Suc | Functional | 0.061674 | 0.034267 | 702.9885 | 1.799794 | 0.986267 |
| M9_Glyc - M9_Val | Functional | 0.185018 | 0.034272 | 703.1441 | 5.398458 | 2.67E-05 |
| M9_Glyc - M9_Val_Gluc | Functional | 0.174706 | 0.034272 | 703.1441 | 5.097573 | 1.25E-04 |
| M9_Pcoum - M9_Pcoum_Gluc | Functional | -0.23358 | 0.031679 | 702.0068 | -7.37325 | 0 |
| M9_Pcoum - M9_Quin | Functional | -0.22075 | 0.031679 | 702.0068 | -6.96831 | 0 |
| M9_Pcoum - M9_Quin_Gluc | Functional | -0.3313 | 0.031679 | 702.0068 | -10.4579 | 0 |
| M9_Pcoum - M9_Ser | Functional | -0.00566 | 0.031679 | 702.0068 | -0.17863 | 1 |
| M9_Pcoum - M9_Ser_Gluc | Functional | -0.21027 | 0.031679 | 702.0068 | -6.63734 | 1.63E-08 |
| M9_Pcoum - M9_Suc | Functional | -0.22158 | 0.034267 | 702.9885 | -6.46638 | 5.33E-08 |
| M9_Pcoum - M9_Val | Functional | -0.09824 | 0.034272 | 703.1441 | -2.86645 | 0.407062 |
| M9_Pcoum - M9_Val_Gluc | Functional | -0.10855 | 0.034272 | 703.1441 | -3.16733 | 0.213662 |
| M9_Pcoum_Gluc - M9_Quin | Functional | 0.012828 | 0.031679 | 702.0068 | 0.404947 | 1 |
| M9_Pcoum_Gluc - M9_Quin_Gluc | Functional | -0.09772 | 0.031679 | 702.0068 | -3.08469 | 0.259577 |
| M9_Pcoum_Gluc - M9_Ser | Functional | 0.227921 | 0.031679 | 702.0068 | 7.194619 | 0 |
| M9_Pcoum_Gluc - M9_Ser_Gluc | Functional | 0.023313 | 0.031679 | 702.0068 | 0.735911 | 1 |
| M9_Pcoum_Gluc - M9_Suc | Functional | 0.011996 | 0.034267 | 702.9885 | 0.35007 | 1 |
| M9_Pcoum_Gluc - M9_Val | Functional | 0.13534 | 0.034272 | 703.1441 | 3.948957 | 0.019094 |
| M9_Pcoum_Gluc - M9_Val_Gluc | Functional | 0.125028 | 0.034272 | 703.1441 | 3.648072 | 0.054205 |
| M9_Quin - M9_Quin_Gluc | Functional | -0.11055 | 0.031679 | 702.0068 | -3.48963 | 0.088907 |
| M9_Quin - M9_Ser | Functional | 0.215092 | 0.031679 | 702.0068 | 6.789671 | 4.40E-09 |
| M9_Quin - M9_Ser_Gluc | Functional | 0.010485 | 0.031679 | 702.0068 | 0.330964 | 1 |
| M9_Quin - M9_Suc | Functional | -8.33E-04 | 0.034267 | 702.9885 | -0.0243 | 1 |
| M9_Quin - M9_Val | Functional | 0.122511 | 0.034272 | 703.1441 | 3.574647 | 0.06851 |
| M9_Quin - M9_Val_Gluc | Functional | 0.112199 | 0.034272 | 703.1441 | 3.273762 | 0.163214 |
| M9_Quin_Gluc - M9_Ser | Functional | 0.325642 | 0.031679 | 702.0068 | 10.2793 | 0 |
| M9_Quin_Gluc - M9_Ser_Gluc | Functional | 0.121034 | 0.031679 | 702.0068 | 3.820597 | 0.03028 |
| M9_Quin_Gluc - M9_Suc | Functional | 0.109717 | 0.034267 | 702.9885 | 3.201812 | 0.196256 |
| M9_Quin_Gluc - M9_Val | Functional | 0.233061 | 0.034272 | 703.1441 | 6.800262 | 3.85E-09 |
| M9_Quin_Gluc - M9_Val_Gluc | Functional | 0.222749 | 0.034272 | 703.1441 | 6.499376 | 4.28E-08 |
| M9_Ser - M9_Ser_Gluc | Functional | -0.20461 | 0.031679 | 702.0068 | -6.45871 | 5.61E-08 |
| M9_Ser - M9_Suc | Functional | -0.21592 | 0.034267 | 702.9885 | -6.30124 | 1.52E-07 |
| M9_Ser - M9_Val | Functional | -0.09258 | 0.034272 | 703.1441 | -2.70133 | 0.536456 |
| M9_Ser - M9_Val_Gluc | Functional | -0.10289 | 0.034272 | 703.1441 | -3.00222 | 0.311118 |
| M9_Ser_Gluc - M9_Suc | Functional | -0.01132 | 0.034267 | 702.9885 | -0.33027 | 1 |
| M9_Ser_Gluc - M9_Val | Functional | 0.112027 | 0.034272 | 703.1441 | 3.268724 | 0.165386 |
| M9_Ser_Gluc - M9_Val_Gluc | Functional | 0.101715 | 0.034272 | 703.1441 | 2.967838 | 0.334197 |
| M9_Suc - M9_Val | Functional | 0.123344 | 0.036717 | 704.2535 | 3.359273 | 0.129531 |
| M9_Suc - M9_Val_Gluc | Functional | 0.113032 | 0.036717 | 704.2535 | 3.078425 | 0.263282 |
| M9_Val - M9_Val_Gluc | Functional | -0.01031 | 0.03658 | 702.0068 | -0.2819 | 1 |
| M9 - M9_EToH100 | Mutant | -0.23685 | 0.024265 | 703.7745 | -9.76094 | 0 |
| M9 - M9_EToH50 | Mutant | -0.09047 | 0.024265 | 703.7745 | -3.72817 | 0.041588 |
| M9 - M9_Frax | Mutant | 0.003272 | 0.035846 | 704.2557 | 0.091272 | 1 |
| M9 - M9_Frax_Gluc | Mutant | -7.69E-04 | 0.021348 | 704.3668 | -0.03604 | 1 |
| M9 - M9_Gluc | Mutant | -0.39517 | 0.022401 | 702.0068 | -17.6411 | 0 |
| M9 - M9_Glut | Mutant | -0.3261 | 0.024265 | 703.7745 | -13.439 | 0 |
| M9 - M9_Glut_Gluc | Mutant | -0.44749 | 0.024265 | 703.7745 | -18.4416 | 0 |
| M9 - M9_Glyc | Mutant | -0.34797 | 0.022401 | 702.0068 | -15.534 | 0 |
| M9 - M9_Pcoum | Mutant | -0.00366 | 0.022401 | 702.0068 | -0.16356 | 1 |
| M9 - M9_Pcoum_Gluc | Mutant | -0.29611 | 0.022401 | 702.0068 | -13.2188 | 0 |
| M9 - M9_Quin | Mutant | -0.2413 | 0.022401 | 702.0068 | -10.7721 | 0 |
| M9 - M9_Quin_Gluc | Mutant | -0.28882 | 0.022401 | 702.0068 | -12.8933 | 0 |
| M9 - M9_Ser | Mutant | -0.00944 | 0.022401 | 702.0068 | -0.42123 | 1 |
| M9 - M9_Ser_Gluc | Mutant | -0.46768 | 0.022401 | 702.0068 | -20.8777 | 0 |
| M9 - M9_Suc | Mutant | -0.24769 | 0.024265 | 703.7745 | -10.2076 | 0 |
| M9 - M9_Val | Mutant | -0.10255 | 0.024273 | 704.0094 | -4.2249 | 0.006576 |
| M9 - M9_Val_Gluc | Mutant | -0.36856 | 0.024273 | 704.0094 | -15.1842 | 0 |
| M9_EToH100 - M9_EToH50 | Mutant | 0.146388 | 0.025866 | 702.0068 | 5.659439 | 6.49E-06 |
| M9_EToH100 - M9_Frax | Mutant | 0.240125 | 0.037314 | 698.5572 | 6.435326 | 6.57E-08 |
| M9_EToH100 - M9_Frax_Gluc | Mutant | 0.236084 | 0.023262 | 704.5972 | 10.149 | 0 |
| M9_EToH100 - M9_Gluc | Mutant | -0.15832 | 0.024265 | 703.7745 | -6.5245 | 3.61E-08 |
| M9_EToH100 - M9_Glut | Mutant | -0.08925 | 0.025866 | 702.0068 | -3.45046 | 0.09985 |
| M9_EToH100 - M9_Glut_Gluc | Mutant | -0.21064 | 0.025866 | 702.0068 | -8.14349 | 0 |
| M9_EToH100 - M9_Glyc | Mutant | -0.11112 | 0.024265 | 703.7745 | -4.57937 | 0.001457 |
| M9_EToH100 - M9_Pcoum | Mutant | 0.233189 | 0.024265 | 703.7745 | 9.609949 | 0 |
| M9_EToH100 - M9_Pcoum_Gluc | Mutant | -0.05926 | 0.024265 | 703.7745 | -2.44202 | 0.738087 |
| M9_EToH100 - M9_Quin | Mutant | -0.00445 | 0.024265 | 703.7745 | -0.1834 | 1 |
| M9_EToH100 - M9_Quin_Gluc | Mutant | -0.05196 | 0.024265 | 703.7745 | -2.14153 | 0.907954 |
| M9_EToH100 - M9_Ser | Mutant | 0.227417 | 0.024265 | 703.7745 | 9.372085 | 0 |
| M9_EToH100 - M9_Ser_Gluc | Mutant | -0.23082 | 0.024265 | 703.7745 | -9.51241 | 0 |
| M9_EToH100 - M9_Suc | Mutant | -0.01084 | 0.025866 | 702.0068 | -0.41899 | 1 |
| M9_EToH100 - M9_Val | Mutant | 0.134303 | 0.02606 | 705 | 5.153615 | 9.46E-05 |
| M9_EToH100 - M9_Val_Gluc | Mutant | -0.13171 | 0.02606 | 705 | -5.05414 | 1.56E-04 |
| M9_EToH50 - M9_Frax | Mutant | 0.093737 | 0.037314 | 698.5572 | 2.512153 | 0.686351 |
| M9_EToH50 - M9_Frax_Gluc | Mutant | 0.089696 | 0.023262 | 704.5972 | 3.855949 | 0.026725 |
| M9_EToH50 - M9_Gluc | Mutant | -0.30471 | 0.024265 | 703.7745 | -12.5573 | 0 |
| M9_EToH50 - M9_Glut | Mutant | -0.23564 | 0.025866 | 702.0068 | -9.1099 | 0 |
| M9_EToH50 - M9_Glut_Gluc | Mutant | -0.35703 | 0.025866 | 702.0068 | -13.8029 | 0 |
| M9_EToH50 - M9_Glyc | Mutant | -0.25751 | 0.024265 | 703.7745 | -10.6121 | 0 |
| M9_EToH50 - M9_Pcoum | Mutant | 0.086802 | 0.024265 | 703.7745 | 3.577181 | 0.067966 |
| M9_EToH50 - M9_Pcoum_Gluc | Mutant | -0.20564 | 0.024265 | 703.7745 | -8.47479 | 0 |
| M9_EToH50 - M9_Quin | Mutant | -0.15084 | 0.024265 | 703.7745 | -6.21617 | 2.57E-07 |
| M9_EToH50 - M9_Quin_Gluc | Mutant | -0.19835 | 0.024265 | 703.7745 | -8.17429 | 0 |
| M9_EToH50 - M9_Ser | Mutant | 0.08103 | 0.024265 | 703.7745 | 3.339316 | 0.136868 |
| M9_EToH50 - M9_Ser_Gluc | Mutant | -0.37721 | 0.024265 | 703.7745 | -15.5452 | 0 |
| M9_EToH50 - M9_Suc | Mutant | -0.15723 | 0.025866 | 702.0068 | -6.07842 | 5.88E-07 |
| M9_EToH50 - M9_Val | Mutant | -0.01208 | 0.02606 | 705 | -0.46372 | 1 |
| M9_EToH50 - M9_Val_Gluc | Mutant | -0.2781 | 0.02606 | 705 | -10.6715 | 0 |
| M9_Frax - M9_Frax_Gluc | Mutant | -0.00404 | 0.035267 | 701.7958 | -0.11459 | 1 |
| M9_Frax - M9_Gluc | Mutant | -0.39844 | 0.035846 | 704.2557 | -11.1153 | 0 |
| M9_Frax - M9_Glut | Mutant | -0.32937 | 0.037314 | 698.5572 | -8.82722 | 0 |
| M9_Frax - M9_Glut_Gluc | Mutant | -0.45077 | 0.037314 | 698.5572 | -12.0805 | 0 |
| M9_Frax - M9_Glyc | Mutant | -0.35125 | 0.035846 | 704.2557 | -9.79863 | 0 |
| M9_Frax - M9_Pcoum | Mutant | -0.00694 | 0.035846 | 704.2557 | -0.19348 | 1 |
| M9_Frax - M9_Pcoum_Gluc | Mutant | -0.29938 | 0.035846 | 704.2557 | -8.3518 | 0 |
| M9_Frax - M9_Quin | Mutant | -0.24458 | 0.035846 | 704.2557 | -6.82288 | 2.87E-09 |
| M9_Frax - M9_Quin_Gluc | Mutant | -0.29209 | 0.035846 | 704.2557 | -8.14839 | 0 |
| M9_Frax - M9_Ser | Mutant | -0.01271 | 0.035846 | 704.2557 | -0.3545 | 1 |
| M9_Frax - M9_Ser_Gluc | Mutant | -0.47095 | 0.035846 | 704.2557 | -13.1379 | 0 |
| M9_Frax - M9_Suc | Mutant | -0.25096 | 0.037314 | 698.5572 | -6.72577 | 8.30E-09 |
| M9_Frax - M9_Val | Mutant | -0.10582 | 0.036926 | 704.9536 | -2.86577 | 0.407568 |
| M9_Frax - M9_Val_Gluc | Mutant | -0.37184 | 0.036926 | 704.9536 | -10.0697 | 0 |
| M9_Frax_Gluc - M9_Gluc | Mutant | -0.3944 | 0.021348 | 704.3668 | -18.4753 | 0 |
| M9_Frax_Gluc - M9_Glut | Mutant | -0.32533 | 0.023262 | 704.5972 | -13.9858 | 0 |
| M9_Frax_Gluc - M9_Glut_Gluc | Mutant | -0.44672 | 0.023262 | 704.5972 | -19.2042 | 0 |
| M9_Frax_Gluc - M9_Glyc | Mutant | -0.3472 | 0.021348 | 704.3668 | -16.2643 | 0 |
| M9_Frax_Gluc - M9_Pcoum | Mutant | -0.00289 | 0.021348 | 704.3668 | -0.13559 | 1 |
| M9_Frax_Gluc - M9_Pcoum_Gluc | Mutant | -0.29534 | 0.021348 | 704.3668 | -13.8348 | 0 |
| M9_Frax_Gluc - M9_Quin | Mutant | -0.24053 | 0.021348 | 704.3668 | -11.2675 | 0 |
| M9_Frax_Gluc - M9_Quin_Gluc | Mutant | -0.28805 | 0.021348 | 704.3668 | -13.4932 | 0 |
| M9_Frax_Gluc - M9_Ser | Mutant | -0.00867 | 0.021348 | 704.3668 | -0.40596 | 1 |
| M9_Frax_Gluc - M9_Ser_Gluc | Mutant | -0.46691 | 0.021348 | 704.3668 | -21.8715 | 0 |
| M9_Frax_Gluc - M9_Suc | Mutant | -0.24692 | 0.023262 | 704.5972 | -10.6149 | 0 |
| M9_Frax_Gluc - M9_Val | Mutant | -0.10178 | 0.023406 | 704.2251 | -4.3485 | 0.003954 |
| M9_Frax_Gluc - M9_Val_Gluc | Mutant | -0.36779 | 0.023406 | 704.2251 | -15.7137 | 0 |
| M9_Gluc - M9_Glut | Mutant | 0.069069 | 0.024265 | 703.7745 | 2.846419 | 0.422189 |
| M9_Gluc - M9_Glut_Gluc | Mutant | -0.05232 | 0.024265 | 703.7745 | -2.15619 | 0.901985 |
| M9_Gluc - M9_Glyc | Mutant | 0.047199 | 0.022401 | 702.0068 | 2.107048 | 0.921022 |
| M9_Gluc - M9_Pcoum | Mutant | 0.391509 | 0.022401 | 702.0068 | 17.47753 | 0 |
| M9_Gluc - M9_Pcoum_Gluc | Mutant | 0.099063 | 0.022401 | 702.0068 | 4.422314 | 0.002894 |
| M9_Gluc - M9_Quin | Mutant | 0.153869 | 0.022401 | 702.0068 | 6.868947 | 1.50E-09 |
| M9_Gluc - M9_Quin_Gluc | Mutant | 0.106355 | 0.022401 | 702.0068 | 4.747825 | 6.78E-04 |
| M9_Gluc - M9_Ser | Mutant | 0.385737 | 0.022401 | 702.0068 | 17.21987 | 0 |
| M9_Gluc - M9_Ser_Gluc | Mutant | -0.0725 | 0.022401 | 702.0068 | -3.23663 | 0.179721 |
| M9_Gluc - M9_Suc | Mutant | 0.147482 | 0.024265 | 703.7745 | 6.077873 | 5.89E-07 |
| M9_Gluc - M9_Val | Mutant | 0.292622 | 0.024273 | 704.0094 | 12.05555 | 0 |
| M9_Gluc - M9_Val_Gluc | Mutant | 0.026609 | 0.024273 | 704.0094 | 1.096244 | 0.999995 |
| M9_Glut - M9_Glut_Gluc | Mutant | -0.12139 | 0.025866 | 702.0068 | -4.69303 | 8.72E-04 |
| M9_Glut - M9_Glyc | Mutant | -0.02187 | 0.024265 | 703.7745 | -0.90129 | 1 |
| M9_Glut - M9_Pcoum | Mutant | 0.322439 | 0.024265 | 703.7745 | 13.28803 | 0 |
| M9_Glut - M9_Pcoum_Gluc | Mutant | 0.029993 | 0.024265 | 703.7745 | 1.236055 | 0.999958 |
| M9_Glut - M9_Quin | Mutant | 0.0848 | 0.024265 | 703.7745 | 3.494673 | 0.087565 |
| M9_Glut - M9_Quin_Gluc | Mutant | 0.037285 | 0.024265 | 703.7745 | 1.536552 | 0.998467 |
| M9_Glut - M9_Ser | Mutant | 0.316667 | 0.024265 | 703.7745 | 13.05016 | 0 |
| M9_Glut - M9_Ser_Gluc | Mutant | -0.14157 | 0.024265 | 703.7745 | -5.83433 | 2.43E-06 |
| M9_Glut - M9_Suc | Mutant | 0.078413 | 0.025866 | 702.0068 | 3.03148 | 0.292198 |
| M9_Glut - M9_Val | Mutant | 0.223553 | 0.02606 | 705 | 8.578413 | 0 |
| M9_Glut - M9_Val_Gluc | Mutant | -0.04246 | 0.02606 | 705 | -1.62934 | 0.996378 |
| M9_Glut_Gluc - M9_Glyc | Mutant | 0.09952 | 0.024265 | 703.7745 | 4.101318 | 0.010729 |
| M9_Glut_Gluc - M9_Pcoum | Mutant | 0.443829 | 0.024265 | 703.7745 | 18.29064 | 0 |
| M9_Glut_Gluc - M9_Pcoum_Gluc | Mutant | 0.151384 | 0.024265 | 703.7745 | 6.238664 | 2.24E-07 |
| M9_Glut_Gluc - M9_Quin | Mutant | 0.20619 | 0.024265 | 703.7745 | 8.497282 | 0 |
| M9_Glut_Gluc - M9_Quin_Gluc | Mutant | 0.158675 | 0.024265 | 703.7745 | 6.539161 | 3.27E-08 |
| M9_Glut_Gluc - M9_Ser | Mutant | 0.438058 | 0.024265 | 703.7745 | 18.05277 | 0 |
| M9_Glut_Gluc - M9_Ser_Gluc | Mutant | -0.02018 | 0.024265 | 703.7745 | -0.83172 | 1 |
| M9_Glut_Gluc - M9_Suc | Mutant | 0.199803 | 0.025866 | 702.0068 | 7.72451 | 0 |
| M9_Glut_Gluc - M9_Val | Mutant | 0.344943 | 0.02606 | 705 | 13.23653 | 0 |
| M9_Glut_Gluc - M9_Val_Gluc | Mutant | 0.07893 | 0.02606 | 705 | 3.028777 | 0.293903 |
| M9_Glyc - M9_Pcoum | Mutant | 0.344309 | 0.022401 | 702.0068 | 15.37049 | 0 |
| M9_Glyc - M9_Pcoum_Gluc | Mutant | 0.051864 | 0.022401 | 702.0068 | 2.315266 | 0.821383 |
| M9_Glyc - M9_Quin | Mutant | 0.10667 | 0.022401 | 702.0068 | 4.761899 | 6.35E-04 |
| M9_Glyc - M9_Quin_Gluc | Mutant | 0.059155 | 0.022401 | 702.0068 | 2.640777 | 0.585206 |
| M9_Glyc - M9_Ser | Mutant | 0.338537 | 0.022401 | 702.0068 | 15.11282 | 0 |
| M9_Glyc - M9_Ser_Gluc | Mutant | -0.1197 | 0.022401 | 702.0068 | -5.34368 | 3.56E-05 |
| M9_Glyc - M9_Suc | Mutant | 0.100283 | 0.024265 | 703.7745 | 4.132745 | 0.009491 |
| M9_Glyc - M9_Val | Mutant | 0.245423 | 0.024273 | 704.0094 | 10.11102 | 0 |
| M9_Glyc - M9_Val_Gluc | Mutant | -0.02059 | 0.024273 | 704.0094 | -0.84829 | 1 |
| M9_Pcoum - M9_Pcoum_Gluc | Mutant | -0.29245 | 0.022401 | 702.0068 | -13.0552 | 0 |
| M9_Pcoum - M9_Quin | Mutant | -0.23764 | 0.022401 | 702.0068 | -10.6086 | 0 |
| M9_Pcoum - M9_Quin_Gluc | Mutant | -0.28515 | 0.022401 | 702.0068 | -12.7297 | 0 |
| M9_Pcoum - M9_Ser | Mutant | -0.00577 | 0.022401 | 702.0068 | -0.25767 | 1 |
| M9_Pcoum - M9_Ser_Gluc | Mutant | -0.46401 | 0.022401 | 702.0068 | -20.7142 | 0 |
| M9_Pcoum - M9_Suc | Mutant | -0.24403 | 0.024265 | 703.7745 | -10.0566 | 0 |
| M9_Pcoum - M9_Val | Mutant | -0.09889 | 0.024273 | 704.0094 | -4.07395 | 0.011926 |
| M9_Pcoum - M9_Val_Gluc | Mutant | -0.3649 | 0.024273 | 704.0094 | -15.0333 | 0 |
| M9_Pcoum_Gluc - M9_Quin | Mutant | 0.054806 | 0.022401 | 702.0068 | 2.446633 | 0.734789 |
| M9_Pcoum_Gluc - M9_Quin_Gluc | Mutant | 0.007292 | 0.022401 | 702.0068 | 0.325511 | 1 |
| M9_Pcoum_Gluc - M9_Ser | Mutant | 0.286674 | 0.022401 | 702.0068 | 12.79756 | 0 |
| M9_Pcoum_Gluc - M9_Ser_Gluc | Mutant | -0.17157 | 0.022401 | 702.0068 | -7.65895 | 0 |
| M9_Pcoum_Gluc - M9_Suc | Mutant | 0.048419 | 0.024265 | 703.7745 | 1.995399 | 0.954497 |
| M9_Pcoum_Gluc - M9_Val | Mutant | 0.19356 | 0.024273 | 704.0094 | 7.974329 | 0 |
| M9_Pcoum_Gluc - M9_Val_Gluc | Mutant | -0.07245 | 0.024273 | 704.0094 | -2.98498 | 0.322573 |
| M9_Quin - M9_Quin_Gluc | Mutant | -0.04751 | 0.022401 | 702.0068 | -2.12112 | 0.915849 |
| M9_Quin - M9_Ser | Mutant | 0.231868 | 0.022401 | 702.0068 | 10.35092 | 0 |
| M9_Quin - M9_Ser_Gluc | Mutant | -0.22637 | 0.022401 | 702.0068 | -10.1056 | 0 |
| M9_Quin - M9_Suc | Mutant | -0.00639 | 0.024265 | 703.7745 | -0.26322 | 1 |
| M9_Quin - M9_Val | Mutant | 0.138753 | 0.024273 | 704.0094 | 5.716403 | 4.72E-06 |
| M9_Quin - M9_Val_Gluc | Mutant | -0.12726 | 0.024273 | 704.0094 | -5.24291 | 6.00E-05 |
| M9_Quin_Gluc - M9_Ser | Mutant | 0.279382 | 0.022401 | 702.0068 | 12.47204 | 0 |
| M9_Quin_Gluc - M9_Ser_Gluc | Mutant | -0.17886 | 0.022401 | 702.0068 | -7.98446 | 0 |
| M9_Quin_Gluc - M9_Suc | Mutant | 0.041127 | 0.024265 | 703.7745 | 1.694902 | 0.993733 |
| M9_Quin_Gluc - M9_Val | Mutant | 0.186268 | 0.024273 | 704.0094 | 7.673925 | 0 |
| M9_Quin_Gluc - M9_Val_Gluc | Mutant | -0.07975 | 0.024273 | 704.0094 | -3.28538 | 0.158283 |
| M9_Ser - M9_Ser_Gluc | Mutant | -0.45824 | 0.022401 | 702.0068 | -20.4565 | 0 |
| M9_Ser - M9_Suc | Mutant | -0.23825 | 0.024265 | 703.7745 | -9.81871 | 0 |
| M9_Ser - M9_Val | Mutant | -0.09311 | 0.024273 | 704.0094 | -3.83616 | 0.028665 |
| M9_Ser - M9_Val_Gluc | Mutant | -0.35913 | 0.024273 | 704.0094 | -14.7955 | 0 |
| M9_Ser_Gluc - M9_Suc | Mutant | 0.219985 | 0.024265 | 703.7745 | 9.065782 | 0 |
| M9_Ser_Gluc - M9_Val | Mutant | 0.365125 | 0.024273 | 704.0094 | 15.04255 | 0 |
| M9_Ser_Gluc - M9_Val_Gluc | Mutant | 0.099112 | 0.024273 | 704.0094 | 4.083237 | 0.011507 |
| M9_Suc - M9_Val | Mutant | 0.14514 | 0.02606 | 705 | 5.569484 | 1.06E-05 |
| M9_Suc - M9_Val_Gluc | Mutant | -0.12087 | 0.02606 | 705 | -4.63827 | 0.001119 |
| M9_Val - M9_Val_Gluc | Mutant | -0.26601 | 0.025866 | 702.0068 | -10.2843 | 0 |

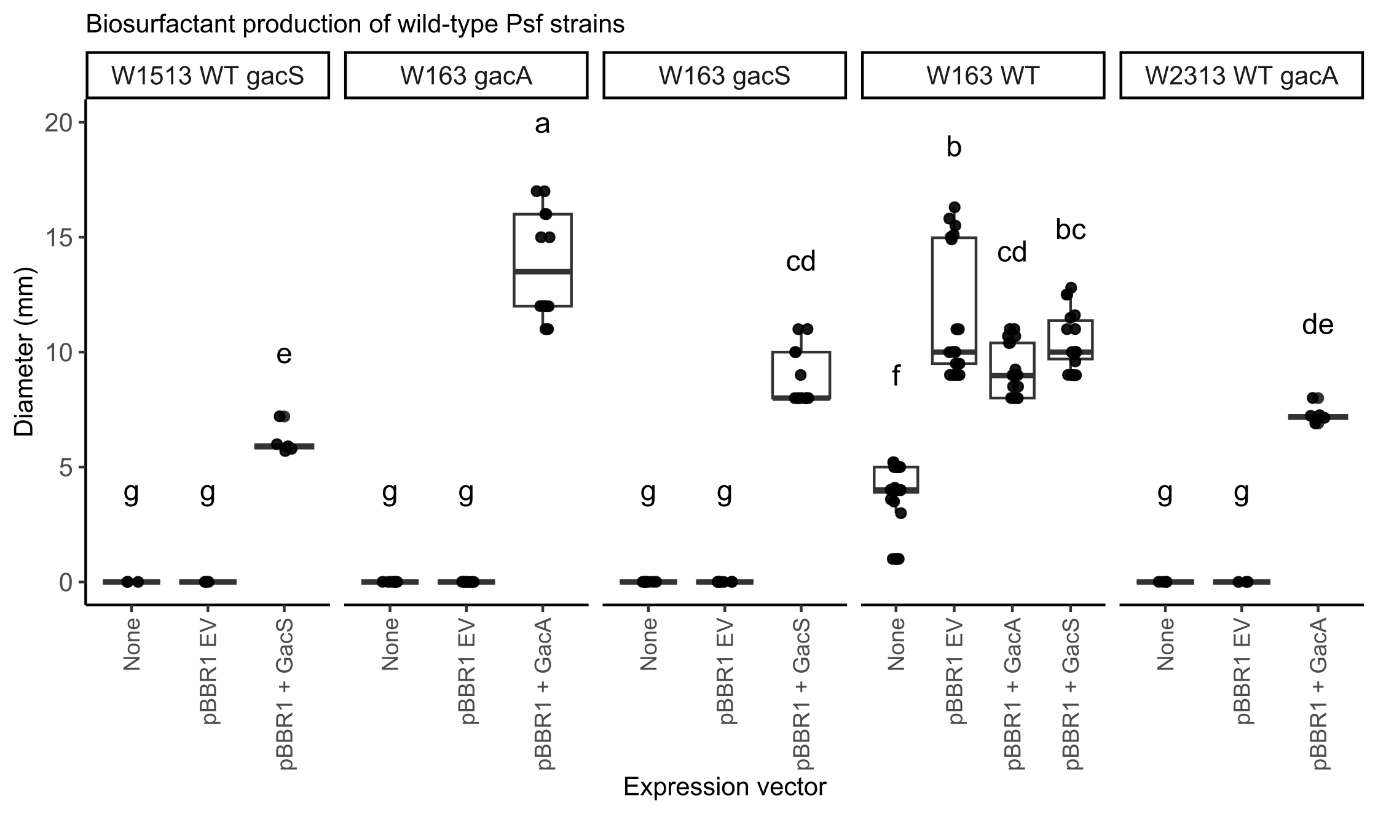

**Figure S13 Biosurfactant production of complemented wildtype and allel-swapped *Psf* strains.** Diameter of biosurfactant production on KB agar after 24 hours incubation. Plots is facetted by strain: WT strains W163, W2313 *gacA^Q91fs^*, W1513 *gacS^Y700fs^*, and mutant strains W163 *gacA^Q91fs^* and W163 *gacS^Y700fs^*. The presence of an expression vector (none, empty vector, *gacA* expression, or *gacS* expression), is indicated on the x-axis. Data points represent independent measurements. For statistical analysis, the mean halo diameter of three technical replicates was calculated for each biological replicate, and one-way ANOVA was performed on these biological replicate means. At least three independent biological replicates were analysed per strain. Letters above the boxplots indicate statistically significant differences in halo diameter, determined by Tukey’s HSD post hoc test (p < 0.05).

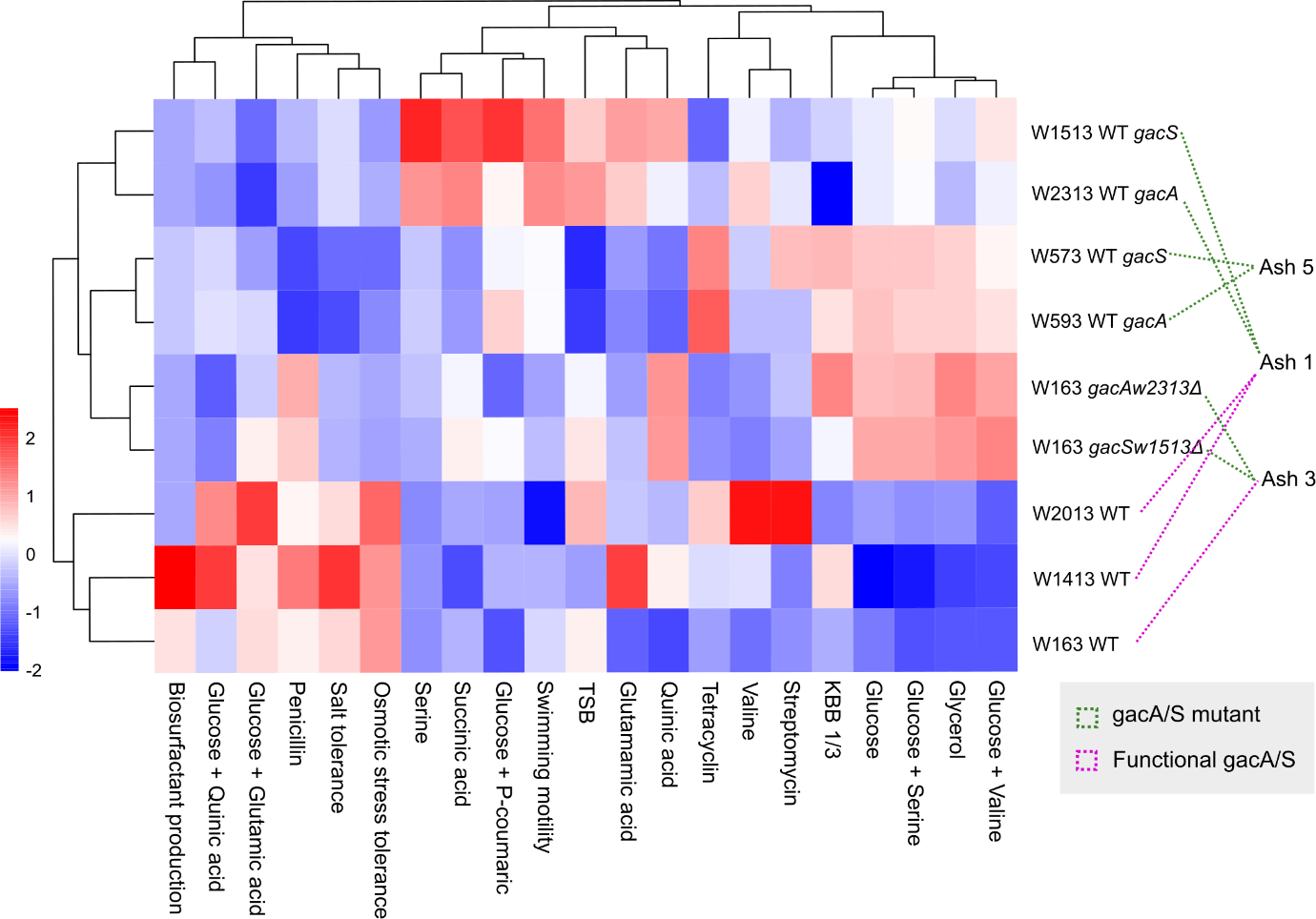

**Figure S14 Heatmap of phenotypic profiles across strains.** Mean phenotype values were calculated for each strain and phenotype, from three biological replicates. Values represent mean optical density (OD) (growth in different media) or diameter (motility and biosurfactant production). A matrix was generated combining data from multiple assays (growth, swimming motility, and biosurfactant production). The resulting strain-by-phenotype matrix was Z-score normalised by phenotype (column-wise scaling), and hierarchical clustering was applied to both strains and phenotypes. Colours indicate relative phenotype values (blue = below mean, white = mean, red = above mean). Strains lacking sufficient replicates or missing data for specific conditions were excluded prior to analysis.

Table S10 Pairwise comparison of *in planta* population sizes (CFU/ml) after 3 months infection across inoculation treatments. A one-way ANOVA was used to test for overall differences among treatments, followed by Tukey’s HSD post hoc test to identify significant pairwise comparisons (*p* < 0.05).

| **Comparison** | **diff** | **lwr** | **upr** | **p adj** |
| --- | --- | --- | --- | --- |
| W16.3_GacAΔ-W15.13_WT | -1.02229 | -1.96110 | -0.08348 | 0.02355 |
| W16.3_GacSΔ-W15.13_WT | -0.84961 | -1.80394 | 0.10472 | 0.11453 |
| W16.3_WT-W15.13_WT | 0.38656 | -0.56777 | 1.34089 | 0.88548 |
| W23.13_WT-W15.13_WT | 0.01502 | -0.93931 | 0.96935 | 1.00000 |
| WT:GacA_1:2-W15.13_WT | 0.54219 | -0.39662 | 1.48100 | 0.59298 |
| WT:GacA_2:1-W15.13_WT | 0.46824 | -0.47057 | 1.40705 | 0.74404 |
| W16.3_GacSΔ-W16.3_GacAΔ | 0.17268 | -0.78165 | 1.12701 | 0.99807 |
| W16.3_WT-W16.3_GacAΔ | 1.40886 | 0.45453 | 2.36319 | 0.00044 |
| W23.13_WT-W16.3_GacAΔ | 1.03731 | 0.08298 | 1.99164 | 0.02397 |
| WT:GacA_1:2-W16.3_GacAΔ | 1.56448 | 0.62567 | 2.50329 | 0.00005 |
| WT:GacA_2:1-W16.3_GacAΔ | 1.49054 | 0.55173 | 2.42935 | 0.00012 |
| W16.3_WT-W16.3_GacSΔ | 1.23618 | 0.26658 | 2.20578 | 0.00398 |
| W23.13_WT-W16.3_GacSΔ | 0.86463 | -0.10497 | 1.83423 | 0.11335 |
| WT:GacA_1:2-W16.3_GacSΔ | 1.39180 | 0.43747 | 2.34613 | 0.00055 |
| WT:GacA_2:1-W16.3_GacSΔ | 1.31786 | 0.36353 | 2.27219 | 0.00129 |
| W23.13_WT-W16.3_WT | -0.37154 | -1.34114 | 0.59806 | 0.90981 |
| WT:GacA_1:2-W16.3_WT | 0.15563 | -0.79870 | 1.10995 | 0.99893 |
| WT:GacA_2:1-W16.3_WT | 0.08168 | -0.87265 | 1.03601 | 0.99997 |
| WT:GacA_1:2-W23.13_WT | 0.52717 | -0.42716 | 1.48150 | 0.64286 |
| WT:GacA_2:1-W23.13_WT | 0.45322 | -0.50110 | 1.40755 | 0.78515 |
| WT:GacA_2:1-WT:GacA_1:2 | -0.07394 | -1.01276 | 0.86487 | 0.99998 |

Table S11 Significant pairwise comparisons of ash sapling symptom scores between treatments. Symptoms were scored on a scale of 0 (no necrosis, callused), 1 (blackened lesion edges), 2 (bark tissue necrosis), 3 (erumpent anker formation). Treatments were assessed for differences using a Kruskal-Wallis test, with pairwise comparisons assessed using a Dunn’s post-hoc test.

| **Comparison** | **Z** | **P unadjusted** | **P adjusted** |
| --- | --- | --- | --- |
| W1513 WT - W163 *gacS_W1513_* | 3.075156 | 0.002104 | 0.006546 |
| W163 *gacS_W1513_* - W163 WT | -2.61591 | 0.008899 | 0.022652 |
| W163 *gacS_W1513_* - W2313 WT | -3.29843 | 9.72E-04 | 0.003403 |
| W163 *gacA_W2313_* - WT W163 2:1 | -2.35458 | 0.018544 | 0.039941 |
| W163 *gacS_W1513_* - WT W163 2:1 | -4.3514 | 1.35E-05 | 5.41E-05 |
| W163 *gacS_W1513_* - WT W163 1:2 | -2.35458 | 0.018544 | 0.043269 |

Table S12 Chi-squared test to assess whether the proportion of motile to non-motile strains is significantly different to the expected ratio at time 0. Swarming motility was assessed for 56 colonies from four inoculation points from a single tree per treatment. Non-swarming is denoted NS.

| **Mixture** | **Obs. swarm** | **Obs, NS** | **Exp swarm (%)** | **Exp swarm** | **Exp NS** | **Swarm (O-E)^2^/E** | **NS (O-E)^2^/E** | **Sum** | **df** | **χ^2^** |
| --- | --- | --- | --- | --- | --- | --- | --- | --- | --- | --- |
| WT:GacAΔ 1:2 | 21 | 35 | 0.33 | 18.48 | 37.52 | 0.343636 | 0.169254 | 35.49394 | 4 | 3.68E-07 |
| WT:GacAΔ 1:2 | 32 | 23 | 0.33 | 18.15 | 36.85 | 10.56873 | 5.205495 |  |  |  |
| WT:GacAΔ 1:2 | 26 | 30 | 0.33 | 18.48 | 37.52 | 3.060087 | 1.507207 |  |  |  |
| WT:GacAΔ 1:2 | 28 | 28 | 0.33 | 18.48 | 37.52 | 4.904242 | 2.415522 |  |  |  |
| WT:GacAΔ 1:2 | 28 | 28 | 0.33 | 18.48 | 37.52 | 4.904242 | 2.415522 |  |  |  |
| WT:GacAΔ 2:1 | 50 | 6 | 0.66 | 36.96 | 19.04 | 4.600693 | 8.930756 | 45.27795 | 4 | 3.48E-09 |

Table S13 Chi-squared test to assess whether the proportion of motile to non-motile strains is significantly different to the expected ratio, 3 months post inoculation. Swarming motility was assessed for 56 colonies from two inoculation points from four trees, per treatment. Non-swarming is denoted NS.

| **Mixture** | **Obs. swarm** | **Obs, NS** | **Exp swarm (%)** | **Exp swarm** | **Exp NS** | **Swarm (O-E)^2^/E** | **NS (O-E)^2^/E** | **Sum** | **df** | **χ^2^** |
| --- | --- | --- | --- | --- | --- | --- | --- | --- | --- | --- |
| WT:GacA_1:2 | 35 | 21 | 0.33 | 18.48 | 37.52 | 14.76788 | 7.273731 | 356.6 | 7 | 5E-73 |
| WT:GacA_1:2 | 40 | 16 | 0.33 | 18.48 | 37.52 | 25.06009 | 12.34303 |  |  |  |
| WT:GacA_1:2 | 41 | 15 | 0.33 | 18.48 | 37.52 | 27.4432 | 13.5168 |  |  |  |
| WT:GacA_1:2 | 38 | 18 | 0.33 | 18.48 | 37.52 | 20.61853 | 10.15539 |  |  |  |
| WT:GacA_1:2 | 53 | 3 | 0.33 | 18.48 | 37.52 | 64.48216 | 31.75987 |  |  |  |
| WT:GacA_1:2 | 39 | 17 | 0.33 | 18.48 | 37.52 | 22.78519 | 11.22256 |  |  |  |
| WT:GacA_1:2 | 48 | 8 | 0.33 | 18.48 | 37.52 | 47.15532 | 23.22576 |  |  |  |
| WT:GacA_1:2 | 36 | 20 | 0.33 | 18.48 | 37.52 | 16.60987 | 8.180981 |  |  |  |
| WT:GacA_2:1 | 51 | 5 | 0.66 | 36.96 | 19.04 | 5.333377 | 10.35303 | 32.54 | 7 | 3E-05 |
| WT:GacA_2:1 | 47 | 9 | 0.66 | 36.96 | 19.04 | 2.727316 | 5.294202 |  |  |  |
| WT:GacA_2:1 | 42 | 14 | 0.66 | 36.96 | 19.04 | 0.687273 | 1.334118 |  |  |  |
| WT:GacA_2:1 | 41 | 15 | 0.66 | 36.96 | 19.04 | 0.441602 | 0.857227 |  |  |  |
| WT:GacA_2:1 | 40 | 16 | 0.66 | 36.96 | 19.04 | 0.250043 | 0.485378 |  |  |  |
| WT:GacA_2:1 | 42 | 14 | 0.66 | 36.96 | 19.04 | 0.687273 | 1.334118 |  |  |  |
| WT:GacA_2:1 | 42 | 14 | 0.66 | 36.96 | 19.04 | 0.687273 | 1.334118 |  |  |  |
| WT:GacA_2:1 | 40 | 16 | 0.66 | 36.96 | 19.04 | 0.250043 | 0.485378 |  |  |  |
